# A framework for testing structural hypotheses of protein dynamics against experimental HDX-MS data

**DOI:** 10.64898/2026.03.02.709114

**Authors:** Alexander I.H. Siddiqui, Rachael Skyner, Maria Musgaard, Srinath Krishnamurthy, Charlotte M. Deane, Oliver M. Crook

## Abstract

Protein dynamics determine biological function, yet extracting structural ensembles from Hydrogen–Deuterium Exchange Mass Spectrometry (HDX-MS) remains a challenging inverse problem. Current ensemble-fitting approaches often achieve good agreement with uptake curves but lack rigorous validation and uncertainty quantification, limiting structural confidence. We propose ValDX, a validation framework for quantitative integration of HDX-MS data with structural ensembles. ValDX combines overlap-aware data splitting, replicate-based uncertainty estimation, and uptake-independent “Work Done” metrics that quantify how much an ensemble must be modified to match experiment. Across 22 ensembles spanning six proteins (58–474 residues), we show that conventional error metrics fail to distinguish structurally representative ensembles from incorrect ones, whereas Work Done metrics robustly discriminate global and local conformational quality. We further demonstrate that clustering yields compact, interpretable ensembles with minimal loss of accuracy, and that staged optimisation enables reliable fitting of both ensemble weights and forward-model parameters without requiring a reference structure. Together, this framework establishes HDX-MS ensemble integration as a quantitative structural hypothesis-testing problem, enabling inference of protein dynamics from HDX-MS data.

## 1 Introduction

Understanding protein dynamics is crucial to understanding biological function, and therefore disease. HDX-MS is a biophysical technique that measures conformational dynamics in solution.[1–4] Orthogonal to other biophysical techniques, such as X-Ray Crystallography (XRC or X-Ray); Cryogenic Electron-Microscopy (CryoEM); or Nuclear Magnetic Resonance (NMR), HDX-MS probes without size limitations and is able to uniquely unravel protein dynamics under diverse conditions.[4, 5] This flexibility combined with increasing throughput means HDX-MS offers unparalleled insights for understanding challenging targets, such as G-Protein Coupled Receptors (GPCRs),[6, 7] antibodies,[8–10] and Intrinsically Disordered Proteins (IDPs).[11, 12]

While HDX-MS is powerful for detecting conformational changes, the technique presents a fundamental interpretive challenge. HDX-MS reports deuterium uptake at the peptide level, not the residue level.[13, 14] An HDX-MS experiment produces uptake curves for overlapping peptide fragments, each representing an average over all exchangeable amide positions within that peptide. This means that a single uptake curve can arise from many different underlying structural scenarios. For example, a peptide region that transiently unfolds a helix may appear identical to one that samples multiple partially protected states. Without additional structural context, determining *which* conformational states give rise to an observed uptake pattern is fundamentally ambiguous.

To address this ambiguity, researchers increasingly turn to molecular simulations for structural hypotheses.[14–19] Given a collection of candidate protein conformations from simulation, one can predict what HDX-MS uptake each conformation would produce, then adjust how much each conformation contributes to find the combination that best matches experiment. This process of adjusting structural contributions, referred to as ensemble reweighting, allows proposed conformational states to be tested against experimental data.[13–20] When successful, ensemble reweighting can reveal which conformations are consistent with experiment and which are not, providing pseudo-structural models of protein behaviour in solution.[13, 15, 20]

However, current implementations of ensemble reweighting face a critical validation problem. In typical scenarios, good agreement does not always lead to correct solutions. The same experimental data can often be fit equally well by dramatically different structural ensembles. This occurs because the structure-to-uptake models used for prediction, while based on physical principles of hydrogen bonding and solvent exposure,[21, 22] are approximate and introduce systematic errors that accumulate at the peptide level.[23, 24] Without a means to distinguish genuinely correct solutions from fortuitously good fits, ensemble reweighting cannot reliably discriminate between competing structural hypotheses.

A second, related problem is overfitting. Because HDX-MS peptides overlap extensively, fitting all peptides simultaneously can mask poor model performance. Errors in one region can be compensated by implausible structural distortions elsewhere. Standard practice reports only training error, how well the model fits the data it was optimised against, but this metric is demonstrably unreliable. An ensemble containing entirely incorrect structures can achieve low training error simply by exploiting the flexibility of the fitting procedure. What is needed instead is a framework that tests whether a model generalises; ie., this is to ask whether a fitted ensemble model can predict uptake for peptides it was not trained on and what structural adjustments were necessary to achieve the fit.

In this work we present ValDX, a validation framework designed to solve these proble,s. ValDX provides the user with two key capabilities. First, it implements rigorous data splitting to evaluate whether ensemble models generalise beyond their training data. Specifically, because HDX-MS peptides overlap, naive random splitting causes information to leak between training and test sets; ValDX employs redundancy-aware and spatially-informed splitting strategies to ensure that validation error reflects true predictive performance. Second, ValDX introduces ‘Work Done’ metrics that quantify the cost of fitting, how much the optimizer must distort the original structural ensemble to match experiment. When fitting requires large distortions, this signals that the starting ensemble may lack the correct conformational states; when small adjustments suffice, this suggests the ensemble was already representative of solution behaviour.

Central to this workflow (Figure 1) is the recognition that standard error metrics alone cannot validate structural conclusions. Data splitting is a well-established approach in machine learning to test whether a model captures genuine patterns rather than noise. By fitting on a subset of data (training set) and evaluating on the remainder (validation set), one can assess generalisability. However, applying data splitting to HDX-MS is not straightforward. Because peptides overlap, and the same residues can appear in multiple peptides, a random split will include peptides in the validation set whose uptake is predictable from training peptides covering similar regions. This information leakage inflates apparent validation performance. ValDX addresses this through multiple split types: random splits (as a baseline), sequential splits (ensuring validation peptides do not overlap with training), redundancy-aware splits (explicitly accounting for shared residue coverage), and spatial splits (grouping peptides by structural region). Exact algorithmic details can be found in the Assessing Global vs. Local Scales through Peptide Splitting section in Materials and Methods.

**Figure 1:**
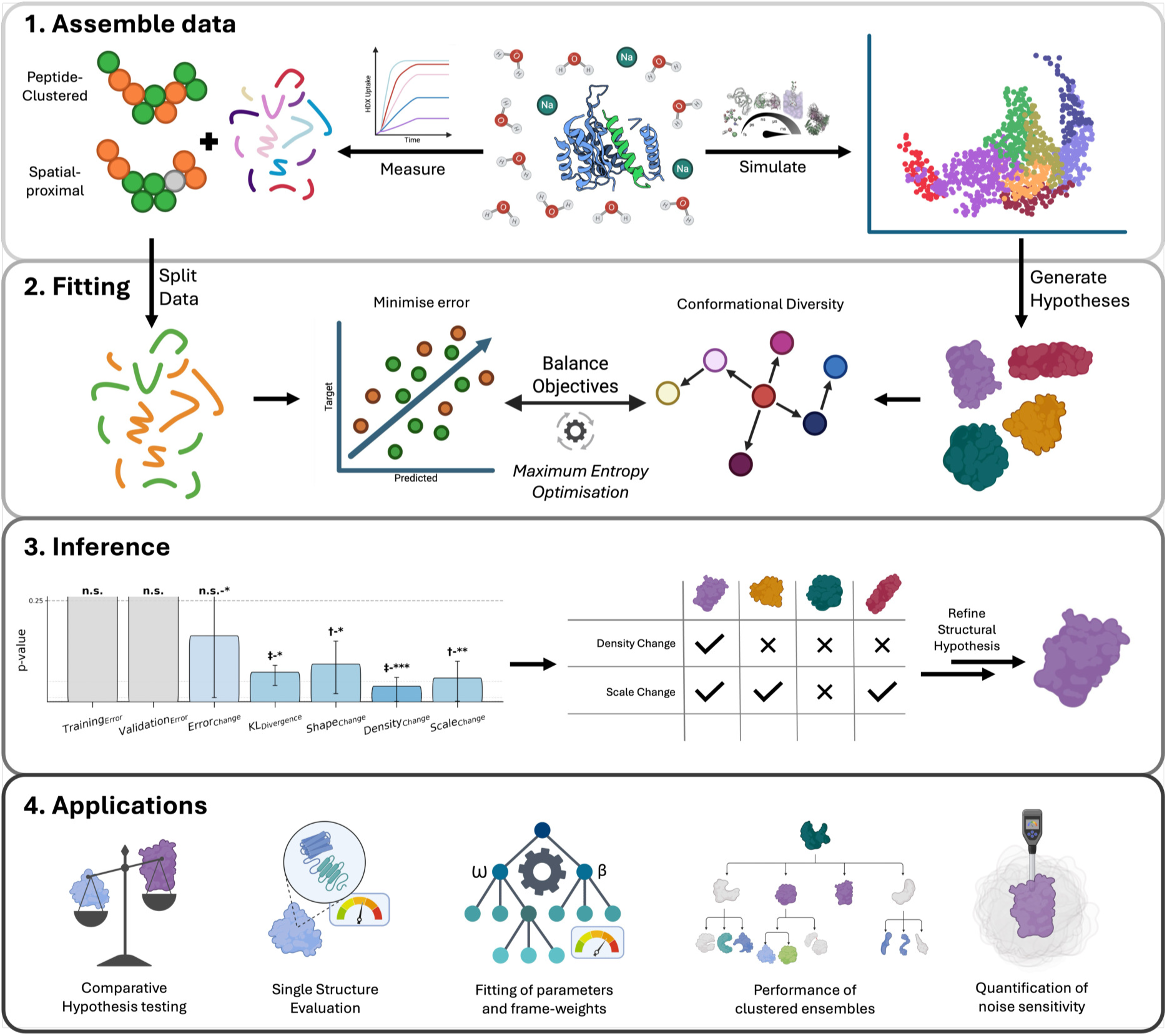
ValDX workflow for integrating HDX-MS data with biomolecular simulations. The workflow consists of three main phases: (1) **Assemble data** - HDX-MS peptide data (left) is split using multiple strategies including Random Split, Sequential Split, Redundant Aware Split, and Spatial Split to create training and validation datasets while preventing data leakage. (2) **Fitting** - Structural hypotheses are generated from principal component analysis (right), from which structural features are computed. The training model is optimized across multiple data splits and replicates using Maximum Entropy Reweighting to balance fitting strength with Work Done. Validation error is computed on held-out data (left). (3) **Inference** - Structural changes during optimization are extrapolated to calculate Work Done during optimization which are more sensitive than error (left). Ensemble hypotheses are selected by work done across data-splits and metrics (right) . (4) **Applications** - The framework enables comparative hypothesis testing, assessment of individual structures, fitting of parameters and weights, performance evaluation of clustered ensembles, and quantification of noise sensitivity, providing a statistically rigorous framework for advancing HDX-MS from qualitative to quantitative structural analysis.

The second cornerstone of ValDX is the quantification of optimisation cost. When ensemble reweighting adjusts structural contributions to fit experimental data, it performs ‘work’ on the ensemble-average distribution. We define three complementary Work Done metrics: Work_shape_ measures how much the protection factors change (capturing whether the optimizer is skewing the dynamics), while Work_scale_ measures the magnitude or scale of protection factor changes. Work_density_ quantifies the extent of protection factor reorganisation, this measures how conformational states are redistributed at the ensemble-average level.

Critically, these metrics are computed independently of HDX-MS uptake predictions. This independence is essential as it eliminates systematic, confounding errors from residue-peptide mismatches. As a result, precise statements can be made about structural hypotheses. For example, if a fit requires dramatic redistribution of weights or model parameters (high Work Done) yet achieves good experimental agreement, this indicates the starting ensemble was structurally deficient. Conversely, ensembles that fit well with minimal weight/parameter adjustment are more likely to represent genuine solution conformations. Complete derivations for these metrics are provided in Derivation of Work Done Metrics within the Additional Experiment Plots.

We systematically apply ValDX across proteins spanning diverse conformational behaviours to demonstrate its broad applicability. These systems range from 58 to 474 residues and include rigid proteins with well-defined folds, flexible multi-domain systems, and intrinsically disordered proteins. Details of each protein and associated HDX-MS data are provided in HDX Data. For each system, we generate standardised structural ensembles using hierarchical protocols (described in Ensemble Generation Experiments in Materials and Methods) to ensure findings are comparable across systems and ensemble generation methods. Pairwise distance comparisons for all ensembles are provided in Ensemble Analyses within the Additional Experiment Plots.

Through these experiments, we establish that conventional training error metrics fail to identify correct structural ensembles even under ideal conditions, while the combination of validation error and Work Done metrics provides robust discrimination. We show that ensemble reweighting is sensitive to the presence of incorrect structures, demonstrate optimal fitting protocols, and determine that meaningful ensemble representation can often be achieved with as few as 10–13 carefully selected structures. Together, these findings provide a rigorous foundation for advancing HDX-MS from a qualitative probe of conformational change to a quantitative structural technique capable of distinguishing between competing mechanistic hypotheses.

## 2 Results

### 2.1 Conceptual Framework for Ensemble Validation

Integrating structural ensembles with HDX-MS data presents a fundamental validation challenge: experimental agreement alone cannot distinguish an ensemble that genuinely represents solution-state dynamics from one that has been forced into agreement through overfitting. This ambiguity arises because the optimisation process possesses sufficient degrees of freedom to accommodate incorrect structural hypotheses, producing low prediction errors that mask underlying deficiencies. The ValDX framework addresses this limitation by quantifying not only predictive accuracy but also the magnitude of ensemble modification required to achieve that accuracy, a quantity termed “Work Done” that provides essential complementary information about structural representativeness.

#### Optimisation Parameters and Their Physical Interpretation

The fitting procedure can modify two distinct sets of parameters, each with clear physical meaning. First, **ensemble population weights** control the fractional contribution of each conformational state to the predicted HDX signal. Adjusting these weights corresponds physically to rebalancing how frequently the protein visits each conformation in solution; a representative ensemble requires minimal population redistribution. Second, the **Best-Vendruscolo model parameters** (*β*_C_ for heavy-atom contacts, *β*_H_ for hydrogen bonds) govern how structural features translate into residue-level protection factors. These parameters primarily compensate for systematic differences between experimental conditions (pH, temperature, quench efficiency) and the conditions under which the model was originally calibrated, manifesting as uniform shifts in predicted exchange rates.

#### Work Done Metrics: Statistical Quantities with Structural Meaning

The Work Done frame-work employs metrics derived from information-theoretic principles within the Maximum Entropy formalism, expressed in energy units (kJ/mol) to facilitate physical interpretation. We emphasise that these quantities represent statistical distances between probability distributions rather than thermodynamic state functions, though the mathematical analogy to thermodynamic potentials provides useful physical intuition. Smaller values consistently indicate more representative ensembles that required less modification to match experimental observations. Uniform energy scaling facilitates comparison of Work Done metrics across systems.

The metric Work_shape_ (denoted Δ*H*_opt_ by analogy to enthalpy) quantifies changes in the *relative pattern* of protection factors across residues. Specifically, mean-centred deviations in the protection factor distribution. Large values indicate that optimisation substantially reorganised which protein regions appear protected versus solvent-exposed, typically signifying incorrect local structure, erroneous hydrogen-bonding networks, or missing conformational states that would explain regional exchange behaviour.

The metric Work_scale_ (Δ*H*_abs_) captures changes in the *overall magnitude* of predicted protection, reflecting uniform scaling of exchange rates across the protein. Elevated values suggest systematic mismatch between experimental conditions and model calibration rather than structural deficiencies, and typically indicate that model parameter recalibration should be considered before rejecting the ensemble hypothesis.

The metric Work_density_ (denoted −*T* Δ*S*_opt_ by analogy to entropic contributions) quantifies the reorganisation of ensemble-average protection factors during optimisation. Large values indicate that the optimiser substantially shifted the distribution of protection factors, suggesting that the initial conformational sampling was skewed relative to the true solution-state populations or that important conformational states were inadequately sampled.

The total Work_opt_ (Δ*G*_opt_) combines shape and density contributions, providing the primary summary metric for ensemble comparison. This quantity represents the total information-theoretic cost of transforming the initial ensemble hypothesis into one that matches experimental observations.

#### Diagnostic Interpretation of Fitting Outcomes

Successful ensemble validation manifests as low prediction error on withheld peptides (MSE_validation_) accompanied by small Work_opt_, indicating that the ensemble matched experimental data without substantial modification. Reproducibility across replicate optimisations and consistency between data partitioning strategies provide additional confidence in the result.

Several characteristic failure modes emerge from the interplay between error and Work metrics. When low validation error accompanies high Work Done, the ensemble achieved experimental agreement through excessive modification. This is a signature of overfitting indicating that the structural hypothesis should be considered cautiously, despite apparent predictive success. Caution applied is proportionate to the spread of metrics across replicate optimisations. Conversely, simultaneous elevation of both error and Work signals a fundamentally inadequate ensemble that could not match experimental observations even with substantial modification, warranting exploration of alternative structural generation approaches. Isolated elevation of Work_scale_ with other metrics remaining low suggests experimental condition mismatch rather than structural problems, directing attention toward model parameter calibration. Elevated Work_density_ in isolation indicates that the ensemble-average protection factor distribution was substantially reorganised, pointing toward inadequate sampling or missing conformational states. Finally, high Work_shape_ alone suggests local structural inaccuracies requiring examination of specific protein regions for misfolding or alternative conformations.

#### Data Splitting Strategies

Because HDX-MS peptides exhibit substantial sequence overlap, standard random splitting risks information leakage wherein validation peptides share exchange information with training peptides. ValDX employs two complementary, overlap-aware splitting strategies designed to probe ensemble quality at different structural scales. **Non-Redundant splitting** clusters peptides by sequence position before partitioning, ensuring validation peptides represent genuinely withheld sequence regions and testing whether the ensemble captures global protein behaviour. **Spatial splitting** withholds peptides covering contiguous three-dimensional regions, testing whether the ensemble accurately represents local or substructural dynamics.

On top of within-split reproducibility, between-split consistency or agreement provides a powerful diagnostic tool. Orthogonal to confidence, agreement is useful to identify true structural inadequacies from artifacts of peptide selection, as different splitting strategies probe distinct structural aspects. Good metric agreement (small distance between averages) between splitting strategies indicates reliable structural representation; discrepancies reveal scale-dependent ensemble deficiencies that inform subsequent structural refinement.

### 2.2 Ensemble integration success is insufficiently explained by training error

Traditional assessments of ensemble-data integration rely on minimising prediction errors, or how well computed HDX-uptake curves match experimental measurements. We investigated whether this error-centric paradigm adequately captures integration success, particularly when multiple structural hypotheses yield similar experimental agreement. To address this question with a known baseline or ground truth, we developed the Iso-Validation benchmark using TeaA, a domain of the membrane transporter complex TeaABC, that undergoes large-scale conformational transitions between open and closed states during substrate transport (Figure 2).[13] These conformational changes, captured in published Molecular Dynamics (MD) trajectories, provide structurally distinct functional states, an ideal system for testing whether current metrics can distinguish correct from incorrect ensemble reweighting.[13] The experimental protocol is detailed in Case Study Protocols within Materials and Methods.

To establish controlled conditions enabling direct accuracy measurement, we generated synthetic HDX-uptake data with known conformational populations. We first clustered the MD ensemble by Root Mean Squared Deviation (RMSD) using a 1.0 Å threshold relative to predefined open and closed reference states (Figure 2A), then artificially modified the population ratio from 5:95 to 40:60 (open:closed) and predicted HDX-uptake curves from this reweighted ensemble. This design guaranteed that the target populations could be recovered through ensemble reweighting alone, enabling us to assess metric performance against ground truth. We created two structural hypotheses: ISO-Trimodal (gold), the parent unclustered ensemble including intermediate conformations, and ISO-Bimodal (indigo), the clustered ensemble representing only the two functional end-states (Figure 2B-C). Critically, ISO-Bimodal contains exclusively ground-truth structures present in the synthetic data, while ISO-Trimodal includes intermediate conformations absent from the target. We performed ensemble optimisation across a range of *γ*_HDXer_ values, a regularisation parameter that controls the balance between fitting the experimental data (high *γ*) versus maintaining the prior ensemble populations (low *γ*), following standard practice in Maximum Entropy reweighting.[13, 17, 25–27] The experimental design employed four split-types with three replicates each: Random (fuchsia) and Sequential (black), representing standard approaches, alongside our Non-Redundant (green) and Spatial (grey) methods that account for peptide overlap. Training error (MSE_Training_) was computed on the fitting peptides, as detailed in HDXer Loss Function. In Figure 2D-E, we plot MSE_Training_ against Apparent Work_HDXer_, with increasing point size indicating ascending *γ*_HDXer_ values.

**Figure 2:**
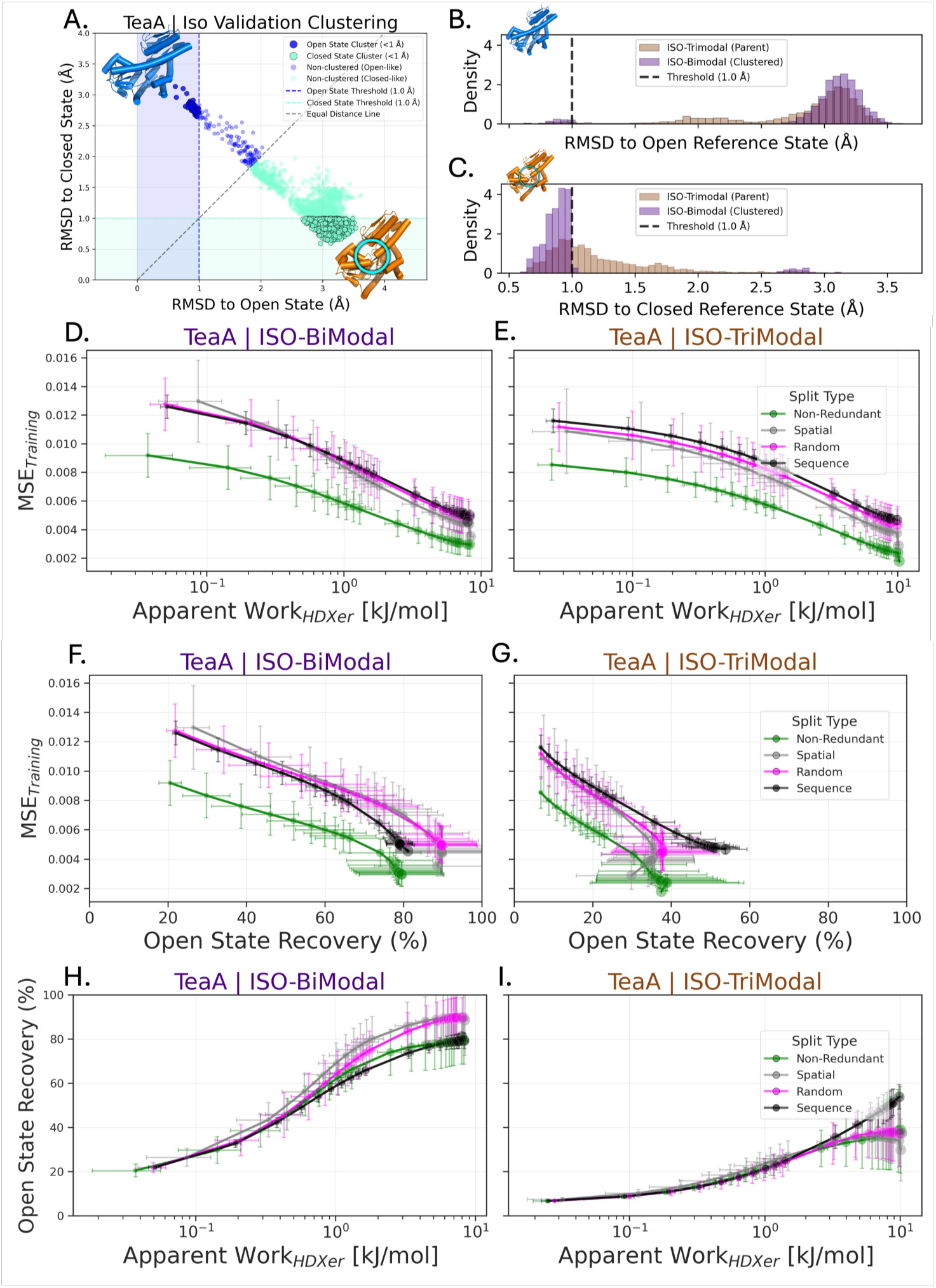
Training error fails to distinguish ensembles with correct versus incorrect conformational populations in Iso-Validation benchmark experiments. **(A)** TeaA domain ensemble clustering by RMSD (*<*1.0 Å) to open (blue) and closed (cyan/orange) reference states, yielding two test ensembles. RMSD density distributions for **(B)** open and **(C)** closed states comparing ISO-Trimodal (unclustered parent, gold) and ISO-Bimodal (clustered, indigo) ensembles. Both ensembles fitted to synthetic HDX data with known 40:60 open:closed population. Training error (MSE_Training_) versus Apparent Work (Work_HDXer_) for **(D)** ISO-Bimodal and **(E)** ISO-Trimodal across four split-types (Non-Redundant, green; Spatial, grey; Random, fuchsia; Sequential, black); point size indicates regularisation strength (*γ*_HDXer_). Ground-truth Open State Recovery (%, normalised to target 40%) versus MSE_Training_ for **(F)** ISO-Bimodal and **(G)** ISO-Trimodal, demonstrating training error cannot predict population recovery for ensembles containing incorrect intermediate structures. Open State Recovery versus Apparent Work_HDXer_ for **(H)** ISO-Bimodal and **(I)** ISO-Trimodal, showing Work Done metrics consistently correlate with ground-truth performance independent of split-type choice. Results establish that matching experimental uptake curves is insufficient to verify conformational population accuracy. Work Done metrics provide essential complementary information about ensemble quality.

Training error failed to distinguish between ensembles despite their fundamentally different structural content, important given that ISO-Bimodal contains only true conformational states while ISO-Trimodal includes irrelevant intermediates. Both ensembles achieved similarly low MSE_Training_ values, demonstrating that uptake agreement alone cannot verify population recovery. While this inseparability might initially suggest insufficient model sensitivity, closer examination revealed a more subtle pattern. The variation in training error between different split-types of the *same* ensemble exceeded the variation between *different* ensembles using identical splits. Since split-replicates maintain consistent peptide distributions across both ensemble experiments, this pattern indicates that optimisation is highly dependent on which specific peptides are used for training—peptide sampling artifacts dominate over genuine structural differences.

We tested training error directly against ground truth by comparing MSE_Training_ to Open State Recovery%, our uptake-independent metric normalised to the target population of 0.4 (40%). Because our synthetic data derived from structures with known population ratios, the recovered fraction of open-state structures provided a direct, model-independent measure of integration success. The comparison revealed an informative asymmetry. For ISO-Bimodal (Figure 2F), lower training error did correlate with improved ground-truth recovery, reaching a performance plateau between 80–90% across all split-types, the ensemble containing only correct structures could be successfully reweighted when guided by uptake error alone. However, for ISO-Trimodal (Figure 2G), the relationship collapsed. Open State Recovery struggled to exceed 40% for three of four split-types despite achieving low training error comparable to ISO-Bimodal. The presence of intermediate structures, plausible conformations absent from the true ensemble, prevented accurate population recovery even when uptake curves matched experimental targets. Since our optimisation employed the self-consistent Best-Vendruscolo (BV) protection factor model (Equation BV Model), we could exclude peptide-level uptake prediction limitations as the source of this failure. These results establish that predicted-experimental uptake error is insufficient for assessing whether ensemble populations reflect true conformational distributions.

Given the limitations of error-based metrics, we investigated whether Work Done metrics, which quantify ensemble modification during optimisation, might better assess integration quality. We compared ground-truth Open State Recovery% against Apparent Work_HDXer_, the Work Done computed by HDXer on training data. The relationship proved substantially more informative: for both ensembles, increasing Apparent Work_HDXer_ consistently indicated improved Open State Recovery% (Figure 2H-I). Importantly, Apparent Work_HDXer_ distributions showed greater consistency across split-types than MSE_Training_ distributions, as detailed in Figure S9. This stability is critical because biomolecular simulations vary in quality and representativeness; a metric that fluctuates based on arbitrary peptide selection cannot reliably guide ensemble evaluation.

To quantify metric performance systematically, we performed multiple linear regression to assess the independent predictive utility of MSE_Training_ and Apparent Work_HDXer_ for ground-truth Open State Recovery% (complete methodology in Metric Quality Quantification). By fitting both metrics simultaneously in a standardised model (Equation S1), we isolated their distinct contributions to explaining population recovery. The results, shown in Figure S10.A, revealed Work Done as the dominant predictor. For ISO-Bimodal, Apparent Work_HDXer_ showed strong positive association (standardised coefficient *θ* = 9.62, partial *R*^2^ = 0.066) while training error contributed weakly and negatively (*θ* = −6.37, partial *R*^2^ = 0.040). This trend intensified for ISO-Trimodal, Apparent Work_HDXer_ maintained robust predictive power (*θ* = 10.62, partial *R*^2^ = 0.129) while training error became essentially uninformative (*θ* = −0.13, partial *R*^2^ ≈ 0.000). These standardised coefficients demonstrate that Apparent Work_HDXer_ provides substantially greater insight than MSE_Training_, particularly when ensembles contain structurally plausible but functionally incorrect conformations.

Beyond predictive value, we assessed metric robustness using variance decomposition analysis. We computed stability indices quantifying the proportion of total variance arising from within-split rather than between-split variation (Equation S5). Apparent Work_HDXer_ proved remarkably stable across split-types for both ensembles (ISO-Bimodal: 0.809; ISO-Trimodal: 0.977), indicating minimal sensitivity to peptide selection (Figure S10.D). In stark contrast, MSE_Training_ exhibited extremely low stability (ISO-Bimodal: 0.079; ISO-Trimodal: 0.070), confirming strong dependence on which peptides comprise the training set. Coefficient of variation analysis reinforced this conclusion (Figure S10.E): Apparent Work_HDXer_ showed low variability across splits (CV = 0.018–0.049) while MSE_Training_ exhibited approximately four-fold higher variability (CV = 0.171–0.191). ANOVA analysis (Figure S10.B-C) confirmed this pattern quantitatively. Split-type choice explained substantial variance in MSE_Training_ (effect size *η*^2^ = 0.132–0.147; *F* = 11.7–13.2, *p <* 0.001) but had negligible impact on Apparent Work_HDXer_ (*η*^2^ = 0.000–0.003; *F* = 0.0–0.2, *p >* 0.05).

This robustness is particularly significant for practical applications where simulation quality cannot be guaranteed *a priori*. The contrasting behaviours demonstrate that MSE_Training_ primarily reflects peptide sampling artifacts, arbitrary consequences of which peptides were selected for fitting, rather than genuine ensemble quality differences. In contrast, Apparent Work_HDXer_ captures structural representativeness independent of data partitioning strategy. Taken together, these analyses establish Apparent Work_HDXer_ as a superior metric for assessing ensemble optimisation, combining greater predictive power for ground-truth recovery with remarkable stability across experimental design choices.

These results motivated development of the complete ValDX workflow. While Iso-Validation enabled direct ground-truth comparison through synthetic data, realistic integration scenarios lack such reference points. In practice, validation error (MSE_Validation_), error computed on peptides withheld during optimisation, becomes our only available proxy for assessing generalisation to unseen data. However, when applied to experimental peptide datasets, Random and Sequential splits provided no additional discriminatory power beyond training error. Consequently, all subsequent experiments employ exclusively Non-Redundant and Spatial split-types, which account for peptide redundancy and provide complementary structural perspectives.

### 2.3 Work Done metrics reveal distinct ensemble representativeness at global and local structural scales

Having established that training error alone is insufficient to evaluate ensemble quality, we applied the ValDX framework to real experimental HDX-MS data to test whether Work Done metrics could distinguish between structurally similar yet biologically distinct ensemble proposals. We chose Bovine Pancreatic Trypsin Inhibitor (BPTI), a well-characterised small protein commonly used for method validation whose conformational dynamics have been extensively studied.[28–33] This protein’s rigid core and flexible terminal regions make it an ideal test case for evaluating whether our metrics can detect subtle conformational differences that arise from different ensemble generation approaches. This is a critical capability for assessing structural hypotheses in less well-studied systems.

We compared two ensemble generation methods that sample BPTI conformational space differently. The first, MD-1Start, represents conventional equilibrium molecular dynamics initiated from a single high-quality crystal structure (five independent 20 ns simulations, 100 ns total production time). The second, AF2-Filtered, leverages recent advances in deep learning-based structure prediction: we generated diverse conformations using AlphaFold2 with Multiple Sequence Alignment (MSA) subsampling via ColabFold,[34] then filtered the output to remove physically implausible geometries. This comparison directly tests whether statistical approaches, which sample global conformational changes through evolutionary variation rather than local physical dynamics, can capture relevant solution-state protein behaviour when compared to traditional simulation methods for BPTI. Both ensembles underwent identical fitting procedures: reweighting ensemble populations to align against experimental HDX-uptake without modifying structures themselves (RW-only), across three replicate data partitions for both Non-Redundant splits (testing global consistency over all peptides, accounting for overlap) and Spatial splits (testing local consistency by withholding spatially contiguous peptide clusters). Complete ensemble generation and fitting protocols are detailed in Reweighting-Only Protocol under Case Study Protocols in Materials and Methods, with AlphaFold2 procedures in AlphaFold Experiments.

The two ensembles exhibit meaningful structural differences despite sampling the same protein to an equivalent level of computational expense. To quantify these differences objectively, we computed the Wasserstein_1_ distance (*W*_1_) for pairwise *C_α_*distances across all residue pairs (Figure 3.A). *W*_1_ measures the minimum local structural transformation required to convert one conformational distribution of contacts into another, as detailed in Quantifying Ensemble Similarity and Conformational Coverage. The largest structural divergence appeared at BPTI’s flexible N- and C-termini (*W*_1_ = 0.5 Å) and, more significantly, at the disulphide bridge-containing outer loop of the hairpin turn. This hairpin region is particularly informative as localised conformational diversity here correlates with global conformational changes relevant to BPTI’s solution behaviour.[35, 36] While 20 ns per replicate (100 ns total) is clearly not sufficient to obtain conformational coverage in MD-1Start, even publically available 1 ms MD simulations show incomplete sampling of this region, relative to predictions from experimental studies.[37] Complete coverage of the *in vitro* disulphide torsion angles involves isomerisation. While molecular mechanics prohibit bond breaking, we address this by generating ensembles with multiple starting conformations or topologies (MD-10Start and MD-TFES).

**Figure 3:**
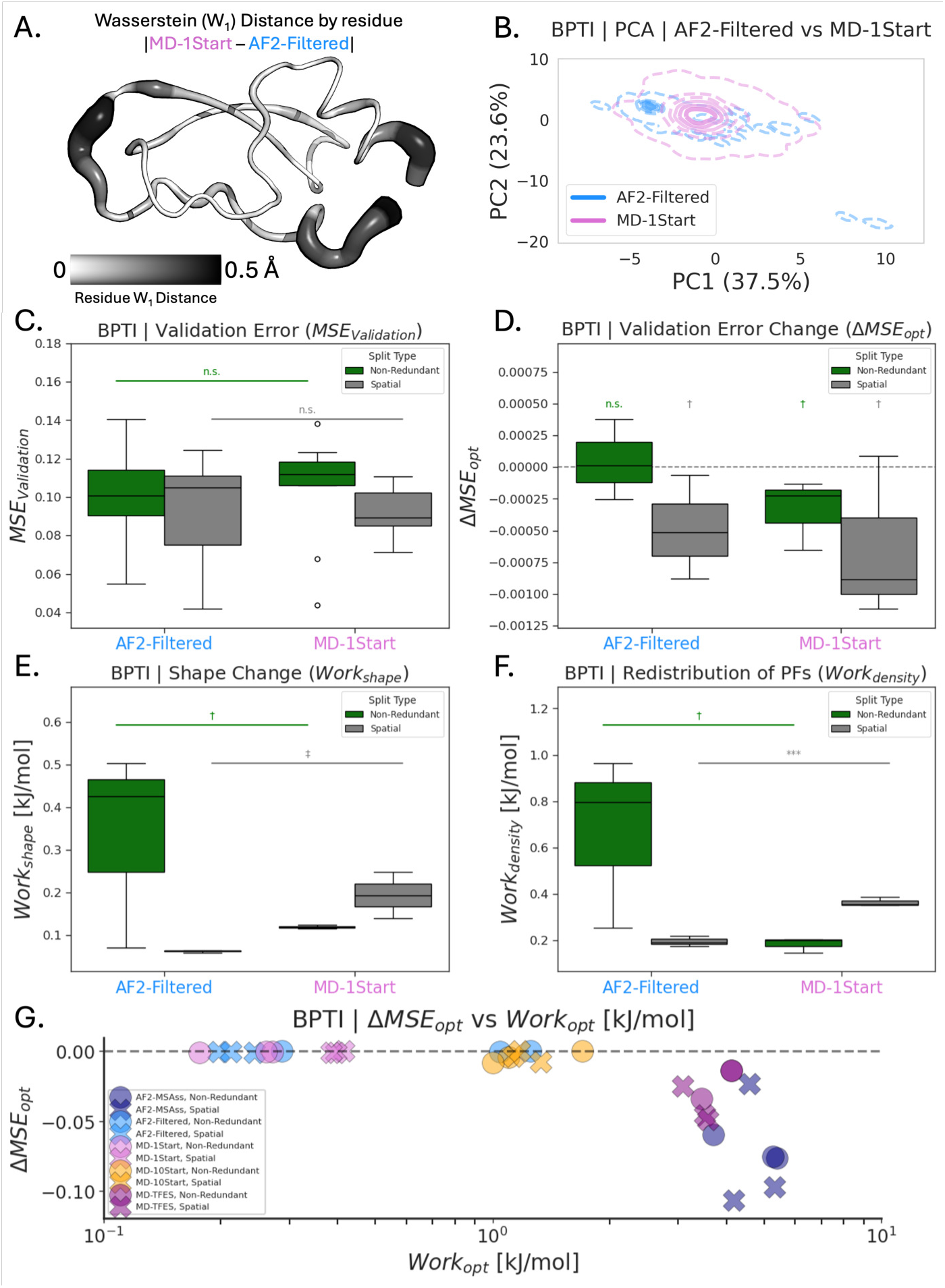
Work Done metrics distinguish ensemble representativeness at multiple structural scales in BPTI. AlphaFold2 MSA-subsampling captures a rare hairpin-opening state (blue) missed by conventional MD (pink). **(A)** Structural divergence quantified by Wasserstein distance (*W*_1_) between MD-1Start and AF2-Filtered ensembles, with largest differences (0.5 Å) at flexible termini and disulphide-containing outer loop. **(B)** Principal component analysis reveals AF2-Filtered samples a conformational superset of MD-1Start, including a minor basin corresponding to outer-hairpin opening. **(C)** Validation error (MSE_Validation_) fails to distinguish ensembles for either global (Non-Redundant, green) or local (Spatial, grey) data partitions despite structural differences. **(D)** Validation error change upon optimisation (ΔMSE_opt_) shows both ensembles require minimal adjustment, with statistically insignificant differences. **(E)** Energy-scaled prediction magnitude changes (Work_shape_) successfully separate ensembles, revealing MD-1Start better represents global structure (Non-Redundant) while AF2-Filtered better represents local structure (Spatial). **(F)** Protection factor reorganisation intensity (Work_density_) strengthens discrimination, with highly significant Spatial-split difference indicating fundamental distinctions in sub-structural conformational coverage. **(G)** Systematic comparison of ensemble generation approaches plotting validation error change versus total Work Done reveals MD-1Start and AF2-Filtered as most representative (smallest modifications required), clearly separated from enhanced-sampling MD-TFES and geometrically unfiltered AF2-MSAss. Work Done metrics provide quantitative assessment of conformational representativeness that operates simultaneously at global and local structural scales, enabling objective comparison of structural hypotheses.

Principal Component Analysis (PCA) revealed the global structural relationship between AF2-Filtered and MD-1Start: AF2-Filtered forms a superset of MD-1Start, sharing the primary conformational basin while additionally sampling a minor well corresponding to opening of the outer-hairpin turn. (Figure 3.B, details in Quantifying Ensemble Similarity and Conformational Coverage). These ensemble properties raise an important biological question: do rare, hairpin-opening states exist in solution, or does AlphaFold2’s sampling strategy introduce conformational heterogeneity absent from the true ensemble? Answering this question requires methods that go beyond structural comparison alone.

Traditional validation metrics failed to resolve this biological question. Despite their structural differences, neither Non-Redundant nor Spatial data partitions revealed significant differences in validation error (MSE_Validation_) between MD-1Start and AF2-Filtered (*p* = 0.765 to 0.975, Figure 3.C). We examined whether measuring the change in validation error before and after optimisation (ΔMSE_opt_) might provide better discrimination. If one ensemble requires less error reduction to fit the data, it should represent the solution state more faithfully. However, both ensembles benefited only modestly from reweighting, with MD-1Start showing slightly greater improvement that remained statistically insignificant (*p* = 0.17 to 0.824, Figure 3.D). These results confirm our Iso-Validation findings. When structure-uptake prediction models have inherent accuracy limitations, validation error lacks sufficient statistical power to distinguish between plausible ensemble hypotheses, even when we measure error reduction rather than absolute error magnitude.

Work Done metrics successfully distinguished the ensembles where validation error could not. We first measured Work_shape_, which quantifies the energy-scaled magnitude of changes in predicted protection factor distributions between the initial and optimised ensembles. Operationally, this metric captures how much the optimizer needed to shift the overall HDX-uptake predictions (faster or slower exchange) to match experimental data. Smaller values indicate the initial ensemble already predicted exchange rates close to experimental observations, suggesting more representative structural proposals. The results strongly supported this interpretation. Work_shape_ successfully separated the two ensembles with improved statistical significance compared to validation error (Non-Redundant: *p* = 0.249; Spatial: *p* = 0.0517, Figure 3.E). Notably, ensemble performance depended on the structural scale probed by each data partition: MD-1Start required less modification than AF2-Filtered when tested globally (Non-Redundant split), indicating better representation of BPTI’s overall conformational ensemble, but required more modification when tested locally (Spatial split), suggesting AF2-Filtered captures sub-structural flexibility more effectively.

To understand whether these differences reflected conformational density coverage or prediction magnitude shifts, we examined Work_density_, the information-theoretic entropy change quantifying the reorganisation of ensemble-average protection factors during optimisation. This metric revealed substantially stronger discrimination (Non-Redundant: *p* = 0.15; Spatial: *p* = 0.000678, Figure 3.F), with the highly significant Spatial split result demonstrating that the two ensembles differ fundamentally in how well they represent local conformational heterogeneity. Large Work_density_ values indicate the optimizer needed to substantially reorganise the distribution of protection factors, either concentrating density onto specific conformational modes or spreading it more broadly to match experimental exchange patterns.

The complementary information from both metrics allowed quantitative, multi-scale assessment of ensemble representativeness. Considering Work_shape_ and Work_density_ together, the data suggest that MD-1Start more faithfully represents BPTI’s global conformational ensemble (requiring minimal protection factor reorganisation and magnitude adjustment when fitting all peptides simultaneously), while AF2-Filtered provides superior coverage of sub-structural flexibility (requiring less conformational reorganisation when fitting spatially localised peptide clusters independently). The similarity between AF2-Filtered’s Spatial-split performance and MD-1Start’s Non-Redundant-split performance in terms of protection factor reorganisation suggests that AlphaFold2’s MSA-driven sampling captures aspects of local conformational heterogeneity reasonably well, even though it samples a broader conformational range globally. Notably, despite both ensembles showing a low median *W*_1_ distance of 0.25 Å for ensemble-average pairwise *C_α_*distances and only modest shifts in PCA projections, these Work Done analyses revealed functionally important differences invisible to global structural metrics. This demonstrates that conformational representativeness operates at multiple structural scales.

To place these findings in broader context, we compared MD-1Start and AF2-Filtered against a systematic panel of ensemble generation approaches by plotting validation error change (ΔMSE_opt_) against total Work Done (Work_opt_, Figure 3.G). The panel of ensembles, whose protocols are de-tailed in Materials and Methods, was designed not to represent optimised workflows but rather to systematically probe how different sampling strategies affect ensemble representativeness: varying initial structure quality, simulation time, and diversity enhancement methods. This allows direct comparison of shared and distinct properties across different systems without confounding protocol-specific optimisation choices. The comparison revealed MD-1Start and AF2-Filtered as our most representative BPTI ensembles, requiring the smallest modifications upon optimisation (particularly evident in Spatial splits, marked by crosses). In Non-Redundant splits (circles), AF2-Filtered’s total Work Done approached that of MD-10Start (equilibrium MD initiated from ten diverse conformational starting points), with overlapping confidence intervals suggesting comparable global representativeness. These three ensembles separated clearly from two substantially less representative proposals. These include: MD-TFES (an ensemble generated using Topology Frontier Expansion Sampling, an enhanced seeding algorithm designed to maximise structural diversity under equilibrium dynamics, see MD Experiments) and the unfiltered AF2-MSAss ensemble prior to geometric filtering.

The relative positioning of these ensembles raises an important question on how to separate genuine population shifts from structural implausibility. To distinguish these scenarios, we quantified optimisation reproducibility across replicate measurements. Since optimisation confidence and parameter uncertainty are inversely related, we computed the multivariate standard deviation (*σ*) of Work_scale_ and Work_density_ across the three data-split replicates (Figure S11). These metrics capture fundamental ensemble-level properties: Work_scale_ measures overall shifts in protection factor magnitude (reflecting systematic over- or under-prediction of exchange rates), while Work_density_ measures the intensity of protection factor reorganisation (reflecting how much the ensemble-average distribution must shift to match data). Low multivariate spread indicates the optimizer converges reproducibly regardless of which specific peptides comprise training versus validation sets, suggesting the ensemble contains primarily valid structures that can be consistently reweighted. High spread indicates fitting instability, different peptide selections drive the optimizer toward incompatible solutions. This suggests structural implausibility or severe conformational imbalance.

MD-1Start proved both representative (requiring small Work Done) and confident (achieving reproducible fits) in both structural scales (*σ* = 0.03 for both splits, Figure S11.C), confirming equilibrium MD from high-quality crystal structures provides reliable conformational sampling for this rigid protein. In contrast, AF2-Filtered showed modestly improved confidence in Spatial splits (*σ* = 0.02) but substantially reduced confidence in Non-Redundant splits (*σ* = 0.40, Figure S11.B) comparable to MD-10Start’s Non-Redundant performance (*σ* = 0.49, Figure S11.D). This asymmetry reveals distinct strengths. MD-1Start produces equally reliable fits at both global and local structural scales, whereas AF2-Filtered fitting is robust primarily for sub-structural regions. The confidence pattern suggests that despite lower reproducibility, AF2-Filtered’s global fitting behaviour resembles MD-10Start, both ensembles require substantial shifts in protection factor magnitude to achieve global consistency (high Work_scale_ in Non-Redundant splits), indicating they systematically over- or under-predict exchange rates even when local conformational heterogeneity is reasonably captured.

This confidence analysis exposed distinct requirements for different ensemble types during optimisation. Despite lower global reproducibility, MD-10Start showed similar Work_scale_ to AF2-Filtered across most replicates in Non-Redundant splits (Figure S11.D versus .B), indicat-ing both ensembles require comparable magnitude adjustments to fit global HDX patterns despite AF2-Filtered’s superior local density coverage. Conversely, MD-1Start required minimal Work_scale_ in both splits yet showed larger Work_density_ in Spatial splits, demonstrating that while its predicted exchange magnitudes match experimental observations accurately at all scales, local conformational differences still demand reorganisation of protection factors to capture regional heterogeneity. MD-10Start’s overlapping behaviour between Non-Redundant and Spatial splits in both Work_scale_ and Work_density_ space indicates a conformational density estimate that is skewed. This means not incorrect in structure quality, but imbalanced in population distribution. This is consistent with this ensemble containing diverse yet individually plausible conformations that collectively require rebalancing.

Meanwhile, MD-TFES demonstrated that large protection factor reorganisation requirements (Work_density_) do not necessarily indicate structural implausibility when accompanied by reasonable fitting confidence. Despite requiring substantial conformational reorganisation, MD-TFES maintained better reproducibility than MD-10Start (*σ* = 0.23 to 0.38 versus 0.31 to 0.49, Figure S11.E), with Work_density_ approaching, but remaining distinctly lower than, the unfiltered AF2-MSAss ensemble (Figure S11.A, *σ* = 1.30 to 1.35). As MD-TFES represents our broadest coverage of geometrically valid structures through enhanced sampling of conformational frontier regions, and thus our most deliberately non-equilibrium density estimate for a small, relatively rigid protein like BPTI, this comparison demonstrates a key capability of our framework: distinguishing genuine conformational diversity present in valid but unnatural density estimates (MD-TFES, which maintains fitting stability) from structural implausibility arising from geometric violations (unfiltered AF2-MSAss, which shows both extreme Work Done requirements and fitting instability). The combination of Work_density_ magnitude and confidence metrics thus provides orthogonal information. Large protection factor reorganisation accompanied by reproducible fitting indicates sampling bias rather than invalid structures, whilst irreproducible fitting indicates geometric problems.

These results establish that quantitative comparison across multiple ensembles enables robust structural hypothesis evaluation without requiring detailed prior system knowledge or direct pairwise ensemble comparisons. By systematically combining data partitioning strategies that probe different structural scales (global Non-Redundant versus local Spatial) with Work Done metrics that capture distinct optimisation requirements (magnitude shifts via Work_shape_/scale versus protection factor reorganisation via Work_density_) and confidence assessments that distinguish sampling artifacts from structural problems (multivariate spread across replicates), structural proposals can be ranked and differentiated as an integrated set rather than in isolation. While BPTI benefits from extensive literature characterisation enabling retrospective validation, this multi-scale, multi-metric analysis provides a practical blueprint for evaluating HDX-MS data against structural ensembles in systems where a structural ground-truth remains unknown, true for most experimental applications. We therefore recommend that others adopt similar complementary assessment approaches when comparing structural hypotheses derived from different generation methods, particularly when traditional validation metrics based solely on predicted-experimental agreement fail to provide adequate discrimination between plausible alternatives.

### 2.4 Replicates facilitate uncertainty quantification in model parameter optimisation

Having established our framework on BPTI using structural reweighting, we investigated whether individual structures could be evaluated using model parameter optimisation alone, without the computationally expensive ensemble reweighting employed previously. We hypothesised that model parameter fitting, which adjusts only the global scale factors controlling uptake magnitude to account for experimental conditions rather than reorganising protection factors, would exhibit increased variance in the modifications required during optimisation. To test this hypothesis, we performed model parameter optimisation experiments fitting solely the parameters that scale predicted uptake curves to match experimental conditions from default model parametrisation, as detailed in Equation BV Model within Reweighting and Optimisation Methods. Since model parameter optimisation requires considerably less computational expense than structural reweighting (approximately one-third of computation time spent on structural feature calculation), we performed 10 replicates for both data splits to account for this reduced complexity. Importantly, because model parameter fitting operates at the ensemble-average level without modifying individual frame weights, we could evaluate not only complete ensembles but also individual structures in isolation. Experiment details can be found in Case Study Protocols.

For this investigation we selected the flexible ubiquitinase HOIL-Interacting Protein-1 (HOIP),[38] a system where conformational flexibility is essential to catalytic function but for which no relevant crystal structure exists. This absence of a reference structure made HOIP an ideal test case for our evaluation framework. We directly compared two structures, shown in Figure 4.A: the highest confidence structure from ColabFold using default MSA settings (AF2-MaxRank, measured by predicted Local Distance Difference Test or pLDDT) and the highest scored structure from the entire AF2-MSAss ensemble (AF2-MaxPLDDT). Both structures are physically plausible and could potentially exist on HOIP’s conformational landscape. AF2-MaxRank adopts an extended conformation that matches the HOLO complex structure obtained using nanobodies as crystallisation co-factors (PDB: 6SC6),[38] representing a direct example of AlphaFold2 overfitting to its training set by reproducing crystallographically stabilised conformations unlikely to populate significantly in solution. In contrast, AF2-MaxPLDDT presents a more compact structure with visibly less accessible surface area, which qualitatively appears more probable for the APO state in solution. Despite being a more confident and plausible prediction, the known flexibility of HOIP suggests this structure alone cannot fully represent the *in vitro* dynamics.

**Figure 4:**
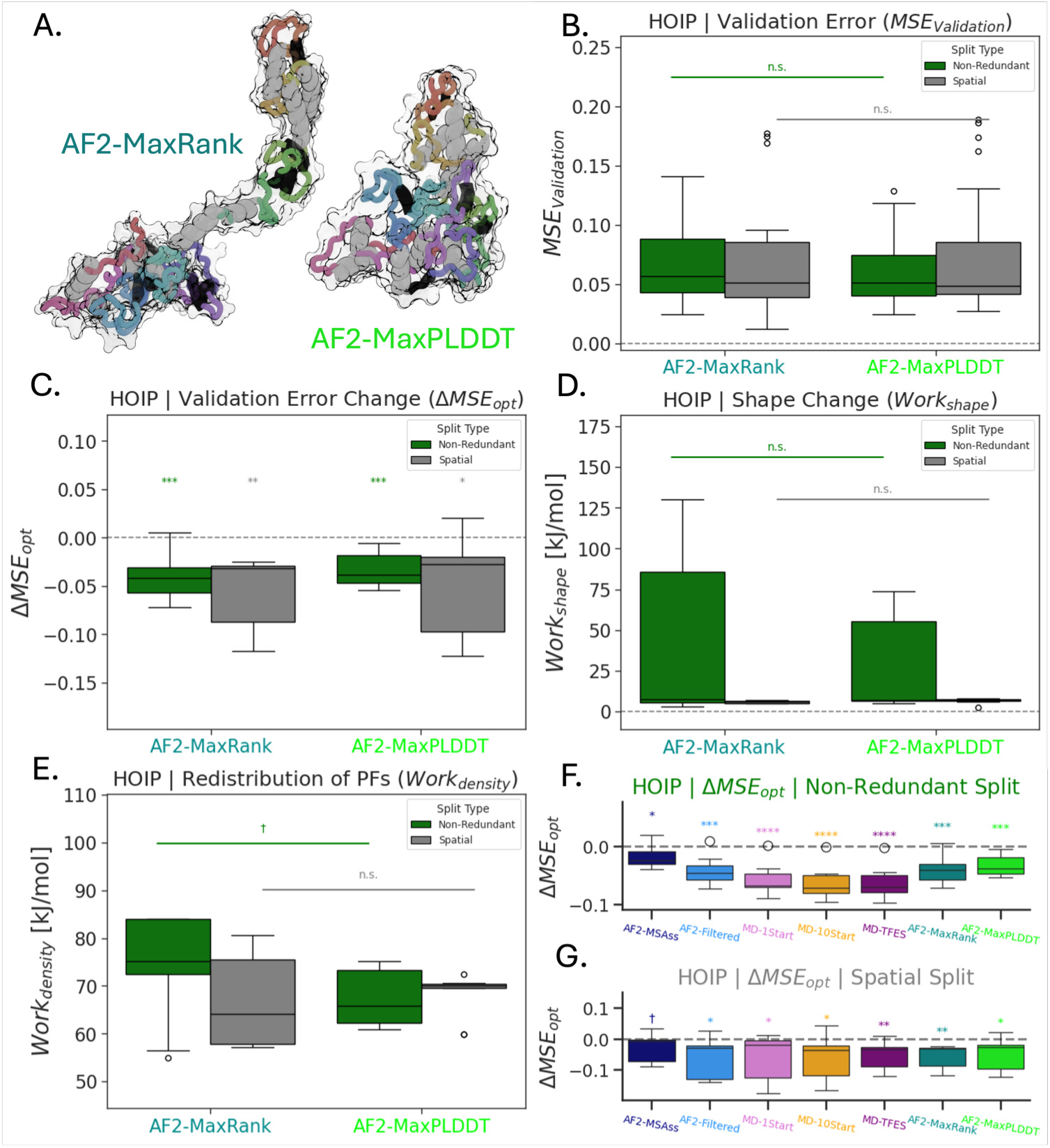
Model parameter optimisation demonstrates replicates facilitate uncertainty quantification in HOIP case study. **(A)** Cartoon representations of AF2-MaxRank (left, forest green) showing extended conformation and AF2-MaxPLDDT (right, neon green) showing compact APO structure, both coloured along residue index with alpha helices (grey) and beta sheets (black). **(B)** Validation error (MSE_Validation_) shows modest but significant differences between structures for both Non-Redundant (green) and Spatial (grey) splits. **(C)** Validation error change upon optimisation (ΔMSE_opt_) demonstrates two orders of magnitude larger scale changes compared to structural reweighting, reflecting lack of regularisation in model parameter optimisation. **(D)** Prediction magnitude adjustments during optimisation (Work_shape_) reveals extremely large values in some Non-Redundant replicates, indicating high variability and lack of certainty in global-level fitting. **(E)** Protection factor reorganisation requirements during optimisation (Work_density_) shows both structures are similar at local density estimates but far from complete coverage, with AF2-MaxPLDDT showing better global conformational density estimation. Comparison of ΔMSE_opt_ across all structural hypotheses for **(F)** Non-Redundant and **(G)** Spatial splits demonstrates that individual structures perform similarly to AF2-Filtered ensemble but show less significant differences than MD ensembles, while AF2-MSAss ensemble shows least change upon optimisation despite containing faulty structures. Results indicate that unconstrained model parameter optimisation may not reliably identify ensemble quality, contrasting with Maximum Entropy structural reweighting approaches.

We first examined validation error to assess whether traditional metrics could distinguish between these structures. Despite the questionable plausibility of the extended AF2-MaxRank conformation in solution, validation error (MSE_Validation_, Figure 4.B) showed no significant differences between structures (*p* = 0.267 to 0.530), consistent with our observations in Figure 3. However, when we examined error change upon optimisation (ΔMSE_opt_, Figure 4.C), we ob-served considerably higher significance than for validation error alone (*p* = 0.0103 to 0.000 335). Moreover, compared to structural reweighting with BPTI, the scale of ΔMSE_opt_ with model parameter optimisation on HOIP was two orders of magnitude larger, which we attribute to the lack of regularisation in model parameter optimisation within HDXer.

We then examined our Work Done metrics to probe structural quality more deeply. For prediction magnitude changes (Work_shape_, Figure 4.D), we observed increased values for both splits relative to the BPTI case, with the Non-Redundant split showing extremely large values in some replicates. This variability across replicates indicated a lack of certainty in the fit for both structures, exacerbated by the unconstrained nature of model parameter optimisation.

For protection factor reorganisation requirements (Work_density_, Figure 4.E), we observed that both structures showed similar behaviour at the local structural scale, with low significance between structures across both splits (Non-Redundant: *p* = 0.0923, Spatial: *p* = 0.591). The considerably large Work_density_ values confirmed these individual structures are far from complete conformational coverage, as expected for highly flexible proteins represented by single conformations. Encouragingly, the more physically plausible AF2-MaxPLDDT structure, which also has higher confidence by pLDDT, showed a better estimate of global conformational density.

To contextualise these individual structure results within our broader ensemble experiments, we compared them across all HOIP structural hypotheses (Figure 4.F-G). The complete cross-ensemble comparisons can be found in section Detailed Ensemble Confidence Assessment. Examining the ensembles, AF2-Filtered (Figure S12.B) showed the second highest local confidence (*σ* = 6.92) but the least global confidence (*σ* = 18.52). Interestingly, AF2-MSAss (Figure S12.A) reported as the most confident ensemble in the Non-Redundant split (*σ* = 11.98), demonstrating that model parameter optimisation is insensitive to individual structure validity since fitting occurs at the ensemble-average level without distinguishing contributions from individual frames.

To extend our analysis beyond uncertainty within individual splits, we developed a measure of agreement between data splits by computing the RMSD of representative average values of Work_density_ and Work_scale_. Agreement between splits suggests both Non-Redundant and Spatial splits provide similar information on fitting success, which can probe structural deficiencies present at both global and local structural-average levels. The use of 10 replicates enabled robust quantification through median values. Despite containing only a single structure, AF2-MaxRank (Figure S12.F) showed better agreement (7.92 kJ/mol) than most ensembles, though extremely high Work_scale_ in some Non-Redundant replicates implied large magnitude adjustments in HDX-uptake predictions are required to fit the global conformation. In terms of median performance, AF2-MaxRank appeared comparable to the AF2-MSAss ensemble (Figure S12.A). While AF2-MSAss showed smaller, more physically plausible changes across many metrics, multiple explanations exist for this behaviour since model parameter optimisation operates solely at the ensemble-average level, including protection factor blurring from averaging over diverse structures or magnitude differences from implausible structural geometries. We therefore conclude that model parameter optimisation cannot provide information at the individual-frame level of an ensemble.

Therefore, the most promising structural hypothesis without reweighting was MD-TFES (Figure S12.E). Despite showing the largest Work_density_ across all hypotheses, MD-TFES demonstrated strong confidence (*σ* = 15.92 to 17.28) in both splits relative to other valid ensembles, particularly in the Non-Redundant split (*σ* = 15.92 to 18.52). Importantly, MD-TFES showed the best split-type agreement amongst ensembles (6.18 kJ/mol), contrasting with MD-10Start which exhibited similar Work_scale_ but poor agreement (18.08 kJ/mol) and worse confidence (*σ* = 17.27 to 17.44). Since MD-TFES contains by design the most diverse range of valid conformations, its strong performance across ensembles aligns with the known flexibility and dynamics of HOIP in solution.

It is encouraging that unconstrained model parameter optimisation can be sensitive to both conformational shape and density representativeness. However, when compared to regularised structural reweighting, we found that valid and invalid ensembles are not as clearly separated by Work Done performance with model parameter optimisation alone.

From these results, we conclude that model parameter optimisation should be performed only on ensembles known to contain plausible structures. When implausible structures are excluded, our results demonstrate that leveraging data splits and replicates enables structural ensemble hypotheses to be quantitatively separated and ranked, though with less reliability than Maximum Entropy reweighting approaches. We therefore suggest that researchers using model parameter optimisation carefully validate ensemble quality before fitting, employ multiple replicates to quantify uncertainty, and preferentially use regularised structural reweighting approaches when computational resources permit.

### 2.5 Optimisation protocol choice affects ensemble-model reliability

Even when working with a structurally representative ensemble, the procedure used to optimise model parameters can itself introduce overfitting. This question carries practical importance because accurate BV-model parameters are essential to unlocking the full potential of HDX-MS.

HDX-MS’s key advantage is the ability to conduct experiments under diverse conditions. These parameters control how experimental conditions (pH, temperature, quench efficiency) affect exchange rates, yet their optimisation operates at the ensemble-average level without examining individual structural frames. We hypothesised that this lack of frame-level regularisation would make fitting outcomes more sensitive to the choice of optimisation protocol.

Previous work has suggested that different protocols for optimising ensemble-model parameters produce comparable performance.[15] To test this claim systematically, we designed an experiment comparing how different protocols perform when optimising model parameters and frame weights in stages. All protocols were fitted to a single parent ensemble, allowing us to isolate the effects of protocol design from ensemble quality. We selected the AF2-Filtered ensemble from BPTI because BPTI is well characterised and, based on Ensemble representativeness assessment reveals scale-dependent structural quality, we were confident this ensemble adequately represents the conformational landscape with sufficient structural coverage. This represents an ideal scenario for model parameter optimisation, allowing us to attribute any performance differences specifically to protocol design rather than deficiencies in the underlying ensemble.

We tested five distinct optimisation protocols, summarised in Table 1. The three baseline approaches were standard HDXer protocols: BV-only (model parameters alone), RW-only (Max-ENT reweighting alone), and MutualBVRW (both methods applied successively at each iteration until convergence). To investigate what happens at each stage of combined approaches, we created two additional protocols. We took the optimised ensembles resulting from BV-only and RW-only and applied the complementary method as a second step, yielding BVafterRW and RWafterBV. Because our Work Done metrics measure changes from the most recent optimisation step, we additionally computed the total distance from the original unoptimised ensemble for these two-step protocols, designating these as BVafterRW* and RWafterBV*. The protocols are distinguished visually in Figure 5 through individual colours and cross-hatching patterns that denote how the methods combine. Full experimental details are provided in Case Study Protocols.

**Figure 5:**
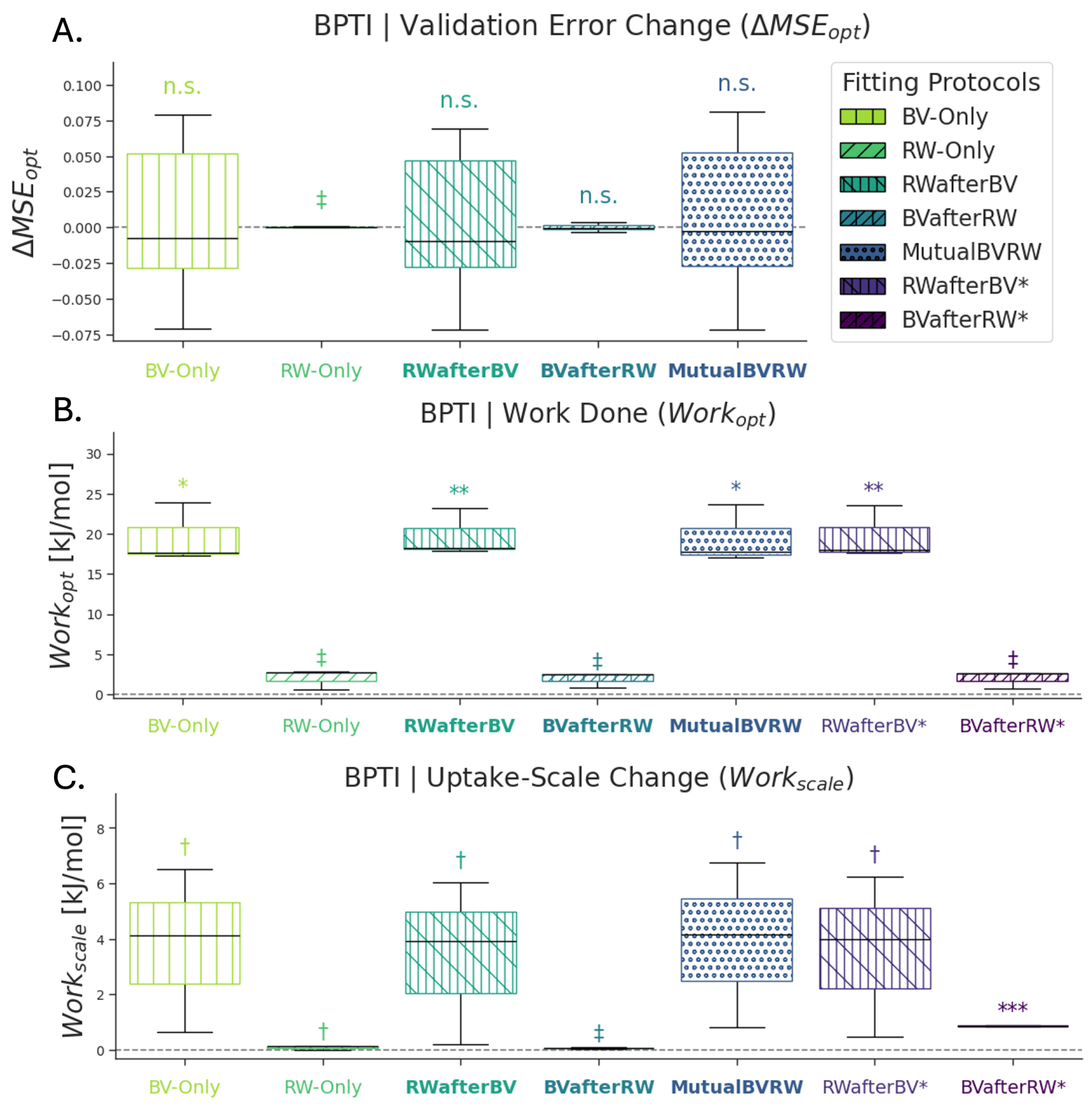
HDXer protocol comparison demonstrates optimisation protocols can lead to overfitting in BPTI case study. Comparison of five optimisation protocols applied to AF2-Filtered ensemble: BV-Only (lime-yellow), RW-Only (green), RWafterBV (teal), BVafterRW (blue), and MutualBVRW (dark blue with dots). Cross-hatching patterns denote composite methods combining individual optimisation approaches. For multi-stage protocols (RWafterBV and BVafterRW), values shown indicate the most recent fitting stage. Therefore, RWafterBV* (purple) and BVafterRW* (dark purple) are provided for Work Done metrics that show the total distance from the original unoptimised ensemble-model. **(A)** Validation error change upon optimisation (ΔMSE_opt_) shows considerable spread for BV-only, RWafterBV, and MutualBVRW protocols. Low significance for BV-containing protocols indicates sensitivity to training peptide selection. **(B)** Free energy change during optimisation (Work_opt_) reveals BV optimisation leads to overfitting (large values) in all cases except when RW-only is performed first (BVafterRW and BVafterRW marked with asterisks). Two-step protocols show ensemble representativeness is important for confident BV-model parameter fitting. **(C)** Enthalpy of scale change (Work_scale_) demonstrates similar trends with overlapping ranges between RWafterBV and other model parameter protocols. Results indicate that unconstrained BV-only optimisation is prone to over-fitting, while sequential application of RW-only followed by BV-only (BVafterRW) provides the most robust parameter optimisation strategy, confirming that protocol choice significantly affects ensemble-model generalisability even with representative ensembles.

**Table 1:** Protocol Comparison: Optimisable Parameters and Distances Measured.

| Fitting Protocol | BV-Only | RW-Only | RWafterBV | BVafterRW | MutualBVRW | RWafterBV* | BVafterRW* |
| --- | --- | --- | --- | --- | --- | --- | --- |
| BV-Parameters | ✓ |  | ✓ | ✓ | ✓ | ✓ | ✓ |
| Frame-Weights |  | ✓ | ✓ | ✓ | ✓ | ✓ | ✓ |
| Protocol Steps | 1 | 1 | 2 | 2 | 1 | 2 | 2 |
| Work Contribution | BV-Only | RW-Only | RW-Only | BV-Only | Both | Both | Both |

We first examined validation error changes (ΔMSE_opt_) across protocols, shown in Figure 5.A. Despite using a known representative ensemble, we observed considerable variability in results. The error distributions showed substantial spread for BV-only, RWafterBV, and MutualBVRW protocols, with RW-only being the only approach showing marginal significance (*p* = 0.0831) compared to the other protocols (*p* = 0.500 to 0.907). This low significance in error changes reinforced our earlier finding that unconstrained BV-only optimisation is sensitive to which peptides comprise the training set. Notably, BVafterRW exhibited the lowest significance (*p* = 0.907) yet also the smallest error spread, providing initial evidence that carefully staged BV-only optimisation following reweighting might locate optimal parameters without overfitting.

To obtain more precise information about optimisation success, we measured the free energy changes during optimisation (Work_opt_), presented in Figure 5.B. Here, we observed a clear and consistent pattern. BV optimisation led to overfitting (indicated by large Work_opt_ values) in all cases except when RW-only had been performed first. The BVafterRW and BVafterRW* protocols showed markedly lower work values, suggesting that reweighting beforehand constrained the parameter search space effectively. Among protocols incorporating BV optimisation, BV-only and MutualBVRW showed similar significance levels (*p* = 0.0116 to 0.0113), while RWafterBV and RWafterBV* demonstrated the strongest significance (*p* = 0.007 53 to 0.009 45), indicating that BV-model parameter changes dominate the protection factor modifications during ensemble optimisation in most scenarios.

We next examined changes in the uptake scale (Work_scale_), shown in Figure 5.C. While the trends paralleled those in Work_opt_, the significance levels differed markedly. Apart from BVafterRW* (*p* = 0.00329), most protocols showed only marginal significance (*p* = 0.0819 to 0.185). Within this marginally significant group, BVafterRW performed slightly better (*p* = 0.0819). These results together suggested that the BVafterRW protocol was most consistently capable of achieving successful fits.

To test the robustness of the optimised parameters more rigorously, we performed a re-optimisation experiment. We took representative BV-model parameters and frame weights from the MutualBVRW, RWafterBV, and BVafterRW protocols and performed an additional BV-only re-optimisation step on each, as detailed in Figure S13.A-C. Because these protocols had already produced ensemble-models with optimised parameters and weights, any changes induced by this additional optimisation step reflected the stability and robustness of the original optimisation outcome.

The re-optimisation results confirmed BVafterRW as the most robust protocol. BVafterRW produced the most certain ensemble-models (*σ* = 1.20 to 3.99) and incurred the smallest median changes in both Work_density_ and Work_scale_ (Figure S13.B). This stability indicated that staged model parameter optimisation following MaxENT reweighting generates the most likely ensemble-models. In contrast, both RWafterBV (*σ* = 6.26 to 6.41, Figure S13.A) and MutualBVRW (*σ* = 6.26 to 6.45, Figure S13.C) showed similar, higher uncertainty levels and larger median work changes in the Non-Redundant split. An interesting contrast emerged in the Spatial split: RWafterBV exhibited larger density changes than MutualBVRW yet showed closer agreement between data splits (1.48 kJ/mol) compared to BVafterRW (1.93 kJ/mol). This pattern suggested that BV-model parameters from RWafterBV were inducing smoothing effects at both global and local levels of the resulting ensemble-models.

These findings demonstrate that forward model parameters meaningfully affect ensemble generalisability. The model parameters, in turn, influence the resulting optimised frame weights. Importantly, we show that suboptimal protocol choices can compromise ensemble-model quality even when the underlying ensemble is already highly representative. The sensitivity of model parameter optimisation to protocol design highlights the value of systematic protocol comparison, even in seemingly ideal scenarios with well-characterised systems. While the BVafterRW protocol (performing Maximum Entropy reweighting before model parameter optimisation) appears to provide the most robust optimisation strategy in this case, practitioners should consider that different protocols may be appropriate depending on the specific characteristics of their ensemble and experimental objectives.

### 2.6 Clustering facilitates interpretable ensemble-integration

Having established that BV-model parameter optimisation should be performed on known representative candidate ensembles (Figure 5), we set out to determine whether workflows can be made more interpretable by using a small number of basis-set structures to iteratively build up a picture of dynamics. One of the central difficulties in ensemble optimisation is the heavy redundancy of biomolecular simulation trajectories. Ensemble experiments in this work con-sist of *>* 10, 000 frames each. Not only are these challenging to interpret, but this also adds computational expense to the optimisation process. While trajectory slicing is commonly used to reduce ensemble size, this may not be ideal for ensembles with non-chronological orderings and can remove important but rare conformations. Clustering offers a solution to reduce ensemble size while retaining structural diversity.

We hypothesised that clustering can be used to assess within-ensemble differences. To test this, we generated proportionally sized ensembles (0.001, 0.01, 0.05, 0.1, 0.2, 0.5, and 1.0) and quantified the effect of downsampling on optimisation performance for all ensemble generation experiments. Since clustering effectively reduces the weighting of oversampled regions and removes noise, comparing clustered ensembles against each other allowed us to understand parent-ensembles in the context of conformational density coverage. We applied k-means clustering on PCA coordinates with 10 dimensions computed on the *C_α_* pairwise coordinates for the residues covered by peptides from HDX-MS. Details of dimensionality reduction and clustering can be found in Quantifying Ensemble Similarity and Conformational Coverage, with full experiment details in Case Study Protocols.

To understand the relationship between cluster proportion and ensemble representativeness, we compared ΔMSE_opt_ across MD-1Start, AF2-Filtered and AF2-MSAss ensembles (Figure 6.A–C, respectively). Across both data-splits, we observed that valid ensembles (MD-1Start and AF2-Filtered) become slightly poorer fits when clustering down, as expected from loss of conformational density information. In striking contrast, the implausible AF2-MSAss ensemble showed dramatic improvement in validation error when clustering down from 1.0 to 0.5, and again from 0.05 to 0.001. Within ensembles, AF2-Filtered exhibited closer agreement in trends across splits than MD-1Start, where split-types diverged. We noted that AF2-Filtered appeared to over-fit slightly across the range 0.01 to 0.2, while MD-1Start showed over-fitting below a proportion of 0.1 in the Spatial split only. Comparing error change between AF2-Filtered and AF2-MSAss suggested that the improvement from removing physically implausible structures by clustering can be greater than the loss in density estimates from reduced ensemble weight-ing.

**Figure 6:**
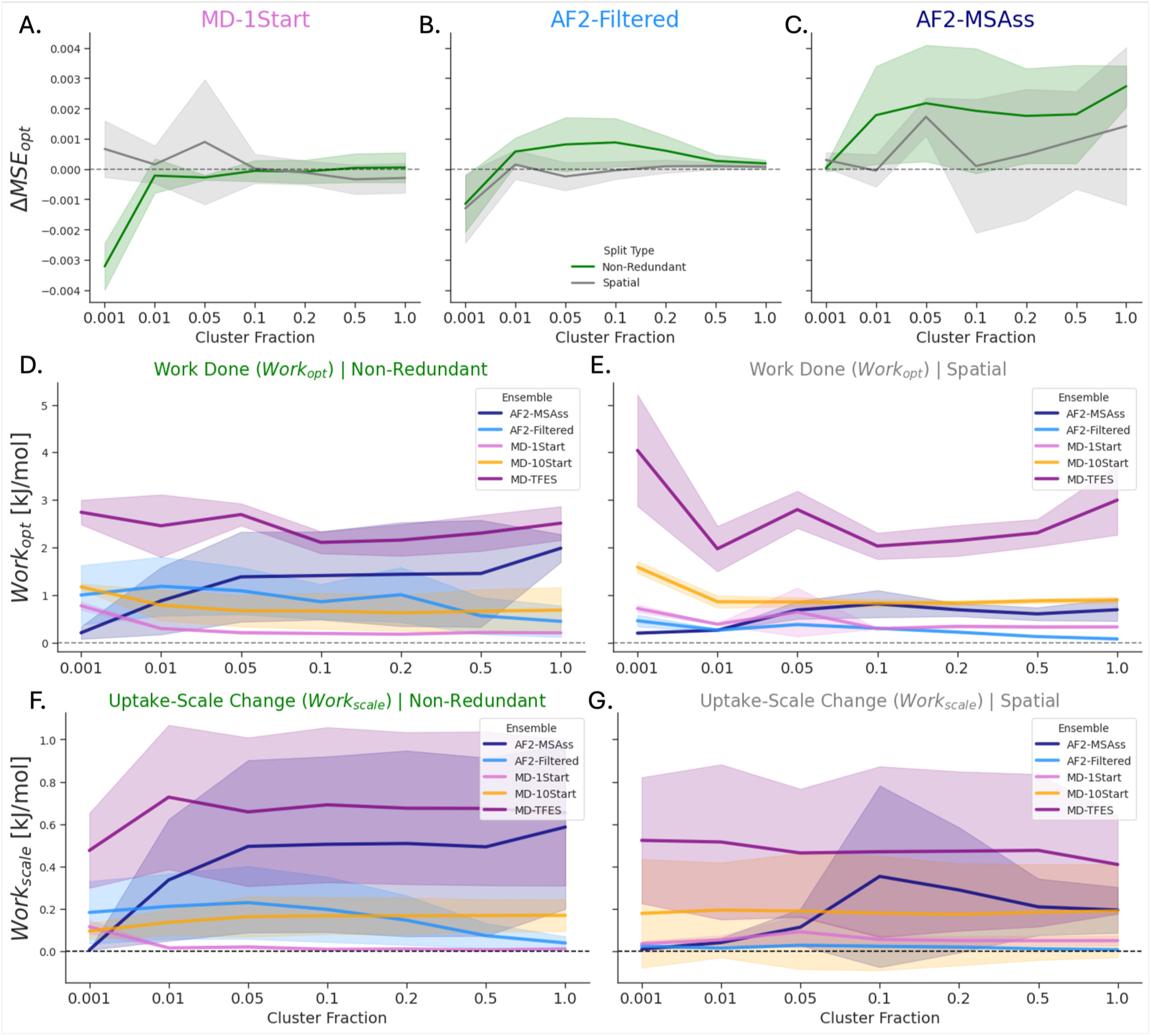
Clustering facilitates interpretable ensemble-integration through systematic down-sampling analysis across BPTI ensembles. Validation error change (ΔMSE_opt_) across cluster proportions (0.001 to 1.0) for **(A)** MD-1Start (pink), **(B)** AF2-Filtered (cyan), and **(C)** AF2-MSAss (navy) ensembles, showing Non-Redundant (green) and Spatial (grey) splits with confidence intervals. Valid ensembles **(A–B)** show slight performance degradation with clustering, while implausible AF2-MSAss **(C)** demonstrates dramatic improvement, particularly from 1.0 to 0.5 and 0.05 to 0.001 proportions. Free energy change during optimisation (Work_opt_) across cluster proportions for **(D)** Non-Redundant and **(E)** Spatial splits reveals AF2-MSAss achieves performance comparable to un-clustered valid ensembles at 0.001 proportion. MD-TFES (purple) shows consistently high but stable performance suggesting systematic deficiency, while MD-10Start (orange) demonstrates robust agreement across splits. Enthalpy of scale changes (Work_scale_) for **(F)** Non-Redundant and **(G)** Spatial splits show MD-TFES and MD-10Start exhibit strong data-split agreement, indicating systematic deviations in overall dynamics scale unaffected by clustering. AF2-MSAss displays large variance and strong correlation to cluster proportion in the Non-Redundant split, indicating uncertainty and a poor structural hypothesis, respectively. These results provide evidence that within-ensemble differences can be used to build a picture of ensemble quality. However, conclusively distinguishing implausible (AF2-MSAss) from noisy (MD-TFES) structural hypotheses requires multiple data-splits and metrics.

To investigate this observation further, we examined Work_opt_ across the cluster sweep (Figure 6.D–E for Non-Redundant and Spatial splits, respectively). The Non-Redundant split (Figure 6.D) confirmed that AF2-MSAss is massively improved by clustering. At a proportion of 0.001, AF2-MSAss showed similar performance to the unclustered (1.0) AF2-Filtered and MD-1Start ensembles. Across most cluster proportions, MD-TFES exhibited high but consistent Work_opt_, suggesting that this structural hypothesis contains some systematic deficiency. The lower variance for MD-TFES relative to AF2-MSAss demonstrated successful disentangling between “oversampled” and physically invalid ensembles at the global level. In the Spatial split (Figure 6.E), MD-TFES proved most sensitive to cluster proportion, while AF2-Filtered, MD-1Start and MD-10Start performed as expected for representative ensembles, requiring additional work at smaller cluster proportions. From these data, MD-10Start showed the most consistent agreement between split-types, indicating a robust proposal of conformational density for this protein-ensemble combination.

Given that the clustered ensemble structural-density distribution can be transformed more dramatically than simply reduced weighting, we recognised that Work_opt_ alone is not sufficient to rank ensembles, particularly for MD-TFES which showed variability across splits and cluster proportions. We therefore compared the scale changes (Work_scale_) across split-types (Figure 6.F–G). These revealed consistent and strong data-split agreement for MD-TFES and MD-10Start ensembles, indicating that there is some systematic deviation in these ensembles related to the overall scale of the dynamics. This deviation is not affected by clustering and is present at both global and local structure levels. This suggested that the sensitivity in Work_opt_ for MD-TFES could be attributed to variability in conformational-density when clustering to different proportions. For AF2-Filtered and MD-1Start ensembles, we observed small differences in uptake scale changes (Work_scale_) over cluster proportions with some split-type disagreement, though this was lower than for the AF2-MSAss ensemble. The large variance in scale change and strong positive correlation to cluster proportion for AF2-MSAss in the Non-Redundant split indicated uncertainty and a poor structural hypothesis, respectively. These results provided evidence that within-ensemble differences can be used to build a picture of ensemble quality. However, conclusively distinguishing implausible (AF2-MSAss) from noisy (MD-TFES) structural hypotheses requires multiple data-splits and metrics.

From these experiments, we conclude that representative ensembles can be significantly down-sampled to proportions between 0.1 and 0.2 with minimal difference to the implied quality. These proportions result in clustered ensembles of approximately 1000 to 2000 structures. Valid ensembles show slight degradation with lower cluster numbers while implausible ensembles (AF2-MSAss) improve, likely due to the removal of invalid structures. In both cases, this provides grounding to optimise on heavily reduced ensemble sizes for interpretability. We therefore suggest that, for implausible ensembles, care should be applied for low cluster numbers as the clustered ensemble may not reflect the full diversity in the parent ensemble. For valid ensembles, a proportion of 0.01 (100 to 127 structures) could result in useful models. In some cases parent ensembles can be clustered down further to a proportion of 0.001, resulting in a human-interpretable number of 10 to 13 structures for ensemble-integration workflows.

### 2.7 Systematic structural-artifacts reveal distinct ensemble deficiencies in conformational sampling and physical plausibility

Having demonstrated in the previous experiment that clustering can probe within-ensemble structural details, we set out to test whether controlled structural-artifacts could more precisely isolate and quantify different aspects of ensemble quality. While clustering simultaneously affects both ensemble composition and structural distribution, we hypothesised that synthetic conformations with specific, well-defined characteristics would allow us to independently assess conformational sampling completeness versus structural plausibility. By introducing structures that mimic real-world ensemble failures (missing unfolded states, physically invalid geometries, or inaccurate hydrogen bonding patterns), we could determine whether our validation framework reliably detects these distinct failure modes.

We chose Bromodomain-containing protein 4 (BRD4) as our test system because it represents a particularly challenging validation scenario. BRD4 consists of two Bromodomains separated by proline and glycine-rich loops that confer substantial intrinsic disorder. This architecture produces an exceptionally broad conformational distribution in solution, making complete sampling unlikely in the ensemble generation experiments performed in this study. Additionally, the proline-rich regions severely limit HDX-MS peptide coverage to just the Bromodomain units themselves. This combination of extensive conformational heterogeneity and sparse experimental coverage represents a realistic test case for precisely probing ensemble quality when validation information is limited.

We designed three types of synthetic structural-artifacts, each targeting a distinct failure mode commonly encountered in ensemble generation. First, we applied Gaussian coordinate noise (Figure 7.A), which mimics unfolded or highly flexible states by inducing global structural perturbations. Second, we performed complete coordinate mixing (Figure 7.D), creating entirely unphysical structures that smooth protection factors across residues and violate fundamental structural constraints. Third, we randomly shuffled the backbone proton identities used to calculate hydrogen-bond acceptor contacts in the BV-model (Figure 7.G), generating structures with altered protection-factor profiles while maintaining overall structural integrity. To ensure systematic evaluation, we first clustered each parent ensemble to exactly 1000 structures, then successively added synthetic structures at increasing proportions: 1, 10, 100, and 500 structures, corresponding to 0.1%, 1.0%, 10%, and 50% of the original ensemble size. The synthetic structures were added in order of their deviation from ensemble-average coordinates, quantified by principal component analysis of pairwise *C_α_* distances. We assessed sensitivity through Pearson correlation coefficients (*R*) and evaluated precision through standard deviation (*SD*) of metric values across the first three structural-artifact levels (0.1%, 1.0%, 10%), treating the 50% level as a negative control that was excluded from these summary statistics. To facilitate comparison across ensemble types and structural-artifacts modes, we normalised all metrics as Z-scores relative to the unmodified parent ensemble (0.0%). Complete descriptions of artifact generation algorithms are provided in Structure Poisoning Experiments, with experimental protocols detailed in Case Study Protocols.

**Figure 7:**
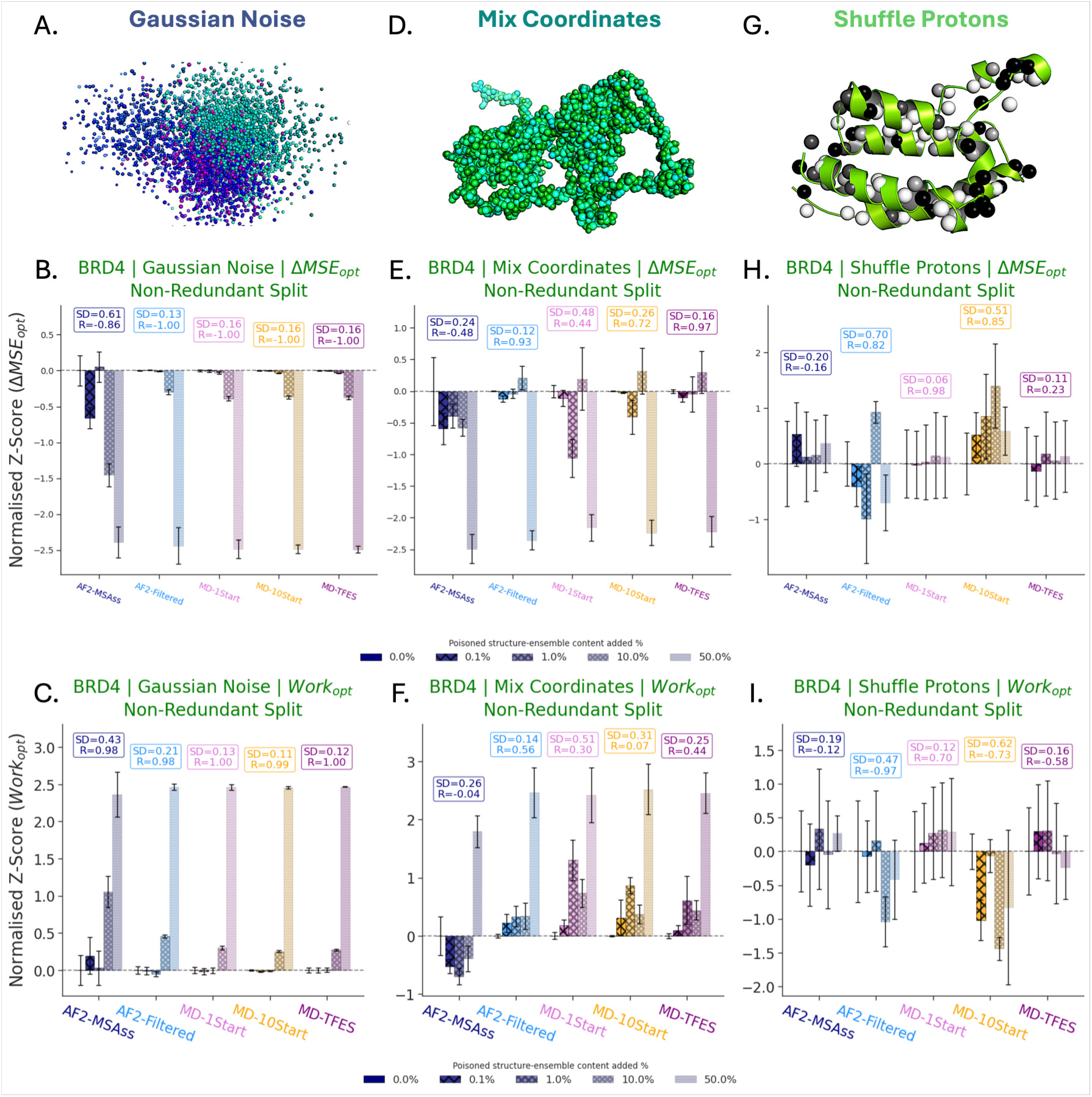
Systematic structural-artifacts reveal distinct ensemble deficiencies through controlled perturbation experiments in BRD4. Three synthetic-artifact types applied to clustered ensembles (k=1000) with successive addition at 0.1%, 1.0%, 10%, and 50% levels: **(A)** Gaussian coordinate noise mimicking missing unfolded states, **(D)** complete coordinate mixing creating unphysical structures that violate fundamental constraints, and **(G)** backbone proton shuffling generating altered protection-factor profiles. Standard deviation (SD) and correlation (R) values quantify response precision and sensitivity across artifact levels (excluding 50% negative control). Normalised Z-scores relative to unmodified ensembles (0.0%) for validation error change (ΔMSE_opt_) in Non-Redundant splits show **(B)** Gaussian noise reduces optimisation work for most ensembles with AF2-MSAss demonstrating reduced sensitivity and elevated variability, **(E)** coordinate mixing creates severe structural violations with increased ensemble separation, and **(H)** proton shuffling produces diverse response patterns with AF2-Filtered showing apparent improvement while others exhibit overfitting signatures. Free energy change (Work_opt_) Z-scores reveal **(C)** Gaussian noise increases optimisation work for all ensembles with pronounced separation between AF2-MSAss and physically valid ensembles, **(F)** coordinate mixing paradoxically improves AF2-MSAss performance while degrading valid ensembles, and **(I)** proton shuffling challenges clear ensemble discrimination with MD-1Start showing highest response consistency across metrics. Results demonstrate that ensemble reweighting detects individual conformation effects at global level, but no single artifact type simultaneously informs on sampling completeness and structural plausibility. Multiple complementary metrics are essential due to complex response patterns. This reveals fundamental BV-model sensitivity limitations in differentiating ensembles from coordinate noise while supporting systematic artifact introduction as a valuable probe for specific aspects of structural hypothesis quality.

We first examined how each type of structural-artifact affected ensemble behaviour, as illustrated in Figure 7. Addition of Gaussian coordinate noise reduced ΔMSE_opt_ (Figure 7.B) for nearly all ensembles tested, indicating these perturbed structures require greater free energy changes during optimisation to achieve comparable fit quality. Valid physical ensembles exhibited tightly correlated responses (*R* = −1.00) with low uncertainty (*SD* = 0.13 to 0.16), while AF2-MSAss showed markedly reduced sensitivity (*R* = −0.86) and substantially greater variability (*SD* = 0.61). This pattern was reinforced by Work_opt_ measurements, which increased with artifact proportion for all ensembles (Figure 7.C), with particularly pronounced separation between AF2-MSAss and the physically valid ensembles. At the ensemble-average level, however, Gaussian noise artifacts produced only marginal metric differences and therefore provided limited power to discriminate among ensembles. The more severe coordinate mixing artifacts created stronger effects, producing increased separation between ensembles for both ΔMSE_opt_ and Work_opt_ (Figure 7.E–F). Remarkably, AF2-MSAss showed apparent improvement in Work_opt_ when these unphysical structures were added, peaking at the 1.0% level and remaining negative across all proportions below 50%. In contrast, MD-1Start exhibited the largest absolute change at 1.0%, while MD-10Start was most affected at the 0.1% level, though the difference was small. AF2-Filtered and MD-TFES showed similar responses to coordinate mixing and were clearly separable from other ensembles below the 50% artifact level. While these results demonstrated sensitivity to individual structures, coordinate mixing provided primarily binary discrimination between plausible and implausible ensembles rather than fine-grained quality assessment. The most discriminating artifact type proved to be proton shuffling, which produced diverse response patterns with substantial separation among ensembles (Figure 7.H–I). Unlike the other perturbations, this mode made it challenging to clearly distinguish physically valid ensembles from the implausible AF2-MSAss ensemble.

From these systematic comparisons, we identified characteristic signatures that reveal distinct aspects of ensemble quality. Analysis of Work_opt_ (Figure 7.I) initially suggested MD-10Start substantially improved with addition of synthetic conformations; however, joint consideration with ΔMSE_opt_ (Figure 7.H) revealed clear overfitting behaviour. AF2-Filtered was unique in showing improvement by ΔMSE_opt_, but exhibited minimal changes in Work_opt_ below 10% artifact content. The high standard deviations observed for AF2-Filtered in both metrics (*SD* = 0.47 to 0.70) indicated substantial uncertainty in this structural hypothesis and suggested under-sampling for this protein system. By contrast, MD-1Start demonstrated exceptional consistency, with low variability (*SD* = 0.06 to 0.12) and strong sensitivity (*R* = 0.70 to 0.85) in Work_opt_. This analysis reinforced the necessity of evaluating multiple metrics simultaneously: examination of ΔMSE_opt_ alone revealed that MD-TFES (*R* = −0.58 to 0.23) showed considerably reduced sensitivity compared to MD-1Start. Notably, MD-10Start exhibited the lowest confidence across both metrics (*SD* = 0.51 to 0.62).

Detailed comparison between MD-1Start and MD-10Start illuminated the relative contributions of simulation length versus conformational diversity to ensemble quality. We concluded that the superior performance of MD-1Start derives from the extended simulation time in a single trajectory rather than from the structural heterogeneity MD-10Start inherits from its diverse AF2-Filtered starting configurations. The reduced sensitivity of MD-TFES to proton shuffling artifacts suggested this ensemble provides a better representation of the true in-solution conformational landscape. Because MD-TFES explicitly samples between local energy minima through seeded exploration, our results indicated these inter-basin conformations are essential for accurately describing BRD4’s solution dynamics.

From this comprehensive investigation, we conclude that no single structural-artifact type can simultaneously inform on conformational sampling completeness and structural plausibility. Within physically valid ensembles, this experiment demonstrated that ensemble reweighting can detect and quantify the effects of individual synthetic conformations at the global optimization level. Despite carefully designed artifacts and systematic metric normalisation, making definitive statements about relative ensemble quality proved challenging, particularly when considering ensembles in isolation. Even for a single artifact type, the response patterns were complex and required multiple complementary metrics to confidently evaluate structural hypotheses. We propose that systematic introduction of synthetic structural-artifacts offers a valuable approach for revealing specific deficiencies in ensemble quality. However, this study also highlighted fundamental sensitivity limitations of the BV-model framework in reliably differentiating between ensembles and simple coordinate perturbations, suggesting the need for orthogonal validation approaches when making strong claims about ensemble accuracy.

## 3 Discussion

We have presented ValDX, a framework for assessing quantitative structural hypothesis from integrative HDX-MS. This framework allows several key advances detailed below.

### Reject ensembles that fit but fail (Iso-Validation, TeaA)

ValDX enables researchers to distinguish between structural hypotheses that achieve identical experimental agreement but contain fundamentally different conformational populations. Our TeaA experiments demonstrated that ISO-Trimodal failed to recover true population ratios (Open State Recovery *<* 40%) despite matching uptake curves as well as ISO-Bimodal, which achieved 80–90% recovery. Work Done metrics detected this failure where training error could not. In practical terms, researchers can confidently reject ensemble proposals that achieve good fits through incorrect structural mechanisms.

### Assess representativeness at multiple structural scales (BPTI)

The combination of Non-Redundant and Spatial splits reveals whether ensembles capture dynamics at global versus local levels. For BPTI, MD-1Start better represented the overall conformational ensemble (lower Work_shape_ in Non-Redundant splits), while AF2-Filtered better captured sub-structural flexibility (lower Work_density_ in Spatial splits). This multi-scale assessment is directly linked to answering a biological question invisible to traditional approaches.

### Rank individual structures without full reweighting (HOIP)

Model parameter optimisation enables rapid evaluation of both single structures and ensembles when reweighting is prohibitive. For HOIP, we demonstrated that AF2-MaxRank adopts an extended conformation matching the crystallographically stabilised HOLO structure, while AF2-MaxPLDDT presents a more compact APO-like state. The framework correctly identified the compact structure (AF2-MaxPLDDT) as more plausible for solution dynamics (Work_density_ indicating better global conformational density). This provides a practical screening tool for selecting simulation starting structures as starting structures can be compared with candidate ensembles in the same framework.

### Avoid overfitting through staged optimisation (Protocol Comparison)

Protocol choice affects ensemble-model reliability even when the underlying ensemble is representative. Our systematic comparison established that performing Maximum Entropy reweighting before model parameter optimisation (BVafterRW) produces the most robust ensemble-models (*σ* = 1.20 to 3.99), while optimising parameters first leads to overfitting. This finding provides a concrete procedural recommendation: reweight first, then optimise parameters.

### Reduce ensembles to interpretable basis sets (Clustering)

Representative ensembles can be clustered from *>*10,000 conformations to 1000–2000 (proportion 0.1–0.2) with minimal impact on implied quality, and in optimal cases to 10–13 conformations (proportion 0.001). This dramatic reduction makes it feasible to visually inspect individual conformations and understand the structural basis for observed dynamics. Importantly, implausible ensembles (AF2-MSAss) improved with clustering through removal of invalid structures, providing a diagnostic: ensembles that improve upon reduction likely contain problematic conformations.

### Distinguish incomplete sampling from invalid structures (BRD4)

Controlled addition of synthetic conformations reveals distinct failure modes. The unfiltered AF2-MSAss ensemble showed extreme Work Done requirements with fitting instability (*σ* = 1.30 to 1.35), indicating geometric violations. MD-TFES showed elevated Work Done with stable fitting, indicating systematic sampling bias rather than structural implausibility. These signatures enable targeted intervention: filter ensembles with implausible structures; extend simulations for ensembles with sampling limitations.

### 3.1 General recommendations

A complete schematic of the ValDX workflow is given in ValDX Workflow Diagram; the general workflow is detailed in ValDX Process Flow along with considerations for common failure modes.

- Optimisation outcomes depend strongly on which peptides and time points are used for fitting; replicates quantify this uncertainty
- Training/validation splits are essential; Non-Redundant and Spatial split-types provide complementary structural perspectives
- Validation error describes generalisability, but Work Done metrics provide the sensitivity required to distinguish structural hypotheses
- Protection factor/HDX-uptake skew (Work_shape_), scale (Work_scale_), and reorganisation (Work_density_) contributions should be separated at the ensemble-average level
- Perform Maximum Entropy reweighting before model parameter optimisation to avoid overfitting
- Cluster ensembles to 1000–2000 structures for efficiency, or to 10–100 basis structures for interpretability
- Validate high-confidence claims by testing sensitivity to synthetic structural artifacts

### 3.2 Ensemble generation considerations

Details of ensemble-generation experiments are provided in Ensemble Generation Experiments. We provide scripts (in Code and Data Availability) for all aspects of ensemble generation; these represent practical working examples rather than optimal protocols. Key observations from our investigations:

- Deep-learning structure predictors may require extensive sampling across parameter choices (MSA subsampling, random seeds) to obtain confident starting structures
- Filter implausible conformations before fitting; this is essential for AlphaFold2 ensembles generated through MSA subsampling
- Well-equilibrated MD from high-quality starting structures produces ensembles with reliable conformational density estimates
- Individual starting structures can be screened using model parameter optimisation before committing to full MD simulations
- Basis-set structures from clustering reduce the combinatorial space for understanding and assessing conformational diversity
- For enhanced-sampling ensembles, set initial population weights to reflect local structure density *a priori*, as the Maximum Entropy prior assumes uniformly important initial populations

#### Concluding remarks

By developing a framework for quantitative validation of structural hypotheses, this work enables rigorous interpretation of HDX-MS measurements in terms of protein conformational dynamics. The ability to distinguish ensembles that fit from ensembles that are correct addresses a fundamental challenge in integrating biophysical data with structural models, providing principled access to the conformational heterogeneity underlying biological function.

## Supporting information

Supplementary manuscript

## 4 Acknowledgements

This work is funded by the Engineering and Physical Sciences Research Council (EPSRC) with grant code EP/S024093/1. This work was also funded by OMass Thereapeutics. OMC acknowledges funding from a New College Todd-Bird Junior Research Fellowship and MRC Fellowship MR/Y010078/1. He acknowledges consulting fees to Pelago Biosciences, Faculty.ai and Market-Cast and is on the scientific advisory board of Evolvere Biosciences. No funder had a role in the research or decision to publish

## 5 Materials and Methods

### 5.1 HDX Data

To ensure robust fitting, HDX-MS data requires preprocessing to mitigate inherent limitations: (1) peptide redundancy can lead to data leakage during validation, (2) uptake curves at extreme time points (e.g., 0 min, saturation) are poorly modeled by single-exponential kinetics, and (3) short peptides provide low information content. Our filtering protocol addresses these limitations before any splitting or optimisation occurs.

We obtained published HDX-MS datasets for five protein systems (HDX-MS Data Summary). These systems cover a diverse range of structural properties, including rigid and flexible proteins, as well as those with only partially complete experimental structures.

**Methods Table M2:** HDX-MS Data Summary.

| System | Name | Type and Source | No. Peptides | Time Points /min |
| --- | --- | --- | --- | --- |
| BPTI | Bovine Pancreatic Trypsin Inhibitor | NMR, APO[13] | 41 | 0.167 1.0 10.0 120.0 |
| BRD4 | Bromodomain containing protein 4 | Both Constructs[39] | 104 | 0.0 0.25 1.0 10.0 60.0 240.0 |
| HOIP | HOIL-1L Interacting Protein | APO[38] | 108 | 0.0 0.5 5.0 |
| MBP | Maltose Binding Protein | APO[40] | 115 | 0.5 4.0 30.0 240.0 |
| LXRa | Liver X Receptor alpha | APO[41] | 87 | 0.5 10.0 |

#### 5.1.1 HDX-MS data processing

To ensure that the data used for ensemble optimisation was informative and compatible with the Best-Vendruscolo (BV) model,[42] we applied the following filtering criteria:

- **Time point filtering:** To exclude poorly informative time points where the single-exponential BV model is inaccurate, we removed time points where the replicate precision (RSD) exceeded 5%, typically corresponding to very short or very long deuteration times. This threshold aligns with high-precision HDX-MS estimates.[43]
- **Peptide filtering:** We removed peptides that were short (*<* 5 residues) or had low information content (saturation difference *<* 0.05).

All filtering was performed before any splitting or optimisation. Protocols for all ensembles generated can be found in Ensemble Generation Experiments, and subsequent ensemble analysis is detailed in Additional Experiment Plots.

##### Assessing Global vs. Local Scales through Peptide Splitting

Because peptides share residues, a validation peptide can be partially ’known’ from training peptides covering similar regions. For example, if residue 50 is measured by five different peptides, information about that residue leaks between training and validation sets, inflating apparent performance. We therefore define splits that minimise shared residue coverage. To test whether an ensemble accurately represents protein dynamics at different structural scales, we designed splitting strategies that isolate distinct types of information.

##### Non-Redundant Split (Global Scale)

This split clusters peptides by sequence region, ensuring validation peptides test entirely unseen segments of the protein. This tests the ensemble’s ability to predict global conformational behavior without relying on local overfitting. Any peptide that shares ≥ 1 residue with a peptide in the other set is removed from the training set. Specifically, using the start and end positions of each peptide as features, a k-means clustering algorithm is applied using a tenth of the total number of peptides as the number of clusters. Clusters are then randomly assigned according to the specified training fraction, remaining dataset-overlapping peptides are then dropped.

##### Spatial Split (Local Scale)

This split withholds peptides covering contiguous 3D regions of the protein structure. This tests the ensemble’s accuracy in capturing local substructural dynamics, such as domain motions or specific binding site flexibility. A residue is randomly selected from the protein, the remaining residues are ordered by the distance of their *C_α_*atom. The training set is filled in order of proximity of peptide-residues. Overlapping peptides are then dropped.

##### Random and Sequential Splits (Baselines)

We also implemented standard random and sequential splits as baselines to demonstrate the insufficiency of methods that do not account for information leakage. These splits typically result in high overlap between training and validation sets. For the random split, peptides are randomly selected for the training set and the remainder used in the validation set. For the sequential split, peptides are ordered based on sequence number and are then selected up to the training fraction; the remainder is used in the validation set.

All splitting algorithms ensured a 50% training fraction, with both training and validation sets containing at least 10% of the original peptides. To improve diversity and prevent redundancy-driven bias, peptides with high overlap density were stochastically dropped (10% dropout rate) prior to splitting.

### 5.2 Ensemble Generation Experiments

To test the ValDX framework, we generated a diverse set of structural ensembles for each protein system. These ensembles were designed to represent distinct scientific hypotheses about how to best capture protein dynamics, ranging from single-structure simulations to evolutionary-based predictions and enhanced sampling. All ensembles were generated with approximately equivalent computational budgets (≈ 100 ns of simulation time) to ensure fair comparison of sampling strategies rather than computational investment. A summary of all generated ensembles is provided in Summary of Structural Ensembles and Hypotheses.

**Methods Table M3:** Summary of Structural Ensembles and Hypotheses.

| Ensemble | Generation Method | Frames | Clustered Size | Hypothesis Tested |
| --- | --- | --- | --- | --- |
| MD-1Start | Single-start equilibrium MD | $\sim 10,000$ | 500 | Conventional MD from a single high-quality structure |
| MD-10Start | Multi-start equilibrium MD | $\sim 10,000$ | 500 | Improved sampling via diverse starting points |
| MD-TFES | Frontier expansion sampling | $> 30,000$ | 500-1000 | Maximal diversity sampling strategy |
| AF2-Filtered | MSA subsampling + geometric filtering | $\sim 12,700$ | 500-1000 | High quality predicted structural ensemble |
| AF2-MSAss | Unfiltered MSA subsampling | $\sim 12,700$ | 1000 | Implausible predicted structures |

All structural ensemble analyses for real-world experiments for protein systems: BPTI, BRD4, HOIP, LXRa and MBP can be found under Ensemble Analyses in the Additional Experiment Plots. This includes; Principal Component Analysis (PCA), Ensemble PCA Comparison; local Cross Correlation (LCC), Ensemble LCC Comparison; as well as pairwise ensemble heatmaps for residue-level RMSD, and Wasserstein distances, Pairwise Ensemble Comparison. Exact de-tails of structural analysis and comparisons performed can be found in the Quantifying Ensem-ble Similarity and Conformational Coverage section.

#### 5.2.1 Iso-Validation Experiments

Simulation data for the Iso-Validation experiment was obtained from the original HDXer tutorial.[16] To produce the plots here we repeated the trajectory clustering on a sliced (1:50) subset of the original trajectory provided. Synthetic HDX-uptake curves were generated without added measurement noise to isolate the effects of ensemble composition from experimental uncertainty.

##### Trajectory Clustering

Trajectory clustering was performed by aligning all frames to each reference (Open and Closed) and computing the RMSD. Frames were sorted into Open and Closed clusters if structures were less than 1.0 Å from one of the reference structures.

##### Bi-Modal Ensemble

The Bi-Modal ensemble is formed by taking the structures directly from the clustering process. By artificially inverting (reweighting) the two major conformational states in this ensemble we produce the HDX uptake curves used for fitting.

##### Tri-Modal Ensemble

The Tri-Modal ensemble is the parent ensemble of the clustered, Bi-Modal ensemble. This contains Open, Closed as well as intermediate structures.

#### 5.2.2 AlphaFold Experiments

##### Featurisation of Ensembles

Since AF2 structures natively do not contain hydrogens, these must be added post-hoc. Backbone amide H atoms were added using standard peptide geometry (N–H bond length 1.02 Å).[44] Backbone amide protons coordinates were computed using barycentric coordinates of the peptide plane.

##### AlphaFold MSA Subsampling

Colabfold was used to generate the AlphaFold2 ensembles.[34] MSA Subsampling,[45] was performed over a range of 1-127 Max Sequences where Max Extra Sequences set to twice the Max Seq value. Each parameter set was used to generate 100 structures (20 seeds) For each protein, a single recycling step was used with no minimisation, generating 12700 structures in total.

##### AlphaFold Ensemble Cleaning

Since AF2 generates structures according to a statistical potential, this leads to the inclusion of unnatural/implausible structures in the ensemble. This is more prevalent at lower recycle counts. To quantify structure confidence, predicted Local Distance Difference Test (pLDDT) from AF2 is used in addition to predicted Template Modelling (pTM). While high structure quality (pLDDT and pTM) correlates well with crystal structures, the design of AF2 means that the generated ensemble is biased towards structures that exist within the training data. For large, flexible or disordered proteins this represents a problem as pLDDT no longer is able to provide a reliable estimate of structure quality. As a result, fixed confidence score cut-offs using AF2 predicted structure quality measures is not sufficient to clean ensembles *a priori*. Other works in the literature use system-specific collective variables to identify relevant conformations, however, this requires existing knowledge of the protein system which may not exist or introduce human bias of what structures *should* look like.

No widely adopted system-agnostic protocol is standard; we therefore developed an approach to filter ensembles using knowledge that exists for almost any protein. These filters are designed to handle some consistent AF2 failure modes, which include: over-folding, mis-folding and under-folding. We hypothesise that the prevalence of implausible structures is linked to the size, flexibility and availability of crystal structures. Our strategy involves the use of multiple, broad measures of protein structure to successively filter structures based on percentiles chosen based on rules defined by broad characteristics of the protein. Filtering uses only AF2/geometry-derived features and does not use HDX data, preventing circular validation.

The filtering pipeline proceeds in three conceptual steps:

1. **Remove misfolded outputs**: Structures were first filtered by AF2 predicted confidence metrics (pLDDT and pTM) using percentile-based thresholds specific to each protein system.
2. **Remove over-/mis-folding**: Structures were then filtered by contact order and local cross correlation (LCC) relative to the maximum pLDDT structure.
3. **Remove under-folded**: Finally, structures were filtered by inter-residue distances using the same maximum and minimum threshold for all proteins.

Validation of the filtering protocol was performed by clustering (k=20) and then comparing the LCC between cluster-centroid structures. All centroids across ensembles show a mean-LCC of at least 0.9. Figures demonstrating each stage of filtering can be found in AF2-MSAss ensemble cleaning using confidence-based quality assessment and AF2-MSAss ensemble cleaning using contact order and local correlation assessment, for pTM, and pLDDT; as well as Contact order, and local Correlation, respectively. Specific percentile parameters for each protein are provided in Parameters for AF2 Ensemble cleaning.

#### 5.2.3 MD Experiments

The purpose of the various MD experiments performed is to create a range of ensembles with distinct differences in their sampling properties. They are intended to represent broad regimes of approaches to MD and not exhaustive approaches to generating ideal ensembles. To ease comparison, experiments were designed to use a similar amount of compute to the AF2 ensembles for each protein (≈ 100 ns of simulations).

##### Simulation Setup

MD experiments were performed using GROMACS. Structures were protonated at pH 7 using GROMACS pdb2gmx with the AMBER ff14SB force field.[46] The system was solvated using the TIP3P water model. Forcefield parameters were chosen to match the original settings by the AMBER MD engine itself; simulation parameters and equilibration protocols were obtained from the repository supporting Kosarim et al.[47] Exact implementations used for this study can be found in the repositories in Code and Data Availability. Key simulation parameters are summarised in MD Simulation Parameters.

**Methods Table M4:** MD Simulation Parameters.

| Parameter | Value |
| --- | --- |
| Force field | AMBER ff14SB |
| Water model | TIP3P |
| protonation pH | 7.0 (GROMPP) |
| Salt concentration | 0.15 M NaCl |
| Thermostat | V-rescale |
| Barostat | Parrinello-Rahman |
| Temperature | 300 K |
| Pressure | 1 bar |
| Timestep | 2 fs |
| Electrostatics | PME |
| Cutoffs | 1.0 nm |
| Constraints | H-bonds (LINCS) |

##### Minimisation and Equilibration Protocol

After topology generation, restraints generated by the GROMACS Pre-Processor (GROMPP) were modified to only cover *C_α_* atoms from residues that are included by the experimental data. Position restraints on *C_α_* atoms were applied during equilibration to prevent large backbone collapse when starting from diverse AF2 geometries, then gradually released over five stages. Since the starting structures cover a wide range of shapes, topologies were generated for each structure separately.

**Methods Table M5:** MD Simulation protocol.

| Step | Length | Restraint Strength Kj/mol |
| --- | --- | --- |
| Minimisation | <10000 steps | 500 |
| EQ1 | 100 ps | 500 |
| EQ2 | 200 ps | 500 |
| EQ3 | 200 ps | 50 |
| EQ4 | 200 ps | 5 |
| EQ5 | 1 ns | 0 |

##### MD-1Start

We selected the single highest confidence AF2 structure and performed 5 replicates of 20 ns of production MD. This represents the standard approach of initiating simulation from a single, high-quality structure to test if conventional MD suffices to sample relevant states. To ensure the stability of the structures 10 ns of unrestrained MD was additionally performed before the start of the production step.

##### MD-10Start

To represent a more diverse approach to generating ensembles with respect to HDX-MS data, the filtered AF2 (AF2-Filtered) ensemble was used to generate representative structures identified through optimisation. k-means (k=10) clustering was used to generate diverse starting structures which was are then equilibrated using the same protocol as MD-1Start. To maintain a comparable computational budget to MD-1Start, each starting structures was run with a single replicate of 5 ns and 10 ns for relaxation and production simulations, respectively.

##### T-FES

To represent the most diverse range of fully plausible structures possible within a reasonable compute budget we modified an equilibrium-MD sampling approach, Frontier Expansion Sampling (FES).[48] We adjusted the algorithm to concurrently sample across diverse starting structures with different topologies multi-Topology Frontier Expansion Sampling (T-FES). To seed the initial conformations we combine both MD ensembles generated previously (MD-1Start and MD-10Start). We perform 20 cycles, capped at 200 short (0.1 ns) simulations per cycle which generates a maximum of 400 ns of short equilibrium-MD simulations. To maintain compatibility with the previous simulations the relevant GROMACS simulation files were converted into the, Open-MM compatible, Amber file formats using ParmED.[49] We retained OpenMM’s default Langevin integrator, but added a Monte-Carlo Barostat to match the Parrinello-Rahman barostat used in the GROMACS simulations. Each short simulation is preceded by an ultra-short equilibration (2000 steps, 1 fs time-step) to facilitate particle-velocity re-initialisation.

#### 5.2.4 Structure Poisoning Experiments

Structure poisoning experiments were designed to evaluate the robustness of ensemble optimisation methods by introducing controlled structural perturbations. These experiments serve as negative controls to assess whether optimisation algorithms can distinguish between physically plausible and artificially corrupted conformations. Three distinct poisoning strategies were implemented to test different aspects of structural integrity (Structure Poisoning Strategies). Detailed algorithms and perturbation procedures are provided in Structure Poisoning Algorithms.

**Methods Table M6:** Structure Poisoning Strategies.

| Poisoning Type | Physical Interpretation | Structures Added |
| --- | --- | --- |
| Gaussian Noise | Mimics missing unfolded/flexible states | 0.1%, 1%, 10%, 50% |
| Coordinate Mixing | Creates physically impossible structures to test detection | 0.1%, 1%, 10%, 50% |
| Proton Shuffling | Alters hydrogen bonding patterns while preserving heavy atoms | 0.1%, 1%, 10%, 50% |

##### Gaussian Noise

Gaussian noise poisoning introduces unrealistic, uncorrelated thermal fluctuations by adding residue-specific random perturbations to atomic coordinates. This method globally unfolds the structure while testing the algorithm’s sensitivity to coordinate precision. The noise magnitude is set to the natural variance observed for each residue across the parent ensemble, ensuring biologically relevant perturbation scales. In terms of protection factors, this has the effect of reducing the scale of computed backbone potentials. The Gaussian noise algorithm calculates residue-specific variance from the ensemble, then applies normally distributed coordinate perturbations scaled to match the observed fluctuations. This approach introduces coordinate perturbations scaled to observed ensemble variance while maintaining global structural topology.

##### Mix Coordinates

Coordinate mixing represents the a conceptually more severe structural perturbation by randomly swapping atomic positions within each frame. At the protection factor level, this preserves global structural density (HDX-uptake scale) but effectively smooths the protection factors. By testing ensembles against protection factor blurring, this should minimise scale differences of poisoned structures and so more directly probes the physicality of ensembles.

##### Shuffle Protons

Proton shuffling specifically targets backbone amide hydrogen atoms, which are the primary reporters in HDX-MS experiments. This method randomly reassigns hydrogen positions while maintaining the positions of all heavy atoms. Since HDX-MS protection factors depend critically on hydrogen bonding patterns and local environment accessibility, at the ensemble-level this is testing the sensitivity to new and unique conformations, at the same global density (HDX-uptake scale).

### 5.3 Quantifying Ensemble Similarity and Conformational Coverage

To evaluate whether generated ensembles capture biologically relevant dynamics, we employed a suite of structural analysis metrics that probe protein behavior at different scales. These metrics allow us to quantify: (1) the global diversity of sampled conformations, (2) the similarity between different ensemble generation protocols, and (3) the local flexibility and correlated motions of specific residue networks.

#### 5.3.1 Dimensionality Reduction

To visualise the high-dimensional conformational landscape covered by each ensemble, we performed Principal Component Analysis (PCA) on the pairwise *C_α_* distances of all structures, residues were filtered by HDX-MS peptide coverage. This projection reveals the major modes of structural variation and allows for comparison of sampling coverage, without needing a reference. For clustering and representative structure selection, we used k-means clustering on the first 10 principal components.

#### 5.3.2 Structural Comparison

##### Root-Mean-Square Deviation (RMSD)

To quantify the global structural difference between ensembles, we computed the Root-Mean-Square Deviation (RMSD) between their average structures. This metric provides a baseline assessment of whether two ensembles are centered on the same mean conformation.

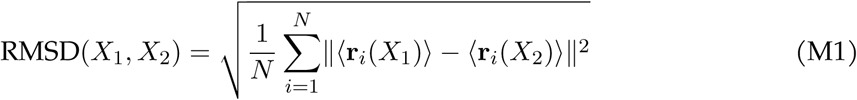

Where:

- ⟨r*_i_*(*X*)⟩ is the ensemble-averaged position of the *C_α_* atom of residue *i*
- *N* is the total number of residues
- Values nearing 0 Å indicate identical average structures
- Large values indicate systematic structural differences between the ensembles

For a more rigorous comparison that accounts for ensemble weights and avoids alignment artifacts, we also computed the pairwise ensemble RMSD using internal coordinates, detailed in Weighted RMSD.

##### Wasserstein Distance (Earth Mover’s Distance)

To quantify how different two ensembles appear at each residue position, we computed the Wasserstein-1 distance (*W*_1_). Unlike RMSD, which compares average structures, *W*_1_ captures the minimal “effort” required to morph one ensemble’s full conformational distribution into another. This provides a global measure of ensemble similarity that accounts for the shape and spread of conformational populations.

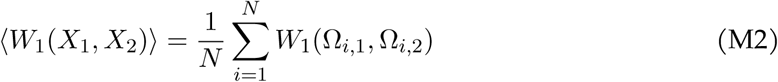

Where:

- *W*_1_ is the Wasserstein distance (units of Å)
- Ω*_i_* is the distribution of pairwise *C_α_* distances for residue *i*
- Values approaching 0 indicate identical conformational distributions
- Larger values indicate divergence in the sampled conformational landscapes

Extended mathematical details, including the Kernel Density Estimation (KDE) bandwidth selection and weighting schemes, are provided in Wasserstein Distance Details.

#### 5.3.3 Dynamics Comparison

##### Root Mean Square Fluctuation (RMSF)

To measure the local flexibility of the protein chain, we computed the Root Mean Square Fluctuation (RMSF). This metric quantifies how much each residue moves around its average position, allowing us to identify rigid domains versus flexible loops.

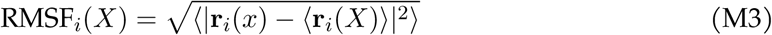

#### Where

- r*_i_*(*x*) is the position of residue *i* in structure *x*
- ⟨·⟩ denotes the ensemble average weighted by structure probabilities
- High values indicate flexible regions (e.g., loops, disordered tails)
- Low values indicate rigid regions (e.g., secondary structure elements, cores) The explicit formulation for weighted ensembles is provided in Weighted RMSF.

##### Local Cross Correlation (LCC)

To identify networks of residues that move together, we computed the Local Cross Correlation (LCC). This metric reveals correlated motions, helping to distinguish between independent flexible regions and rigid-body domain movements that may be functionally relevant for allostery.

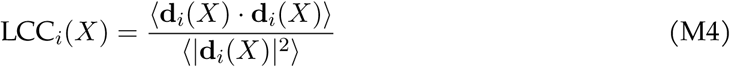

Where:

- d*_i_*(*x*) represents the vector of distances from residue *i* to all neighbors within 8 Å
- Values approaching 1 indicate highly correlated local motions (rigid-body behavior)
- Values near 0 indicate uncorrelated or anti-correlated dynamics
- The 8 Å cutoff captures immediate contacts and local packing environment

We also computed the Inter-Ensemble LCC to compare dynamical correlations across different ensembles; the mathematical definition is provided in Inter-Ensemble LCC.

### 5.4 Reweighting and Optimisation Methods

To align simulated ensembles with experimental HDX-MS data, we employ Maximum Entropy (MaxEnt) reweighting. This approach finds the minimal adjustment to ensemble populations required to match experimental observations, ensuring the result remains physically plausible while satisfying experimental constraints.

We employed three distinct optimisation protocols, differing in which parameters were allowed to vary:

**Methods Table M7:** Optimisation Protocols.

| Protocol | What Changes | What Stays Fixed |
| --- | --- | --- |
| **RW-only** | Frame weights | $\beta_C, \beta_H$ parameters |
| **BV-only** | $\beta_C, \beta_H$ parameters | Frame weights (uniform) |
| **Combined** | Both, in stages | — |

#### 5.4.1 HDXer Loss Function

HDXer implements a Maximum Entropy (MaxEnt) optimisation framework to reweight the simulated ensemble. This process begins by establishing a link between protein structure and experimental exchange rates.

**Step 1: Predicting Protection Factors (BV Model)** To bridge the gap between structural ensembles and experimental HDX data, HDXer employs the Best-Vendruscolo (BV) model. This phenomenological model predicts protection factors (*P_f_*) from protein coordinates by counting structural features that impede solvent exchange.

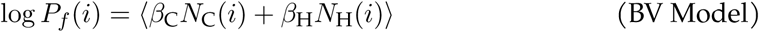

Where:

- *P_f_* (*i*) is the protection factor for residue *i* (dimensionless)
- *β*_C_ and *β*_H_ are empirical scaling coefficients (dimensionless) for contacts and hydrogen bonds
- *N*_C_(*i*) is the number of heavy-atom contacts within 6.5 Å of residue *i*
- *N*_H_(*i*) is the number of hydrogen bonds involving the amide of residue *i*
- ⟨·⟩ denotes the ensemble average

**Step 2: Defining the Ensemble Potential** We define a pseudo-energy for the ensemble based on these predicted protection factors. This energy represents the thermodynamic cost associated with the observed protection pattern.

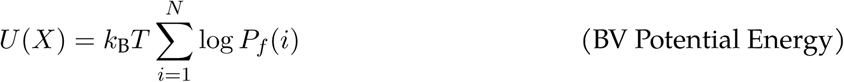

Where:

- *U* (*X*) is the potential energy of ensemble *X* (kJ/mol)
- *k*_B_ is the Boltzmann constant
- *T* is the temperature (Kelvin)

**Step 3: Reweighting the Ensemble** Reweighting the ensemble to match experimental data requires doing “work” against this potential. The corrected potential energy represents the energy of the reweighted ensemble.

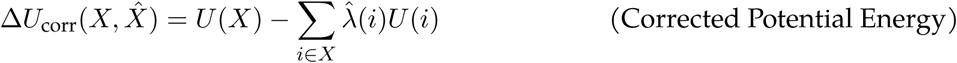

Where:

- Δ*U*_corr_ is the Apparent Work required to reweight the ensemble (kJ/mol)
- *λ*^^^(*i*) are the optimised Lagrangian multipliers for each residue *i*

**Step 4: The Optimisation Objective** The goal is to find the Lagrangian multipliers *λ*^^^(*i*) that minimize the total loss, which consists of the work done (keeping the ensemble close to the prior) and the error relative to experimental data.

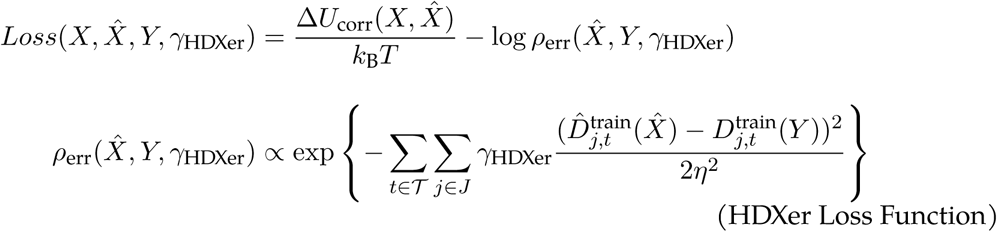

Where:

- Δ*U*_corr_(*X, X*^^^) is the corrected potential energy change (Apparent Work) required to reweight the ensemble (units of kJ/mol)
- *k*_B_*T* is the thermal energy
- *ρ*_err_ is the error density function quantifying agreement with experiment
- *γ*_HDXer_ is the tunable optimisation parameter (dimensionless) that balances fitting accuracy against the work required to reweight the ensemble
- *D*_*j,t*_^^train^(*X*^^^) is the predicted fractional uptake for peptide *j* at time *t*
- *D*_*j,t*_^train^(*Y*) is the experimental fractional uptake data
- *η* = 1 is the fixed uncertainty parameter

**Interpretation of** *γ***_HDXer_** Higher *γ*_HDXer_ values allow more aggressive fitting to experimental data but risk overfitting by forcing large deviations from the prior ensemble. Lower values preserve the prior distribution but may result in poorer fit to experiment. We tested *γ*_HDXer_ values from 1 × 10^0^ to 9 × 10^0^, spanning a range from minimal to strong fitting. Larger values of *γ*_HDXer_ were tested in preliminary analysies but we found that above *γ*_HDXer_ = 9 × 10^0^, the optimiser would consistently fail in some split-replicates.

### 5.5 Validation Framework

#### 5.5.1 ValDX Process Flow

Our overall validation methodology is described in ValDX Workflow Diagram, this section describes the exact implementation details of the ValDX workflow. ValDX is built around the HDXer optimisation package but the overall protocol is transferable.

The workflow starts with data splitting. For each replicate, ensemble-structural features are computed for both the training and validation peptide sets. The training data (peptides and features) are used for fitting across multiple free-parameter choices (*γ_HDXer_*). Within each split-replicate, representative parameters are selected using a parameter selection algorithm (curve-of-the-knee). From this replicate-parameter set, model-parameters and weights are extracted and used to calculate Validation MSE as well as Work Done distances. These metrics are pooled across replicates and can then be compared in downstream analysis.

##### Curve-of-the-Knee Optimiser

The optimal regularization parameter (*γ*_HDXer_) balances fitting accuracy against overfitting. We identified this balance point automatically by plotting apparent work against training error and finding the “elbow”, the point where additional fitting strength provides diminishing returns. This was computed as the point where the local gradient approaches −1 (45° slope) on the work-error tradeoff curve. The knee was selected from the training error versus apparent work curve (both linear scale), using the same *γ* grid (1 − 9 × 10^0^) across all experiments.

This is a naive implementation of the qualitative parameter selection method described by HDXer tutorials and literature.[13, 16] The full algorithm is provided in Curve-of-the-Knee *γ*_HDXer_ Optimisation in Supplementary Information.

#### 5.5.2 Practical Considerations

Due to the complexity of ensemble-data fitting there are numerous possible failure modes, however these can be addressed within each component. For data splitting, overlap can be resolved by adjusting initial peptide redundancy dropout or directly removing extremely long or redundant peptides. Optimisation runs themselves may fail, preliminary investigations showed that larger *γ_HDXer_* runs are more likely to fail; due to the Lagrangian formulation it is unlikely that all simulations within a replicate fail. If replicates are consistently failing, adding additional replicates can overcome this and lowering the scale for *γ_HDXer_*applied leads to more conservative fitting.

### 5.6 Metrics and Statistical Analysis

To quantify the quality of the optimised ensembles, we employed a set of metrics measuring both the fit to experimental data and the deviation from the prior ensemble (Work Done). Detailed thermodynamic derivations and statistical validation of these metrics are provided in the Supplementary Information (Derivation of Work Done Metrics and Metric Quality Quantification).

#### 5.6.1 Metrics at a Glance

**Methods Table M8:**
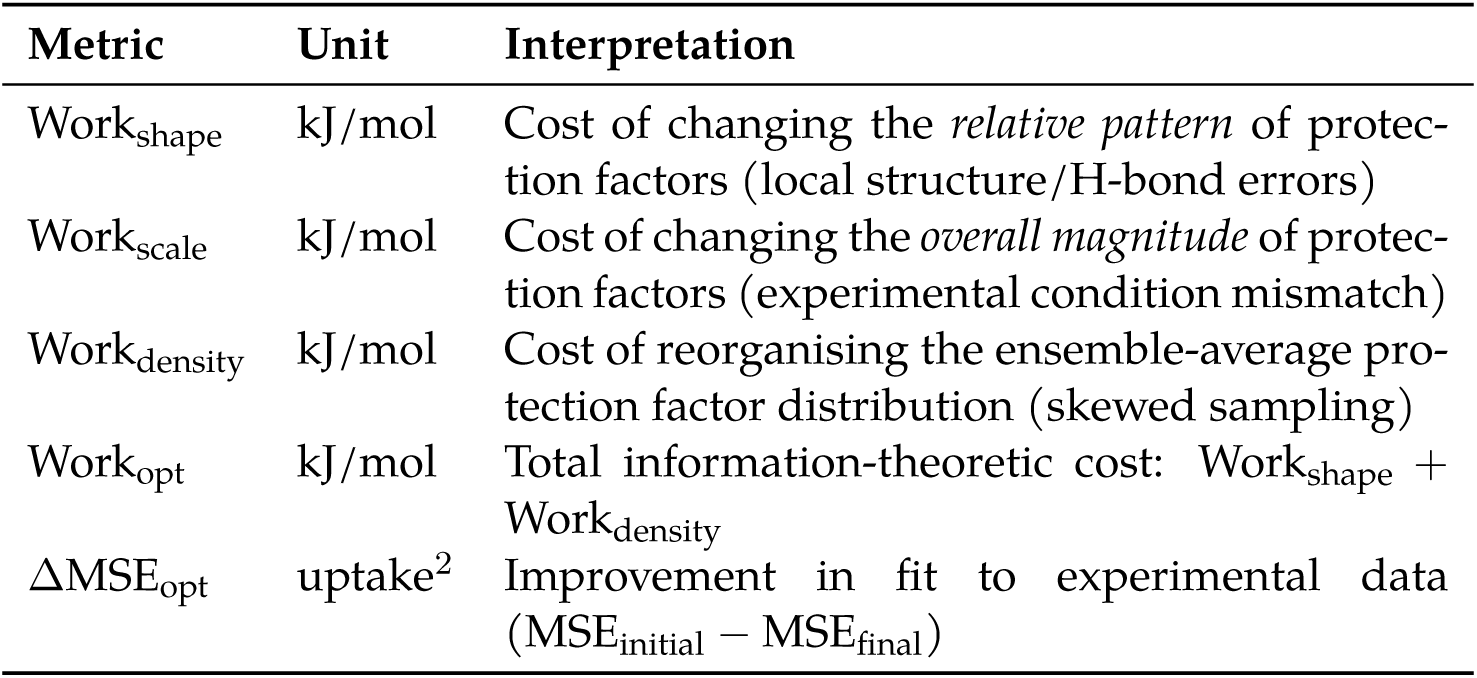
Summary of Optimisation Metrics.

#### 5.6.2 Work Done Metrics

We define “Work Done” as the information-theoretic cost of reweighting the ensemble away from its prior distribution. Unlike the apparent internal energy term reported by HDXer, which reflects only the regularised reweighting objective, our Work metrics quantify both magnitude shifts and distributional reorganisation in protection factor space.

##### Work Shape (Work_shape_)

To quantify how much the *relative pattern* of protection factors changes during optimisation, independent of global shifts in magnitude, we calculate Work_shape_. This metric captures the “structural” changes in the ensemble’s protection profile.

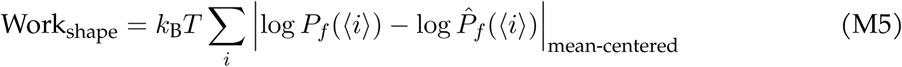

Where:

- *k*_B_*T* is the thermal energy (2.479 kJ/mol at 298 K)
- *P_f_* (⟨*i*⟩) is the initial protection factor of residue *i*
- *P*^^^*_f_* (⟨*i*⟩) is the optimised protection factor of residue *i*
- |*…*|_mean-centered_ denotes the absolute difference after mean-centering both profiles

##### Interpretation

Large values indicate that optimisation substantially reorganised which protein regions appear protected versus solvent-exposed, typically signifying incorrect local structure or erroneous hydrogen-bonding networks.

##### Work Scale (Work_scale_)

To measure the global shift in protection factor magnitude (e.g., the whole protein becoming more or less protected), we calculate Work_scale_.

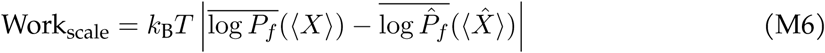

Where:

- log *P_f_* (⟨*X*⟩) is the mean log protection factor of the initial ensemble
- log *P*^^^*_f_* (⟨*X*^^^⟩) is the mean log protection factor of the optimised ensemble

##### Interpretation

Elevated values suggest systematic mismatch between experimental conditions and model calibration rather than structural deficiencies.

##### Work Density (Work_density_)

To quantify the reorganisation of ensemble-average protection factors during optimisation, we calculate Work_density_. This metric captures the entropic contribution to the Work Done.

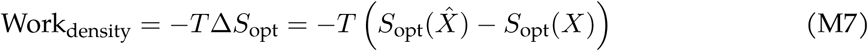

Where:

- *S*_opt_(*X*) is the thermodynamic entropy of the ensemble-average protection factor distribution of *X*
- *T* is the temperature

##### Interpretation

Large values indicate that the optimiser substantially shifted the distribution of protection factors, suggesting that the initial conformational sampling was skewed relative to the true solution-state populations or that important conformational states were inadequately sampled.

##### Total Work (Work_opt_)

The total information-theoretic cost of optimisation is the sum of the shape and density costs, where entropy is computed over the ensemble-average protection factor distribution across residues:

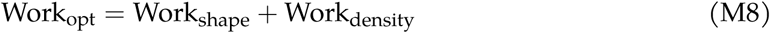

##### Interpretation

This represents the total information-theoretic cost of transforming the initial ensemble hypothesis into one that matches experimental observations.

#### 5.6.3 Operational Metrics

##### MSE Improvement (ΔMSE_opt_)

To quantify the improvement in fit to experimental data, we calculate the absolute reduction in Mean Squared Error.

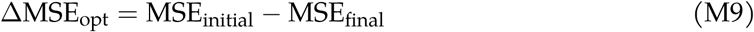

##### Interpretation

Higher values indicate a greater reduction in error when fitting the experimental HDX data.

##### Z-Score

To assess whether the improvement in fit is statistically significant relative to the work done, we compute a Z-score.

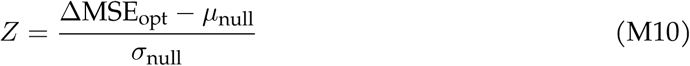

Where:

- *µ*_null_*, σ*_null_ are the mean and standard deviation of ΔMSE_opt_ from a null distribution (random reweighting)

#### 5.6.4 Statistical Analysis

Statistical significance of metric differences between ensemble generation methods was assessed using one-way ANOVA followed by Tukey’s HSD post-hoc tests for pairwise comparisons. Significance levels are reported as: † *p <* 0.25, ‡ *p <* 0.1, * *p <* 0.05, ** *p <* 0.01, *** *p <* 0.001, **** *p <* 0.0001. All statistical analyses were performed using the ‘scipy.stats‘ and ‘statsmodels‘ Python packages.

We also evaluated the predictive utility and stability of these metrics using multiple linear regression and variance decomposition (Partial *R*^2^, Stability Index, CV, *η*^2^). Detailed results of this validation are presented in Supplementary Information Metric Quality Quantification.

### 5.7 Case Study Protocols

#### 5.7.1 Iso-Validation Protocol

- **Inputs:** ISO-BiModal and ISO-TriModal ensembles; Synthetic HDX-uptake curves (generated from inverted ISO-BiModal ensemble, 5% to 40% Open).
- **Optimised parameters:** Frame weights (RW-only).
- **Splits/Replicates:** 3 replicates × 4 split types (Non-Redundant, Spatial, Random, Sequential).
- **Optimisation Grid:** *γ*_HDXer_ ∈ {1*, . . .,* 9} × 10^{−1^,^0^,^1}^.
- **Outputs computed:** MSE_Training_, Apparent Work, Open State Recovery %.

Simulation data was obtained from the HDXer tutorial.[16] Synthetic protection factors were computed using the Radou method (BV-model with switch function).[21]

#### 5.7.2 Reweighting-Only Protocol

- **Inputs:** Clustered ensembles (500 structures, k-means on PCA); HDX-MS Data (HDX-NMR for BPTI).
- **Optimised parameters:** Frame weights (RW-only).
- **Splits/Replicates:** 3 replicates × 2 split types (Non-Redundant, Spatial).
- **Optimisation Grid:** *γ*_HDXer_ ∈ {1*, . . .,* 8} × 10^0^.
- **Outputs computed:** *W*_1_ distance (vs. parent ensemble), MSE_Validation_, ΔMSE_opt_, Δ*H*_opt_, Multi-variate *σ* (using Δ*H*_abs_ and −*T* Δ*S*_opt_).

*W*_1_ distance was computed between the ensemble-average of the unclustered parent ensembles (AF2-Filtered and MD-1Start).

#### 5.7.3 BV-Model Parameter Optimisation Protocol

- **Inputs:** Unclustered parent ensembles; HDX-MS Data.
- Optimised parameters: *β_C_, β_H_* (BV-only).
- **Splits/Replicates:** 10 replicates × 2 split types (Non-Redundant, Spatial).
- **Outputs computed:** Optimal *β_C_, β_H_* values, MSE_Validation_.

Single structures for comparison were selected as: AF2-MaxRank (highest pLDDT from ColabFold default) and AF2-MaxPLDDT (highest pLDDT from entire AF2-MSAss ensemble).

#### 5.7.4 Protocol Comparison Benchmark

- **Inputs:** Ensembles; HDX-MS Data (HDX-NMR for BPTI).
- **Optimised parameters:** All protocols (BV-only, RW-only, RWafterBV, BVafterRW, MutualBVRW).
- **Splits/Replicates:** 3 replicates per stage.
- **Optimisation Grid:** *γ*_HDXer_ ∈ {1*, . . .,* 8} × 10^0^ (for reweighting stages).
- **Outputs computed:** Comparative metrics across protocols.

For multi-stage protocols, replicates were mean-averaged over weights and parameters to provide the prior for the next stage.

#### 5.7.5 Ensemble Size (Clustering) Sweep

- **Inputs:** Parent ensembles (10k-12.7k structures), clustered to sizes {0.1%, 1%, 5%, 10%, 20%, 50%, 100%}; HDX-MS Data (HDX-NMR for BPTI).
- **Optimised parameters:** Frame weights (RW-only).
- **Splits/Replicates:** 3 replicates × 2 split types.
- **Optimisation Grid:** *γ*_HDXer_ ∈ {1*, . . .,* 8} × 10^0^.
- **Outputs computed:** Metrics vs Cluster Size.

Clustering was performed via k-means on 10 PCA dimensions.

#### 5.7.6 Structure Poisoning Protocol

- **Inputs:** Clustered ensembles (1000 structures) + Poisoned structures (0.1%, 1%, 10%, 50%); HDX-MS Data.
- **Optimised parameters:** Frame weights (RW-only).
- **Splits/Replicates:** 3 replicates × 2 split types.
- **Optimisation Grid:** *γ*_HDXer_ ∈ {1*, . . .,* 8} × 10^0^.
- **Outputs computed:** Correlation (*R*) and Standard Deviation (*SD*) of metrics.

Poisoned structures were added sequentially, ordered by Euclidean distance (RMSD in PCA space) to the uncorrupted ensemble-average. The 50% poisoned ensemble served as a negative control.

### 5.8 Computational Implementation Details

All simulations were performed using the University of Oxford’s Advanced Research Cluster (ARC).[50] Traditional MD simulations used GROMACS 2021.3.[51] Enhanced sampling protocols (T-FES) were implemented in OpenMM 8.1.2.[52] AlphaFold2 structure predictions were generated using LocalColabFold (commit 1208ceb).[34] HDXer (commit f06191e) was used for all ensemble reweighting and optimisation.[16]

Key analysis libraries included MDAnalysis (2.6.1),[53] scikit-learn (1.5.2),[54] and SciPy (1.14.1).[55] Full computational environment details, including package versions and performance benchmarks for all simulation types, are provided in Computational Environment and Performance.

### 5.9 Code and Data Availability

To democratise adoption and reproducibility of quantitative HDX-MS workflows, we provide details and scripts for all major components in this work. This includes the ValDX workflow using a modified HDXer, associated plots and data processing scripts. The scripts for ensemble generation experiments are also provided, including: topology generation, colabfold simulations, AF2-ensemble cleaning scripts, and MD simulation files.

#### 5.9.1 HDXer Package

HDXer (version commit f06191 used) is a Python package for MaxENT optimisation for HDX-MS data. The package provides implementations of the Best-Vendruscolo model and frame reweighting optimisation algorithms used in this study. https://github.com/Lucy-Forrest-Lab/HDXer

#### 5.9.2 ValDX

The ValDX validation framework developed for this study was written in python, building on a modified version of the HDXer package to directly extract parameters. The workflow scripts and analysis notebooks are split into separate repositories.

**Workflow** ValDX workflow repository, implementing peptide splitting algorithms, optimisation protocols, and metrics calculation for ensemble validation. https://github.com/alexisiddiqui/ValDX

**Plots and Miscellaneous scripts** Python scripts for generating figures, statistical analyses, and ensemble/data processing reported in this work. https://github.com/alexisiddiqui/interpretable-hdxer

#### 5.9.3 Topology Generation

Notebooks to generate and validate topologies used for MD simulations for each of the starting structures used in MD simulations. This also repository contains the forcefield description files. https://github.com/alexisiddiqui/topology_generation

#### 5.9.4 Traditional MD Simulations

Traditional MD simulations were performed using GROMACS (2021.3) using topologies generated from AF2 structures.

**Simulation files and scripts** Traditional MD simulations were run with a small python package that acts as a wrapper for GROMACS, primarily to manage paths across repeated simu-lations. https://github.com/alexisiddiqui/xMD. Simulation parameters were obtained from https://github.com/intbio/gmx_protocols (commit d70eb3e), the equilibration protocols were modified from those provided in the *gmx_protocols* repository to separate position restraints on heavy atoms and backbone atoms across different stages of equilibration.

#### 5.9.5 Enhanced MD Simulations

Enhanced MD simulations were run using OpenMM (8.1.2), the T-FES protocol was built from Frontier-Expansion-Sampling,[48] modified for simulation correctness and performance.

**Frontier Expansion Sampling** A modified Frontier-Expansion-Sampling protocol was used to perform enhanced sampling MD simulations. These scripts do not correctly apply a barostat for pressure coupling. https://github.com/Gonglab-THU-MD/Frontier-Expansion-Sampling

**T-FES scripts** T-FES modifies the original Frontier-Expansion-Sampling workflow in a few key ways. Primarily, pressure coupling through a Monte-Carlo barostat for correctness and consistency to the traditional MD simulations. The protocol was also extended to simulate across multiple topologies, simultaneously and in parallel. https://github.com/alexisiddiqui/multi-Topology-Frontier-Expansion-Sampling

#### 5.9.6 AF2 Simulations

AF2 was used to generate initial ensembles of our protein systems, this was achieved using MSA subsampling.[34]

**LocalColabFold** LocalColabFold (version commit 1208ceb) was used to perform MSA sub-sampling. https://github.com/YoshitakaMo/localcolabfold

**MSA Subsampling workflow scripts** Simulation, initial analysis, and AF2 ensemble processing scripts for all protein systems in this study are provided here. https://github.com/alexisiddiqui/xFold_Sampling

### 5.10 Conflicts of Interest

The authors declare the following financial interest which may be considered as potential competing interest(s): Rachael Skyner, Maria Musgaard, and Srinath Krishnamurthy are share-holder of OMass Therapeutics. Srinath Krishnamurthy is furthermore an employee of OMass Therapeutics and Rachael Skyner is an employee of Kiin Bio.

#### 5.10.1 Statement on Language Model use

Large Language Models (From providers: Claude, Gemini, and DeepSeek) were used solely for improving the clarity of the manuscript. No generative AI tools were used to analyse, derive, or generate synthetic data. Authors claim complete ownership of the content of this work.

## References

1. Masson GR, Jenkins ML, Burke JE. An overview of hydrogen deuterium exchange mass spectrometry (HDX-MS) in drug discovery. Expert Opin Drug Discov 2017; 12:981–94. doi:10.1080/17460441.2017.1363734

2. Masson GR, Burke JE, Ahn NG, Anand GS, Borchers C, Brier S, et al. Recommendations for performing, interpreting and reporting hydrogen deuterium exchange mass spectrometry (HDX-MS) experiments. Nat Methods 2019; 16:595–602. doi:10.1038/s41592-019-0459-y

3. Narang D, Lento C, Wilson DJ. HDX-MS: An Analytical Tool to Capture Protein Motion in Action. Biomedicines 2020; 8:224. doi:10.3390/biomedicines8070224

4. James EI, Murphree TA, Vorauer C, Engen JR, Guttman M. Advances in Hydrogen/Deuterium Exchange Mass Spectrometry and the Pursuit of Challenging Biological Systems. Chem Rev 2022; 122:7562–623. doi:10.1021/acs.chemrev.1c00279

5. Marciano DP, Dharmarajan V, Griffin PR. HDX-MS guided drug discovery: small molecules and biopharmaceuticals. Curr Opin Struct Biol 2014; 28:105–11. doi:10.1016/j.sbi.2014.08.007

6. West GM, Chien EYT, Katritch V, Gatchalian J, Chalmers MJ, Stevens RC, et al. Ligand-Dependent Perturbation of the Conformational Ensemble for the GPCR *β*2 Adrenergic Receptor Revealed by HDX. Structure 2011; 19:1424–32. doi:10.1016/j.str.2011.08.001

7. Li S, Lee SY, Chung KY. Conformational analysis of g protein-coupled receptor signaling by hydrogen/deuterium exchange mass spectrometry. Methods Enzymol 2015; 557:261–78. doi:10.1016/bs.mie.2014.12.004

8. Ständer S, Grauslund LR, Scarselli M, Norais N, Rand K. Epitope Mapping of Polyclonal Antibodies by Hydrogen–Deuterium Exchange Mass Spectrometry (HDX-MS). Anal Chem 2021; 93:11669–78. doi:10.1021/acs.analchem.1c00696

9. Anderson KW, Bergonzo C, Scott K, Karageorgos IL, Gallagher ES, Tayi VS, et al. HDX-MS and MD Simulations Provide Evidence for Stabilization of the IgG1—Fc*γ*RIa (CD64a) Im-mune Complex Through Intermolecular Glycoprotein Bonds. J Mol Biol 2022; 434:167391. doi:10.1016/j.jmb.2021.167391

10. Lim XX, Chandramohan A, Lim XYE, Bag N, Sharma KK, Wirawan M, et al. Conforma-tional changes in intact dengue virus reveal serotype-specific expansion. Nat Commun 2017; 8:14339. doi:10.1038/ncomms14339

11. Hamuro Y, Derebe MG, Venkataramani S, Nemeth JF. The effects of intramolecular and intermolecular electrostatic repulsions on the stability and aggregation of NISTmAb re-vealed by HDX-MS, DSC, and nanoDSF. Protein Sci 2021; 30:1686–700. doi:10.1002/pro.4129

12. Mitra G. Emerging Role of Mass Spectrometry-Based Structural Proteomics in Elucidating Intrinsic Disorder in Proteins. Proteomics 2021; 21:2000011. doi:10.1002/pmic.202000011

13. Lee PS, Bradshaw RT, Marinelli F, Kihn K, Smith A, Wintrode PL, et al. Interpreting Hydrogen-Deuterium Exchange Experiments with Molecular Simulations: Tutorials and Applications of the HDXer Ensemble Reweighting Software [Article v1.0]. LiveCoMS 2021; 3:1521. doi:10.33011/livecoms.3.1.1521

14. Mohammadiarani H, Shaw VS, Neubig RR, Vashisth H. Interpreting Hydrogen-Deuterium Exchange Events in Proteins Using Atomistic Simulations: Case Studies on Regulators of G-protein Signaling Proteins. J Phys Chem B 2018; 122:9314–23. doi:10.1021/acs.jpcb.8b07494

15. Jia R, Bradshaw RT, Calvaresi V, Politis A. Integrating Hydrogen Deuterium Exchange–Mass Spectrometry with Molecular Simulations Enables Quantification of the Conformational Populations of the Sugar Transporter XylE. J Am Chem Soc 2023; 145:7768–79. doi:10.1021/jacs.2c06148

16. Bradshaw RT, Marinelli F, Faraldo-Gómez JD, Forrest LR. Interpretation of HDX Data by Maximum-Entropy Reweighting of Simulated Structural Ensembles. Biophys J 2020; 118:1649–64. doi:10.1016/j.bpj.2020.02.005

17. Kihn KC, Wilson T, Smith AK, Bradshaw RT, Wintrode PL, Forrest LR, et al. Modeling the native ensemble of PhuS using enhanced sampling MD and HDX-ensemble reweighting. Biophys J 2021; 120:5141–57. doi:10.1016/j.bpj.2021.11.010

18. Puchades C, Kűkrer B, Diefenbach O, Sneekes-Vriese E, Juraszek J, Koudstaal W, et al. Epitope mapping of diverse influenza Hemagglutinin drug candidates using HDX-MS. Sci Rep 2019; 9:4735. doi:10.1038/s41598-019-41179-0

19. Wan H, Ge Y, Razavi A, Voelz VA. Reconciling Simulated Ensembles of Apomyoglobin with Experimental Hydrogen/Deuterium Exchange Data Using Bayesian Inference and Multiensemble Markov State Models. J Chem Theory Comput 2020; 16:1333–48. doi:10.1021/acs.jctc.9b01240

20. Jia R, Martens C, Shekhar M, Pant S, Pellowe GA, Lau AM, et al. Hydrogen-deuterium exchange mass spectrometry captures distinct dynamics upon substrate and inhibitor bind-ing to a transporter. Nat Commun 2020; 11:6162. doi:10.1038/s41467-020-20032-3

21. Skinner SP, Radou G, Tuma R, Houwing-Duistermaat JJ, Paci E. Estimating Constraints for Protection Factors from HDX-MS Data. Biophys J 2019; 116:1194–203. doi:10.1016/j.bpj.2019.02.024

22. Sørensen L, Salbo R. Optimized Workflow for Selecting Peptides for HDX-MS Data Analyses. J Am Soc Mass Spectrom 2018; 29:2278–81. doi:10.1007/s13361-018-2056-1

23. Wollenberg DTW, Pengelley S, Mouritsen JC, Suckau D, Jørgensen CI, Jørgensen TJD. Avoiding H/D Scrambling with Minimal Ion Transmission Loss for HDX-MS/MS-ETD Analysis on a High-Resolution Q-TOF Mass Spectrometer. Anal Chem 2020; 92:7453–61. doi:10.1021/acs.analchem.9b05208

24. Fang M, Wang Z, Cupp-Sutton KA, Welborn T, Smith K, Wu S. High-throughput hydrogen deuterium exchange mass spectrometry (HDX-MS) coupled with subzero-temperature ultrahigh pressure liquid chromatography (UPLC) separation for complex sample analy-sis. Anal Chim Acta 2021; 1143:65–72. doi:10.1016/j.aca.2020.11.022

25. Hummer G, Köfinger J. Bayesian ensemble refinement by replica simulations and reweight-ing. J Chem Phys 2015; 143:243150. doi:10.1063/1.4937786

26. Köfinger J, Stelzl LS, Reuter K, Allande C, Reichel K, Hummer G. Efficient Ensemble Re-finement by Reweighting. J Chem Theory Comput 2019; 15:3390–401. doi:10.1021/acs.jctc.8b01231

27. Cesari A, Reißer S, Bussi G. Using the Maximum Entropy Principle to Combine Simula-tions and Solution Experiments. Computation 2018; 6:15. doi:10.3390/computation6010015

28. Mousa R, Lansky S, Shoham G, Metanis N. BPTI folding revisited: switching a disulfide into methylene thioacetal reveals a previously hidden path. Chem Sci 2018; 9:4814–20. doi:10.1039/C8SC01110A

29. Burley SK, Berman HM, Kleywegt GJ, Markley JL, Nakamura H, Velankar S. Protein Data Bank (PDB): The Single Global Macromolecular Structure Archive. In: Wlodawer A, Dauter Z, Jaskolski M, eds. Protein Crystallography: Methods and Protocols. New York. Springer. 2017. pp. 627–41. doi:10.1007/978-1-4939-7000-1_26

30. Globisch C, Krishnamani V, Deserno M, Peter C. Optimization of an Elastic Network Aug-mented Coarse Grained Model to Study CCMV Capsid Deformation. PLoS One 2013; 8:e60582. doi:10.1371/journal.pone.0060582

31. Bender BJ, Gahbauer S, Luttens A, Lyu J, Webb CM, Stein RM, et al. A practical guide to large-scale docking. Nat Protoc 2021; 16:4799–832. doi:10.1038/s41596-021-00597-z

32. Bonomi M, Bussi G, Camilloni C, Tribello GA, Banáš P, Barducci A, et al. Promoting transparency and reproducibility in enhanced molecular simulations. Nat Methods 2019; 16:670–3. doi:10.1038/s41592-019-0506-8

33. Burendahl S, Nilsson L. Computational studies of LXR molecular interactions reveal an allosteric communication pathway. Proteins 2012; 80:294–306. doi:10.1002/prot.23209

34. Mirdita M, Schütze K, Moriwaki Y, Heo L, Ovchinnikov S, Steinegger M. ColabFold: making protein folding accessible to all. Nat Methods 2022; 19:679–82. doi:10.1038/s41592-022-01488-1

35. Zakharova E, Horvath MP, Goldenberg DP. Functional and Structural Roles of the Cys14-Cys38 Disulfide of Bovine Pancreatic Trypsin Inhibitor. J Mol Biol 2008; 382:998–1013. doi:10.1016/j.jmb.2008.07.063

36. Qin M, Wang W, Thirumalai D. Protein folding guides disulfide bond formation. Proc Natl Acad Sci U S A 2015; 112:11241–6. doi:10.1073/pnas.1503909112

37. Shaw DE, Maragakis P, Lindorff-Larsen K, Piana S, Dror RO, Eastwood MP, et al. Atomic-Level Characterization of the Structural Dynamics of Proteins. Science 2010; 330:341–6. doi:10.1126/science.1187409

38. Tsai YCI, Johansson H, Dixon D, Martin S, Chung Cw, Clarkson J, et al. Single-Domain Antibodies as Crystallization Chaperones to Enable Structure-Based Inhibitor Development for RBR E3 Ubiquitin Ligases. Cell Chem Biol 2020; 27:83–93.e9. doi:10.1016/j.chembiol.2019.11.007

39. Crook OM, Gittens N, Chung Cw, Deane CM. A Functional Bayesian Model for Hydrogen–Deuterium Exchange Mass Spectrometry. J Proteome Res 2023; 22:2959–72. doi:10.1021/acs.jproteome.3c00297

40. Hageman TS, Weis DD. Reliable Identification of Significant Differences in Differential Hydrogen Exchange-Mass Spectrometry Measurements Using a Hybrid Significance Testing Approach. Anal Chem 2019; 91:8008–16. doi:10.1021/acs.analchem.9b01325

41. Belorusova AY, Evertsson E, Hovdal D, Sandmark J, Bratt E, Maxvall I, et al. Structural analysis identifies an escape route from the adverse lipogenic effects of liver X receptor ligands. Commun Biol 2019; 2:431. doi:10.1038/s42003-019-0675-0

42. Best RB, Vendruscolo M. Structural Interpretation of Hydrogen Exchange Protection Fac-tors in Proteins: Characterization of the Native State Fluctuations of CI2. Structure 2006; 14:97–106. doi:10.1016/j.str.2005.09.012

43. Hudgens JW, Gallagher ES, Karageorgos I, Anderson KW, Filliben JJ, Huang RYC, et al. Interlaboratory Comparison of Hydrogen-Deuterium Exchange Mass Spectrometry Measurements of the Fab Fragment of NISTmAb. Anal Chem 2019; 91:7336–45. doi:10.1021/acs.analchem.9b01100

44. Yao L, Vögeli B, Ying J, Bax A. NMR Determination of Amide N-H Equilibrium Bond Length from Concerted Dipolar Coupling Measurements. J Am Chem Soc 2008; 130:16518–20. doi:10.1021/ja805654f

45. Monteiro da Silva G, Cui JY, Dalgarno DC, Lisi GP, Rubenstein BM. High-throughput prediction of protein conformational distributions with subsampled AlphaFold2. Nat Com-mun 2024; 15:2464. doi:10.1038/s41467-024-46715-9

46. Maier JA, Martinez C, Kasavajhala K, Wickstrom L, Hauser KE, Simmerling C. ff14SB: Improving the Accuracy of Protein Side Chain and Backbone Parameters from ff99SB. J Chem Theory Comput 2015; 11:3696–713. doi:10.1021/acs.jctc.5b00255

47. Kosarim NA, Fedulova AS, Shariafetdinova AS, Armeev GA, Shaytan AK. Molecular Dynamics Simulations of Nucleosomes Containing Histone Variant H2A.J. Int J Mol Sci 2024; 25. doi:10.3390/ijms252212136

48. Zhang J, Gong H. Frontier Expansion Sampling: A Method to Accelerate Conformational Search by Identifying Novel Seed Structures for Restart. J Chem Theory Comput 2020; 16:4813–21. doi:10.1021/acs.jctc.0c00064

49. Shirts MR, Klein C, Swails JM, Yin J, Gilson MK, Mobley DL, et al. Lessons Learned from Comparing Molecular Dynamics Engines on the SAMPL5 Dataset. J Comput Aided Mol Des 2017; 31:147–61. doi:10.1007/s10822-016-9977-1

50. Richards A. University of Oxford Advanced Research Computing. 2015. doi:10.5281/zenodo.22558

51. Abraham MJ, Murtola T, Schulz R, Páll S, Smith JC, Hess B, et al. GROMACS: High performance molecular simulations through multi-level parallelism from laptops to supercomputers. SoftwareX 2015; 1-2:19–25. doi:10.1016/j.softx.2015.06.001

52. Eastman P, Swails J, Chodera JD, McGibbon RT, Zhao Y, Beauchamp KA, et al. OpenMM 7: Rapid development of high performance algorithms for molecular dynamics. PLoS Com-put Biol 2017; 13:e1005659. doi:10.1371/journal.pcbi.1005659

53. Gowers R, Linke M, Barnoud J, Reddy T, Melo M, Seyler S, et al. MDAnalysis: A Python Package for the Rapid Analysis of Molecular Dynamics Simulations. In: Proceedings of the 15th Python in Science Conference. pp. 98–105. doi:10.25080/Majora-629e541a-00e

54. Pedregosa F, Varoquaux G, Gramfort A, Michel V, Thirion B, Grisel O, et al. Scikit-learn: Machine Learning in Python. J Mach Learn Res 2011; 12:2825–30. doi:10.5555/1953048.2078195

55. Virtanen P, Gommers R, Oliphant TE, Haberland M, Reddy T, Cournapeau D, et al. SciPy 1.0: Fundamental Algorithms for Scientific Computing in Python. Nat Methods 2020; 17:261–72. doi:10.1038/s41592-019-0686-2

56. Jumper J, Evans R, Pritzel A, Green T, Figurnov M, Ronneberger O, et al. Highly accu-rate protein structure prediction with AlphaFold. Nature 2021; 596:583–9. doi:10.1038/s41586-021-03819-2

