## Supplementary manuscript for "A framework for testing structural hypotheses of protein dynamics against experimental HDX-MS data"

#### S0.1 Additional Experiment Plots

##### S0.1.1 Iso Validation Experiment

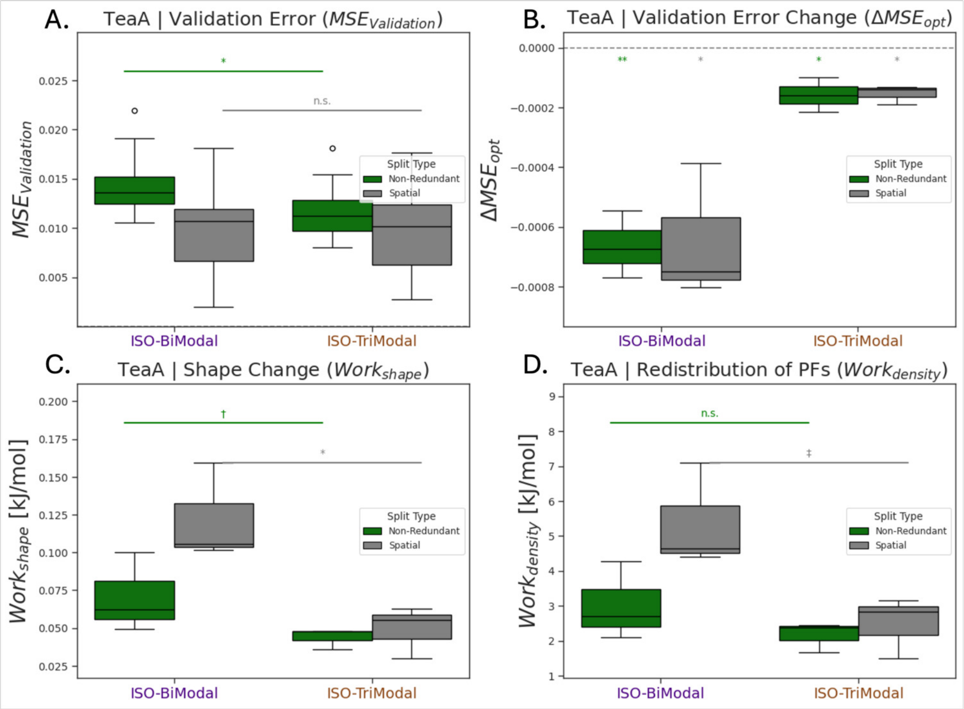


Supplementary Figure S1: Iso-Validation benchmark Work Done metrics demonstrate ensemble-integration assessment capabilities for TeaA membrane transporter synthetic experiment. Comparison between ISO-BiModal (indigo, clustered ensemble) and ISO-TriModal (gold, parent unclustered ensemble) using Non-Redundant (green) and Spatial (grey) splits across multiple replicates. **(A)** Validation error ($\mathrm{MSE}_{\mathrm{Validation}}$) shows significant separation between ensembles in Non-Redundant splits ($p = 2.02 \times 10^{-2}$) but not in Spatial splits, with ISO-BiModal demonstrating lower validation error. **(B)** Validation error change upon optimisation ($\Delta\mathrm{MSE}_{\mathrm{opt}}$) reveals both ensembles show similar improvement patterns with significant differences in Non-Redundant splits ($p = 2.29 \times 10^{-3}$). **(C)** Enthalpy of scale change ($\mathrm{Work}_{\mathrm{scale}}$) demonstrates no significant differences between ensembles in either split type, suggesting similar overall scale adjustments during optimisation. **(D)** Protection factor reorganisation requirements ($\mathrm{Work}_{\mathrm{density}}$) shows significant separation in Spatial splits ($p = 4.48 \times 10^{-2}$) with ISO-BiModal requiring less protection factor reorganisation, indicating better local protection factor estimation. Results support that Work Done validation metrics can differentiate ensemble quality using the complete peptide distribution, providing more realistic assessment than training-set-only Apparent Work metrics. The synthetic ground-truth (population ratio 60:40 open:closed) enables validation of the ValDX workflow's ability to assess ensemble-integration success beyond traditional error-based approaches.

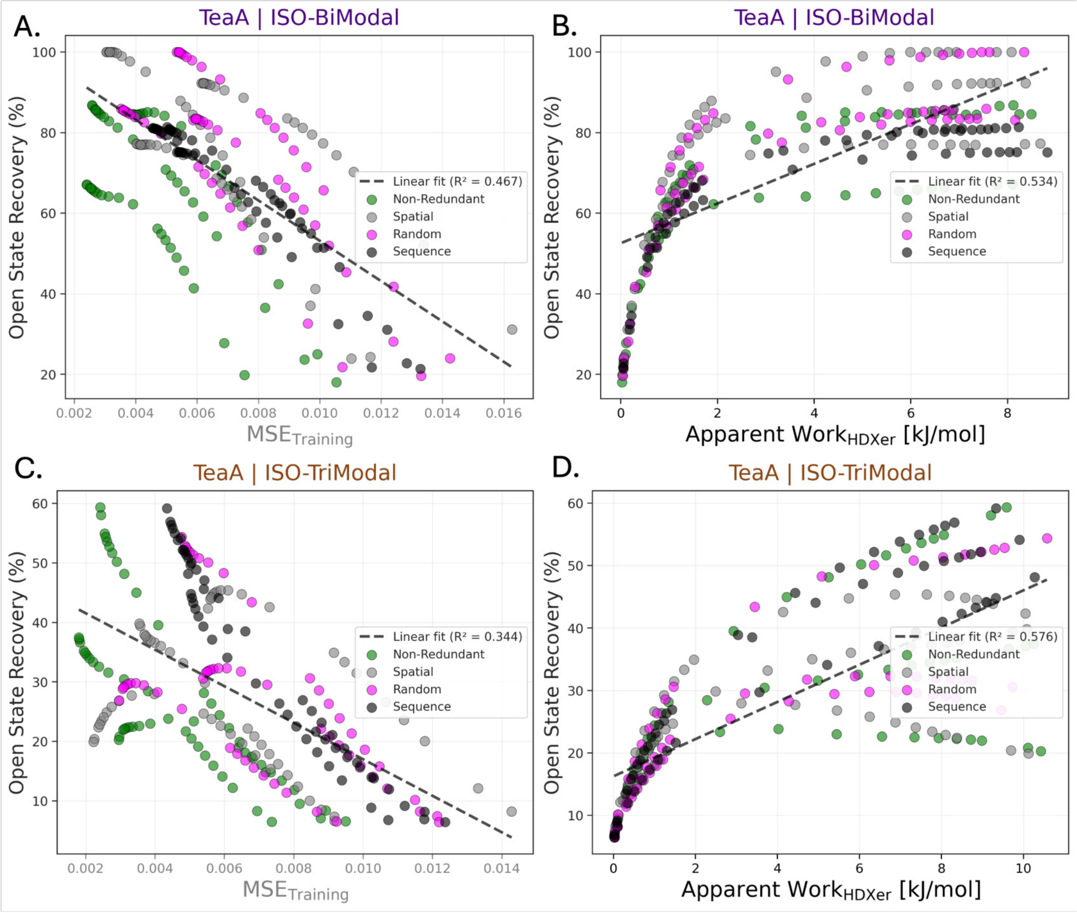


Supplementary Figure S2: Relationships between the two validation metrics ($MSE_{Training}$, Apparent $Work_{HDXer}$) and ground-truth, Open State Recovery (%) across split strategies for the TeaA Iso-Validation experiment. Points are individual replicates coloured by split-type (Non-Redundant, Spatial, Random, Sequence); dashed lines show least-squares linear fits with $R^2$ reported in each panel. **(A,C)** $MSE_{Training}$ is negatively associated with recovery and exhibits strong split-dependent stratification (ISO-BiModal $R^2 = 0.467$; ISO-TriModal $R^2 = 0.344$). **(B,D)** Apparent $Work_{HDXer}$ increases with recovery and collapses split-type variability onto a common monotonic trend, providing the stronger association in both ensembles (ISO-BiModal $R^2 = 0.534$; ISO-TriModal $R^2 = 0.576$). Apparent $Work_{HDXer}$ shows better linear relationship to Open State Recovery than $MSE_{Training}$, the gap between both metrics widens when considering the ISO-TriModal ensemble which contains non ground-truth, intermediate structures.

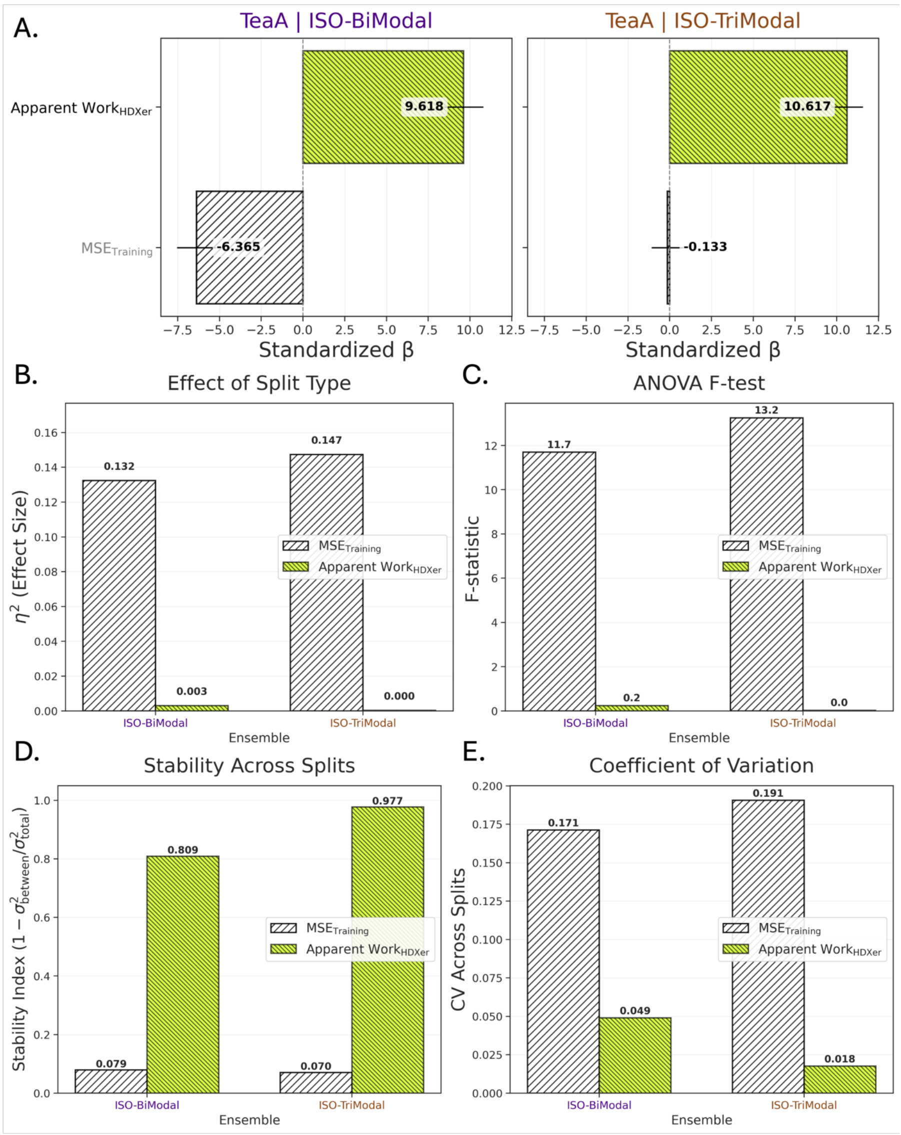


Supplementary Figure S3: Comparison of how the two surrogate metrics, MSE$_{\text{Training}}$ (forward-hatched) and Apparent Work$_{\text{HDXer}}$ (backward-hatched, neon), behave across split strategies in the TeaA Iso-Validation experiment. (**A**) Standardised regression coefficients ($\theta$) from a multivariable model relating recovery outcomes to the two proxies across all splits and replicates. Apparent Work$_{\text{HDXer}}$ is a strong positive predictor (ISO-BiModal: $\theta = 9.618$; ISO-TriModal: $\theta = 10.617$), whereas MSE$_{\text{Training}}$ contributes negatively and weakly (ISO-BiModal: $\theta = -6.365$; ISO-TriModal: $\theta = -0.133$). (**B**) Effect of split type ($\eta^2$): MSE$_{\text{Training}}$ shows a sizeable split effect ($0.132$–$0.147$) while Apparent Work$_{\text{HDXer}}$ is nearly unaffected ($0.003$–$0.000$). (**C**) One-way ANOVA $F$-statistics mirror this, with large $F$ for MSE$_{\text{Training}}$ ($11.7$–$13.2$) and negligible $F$ for Apparent Work$_{\text{HDXer}}$ ($0.2$–$0.0$). (**D**) Stability across splits quantified as $1 - \sigma^2_{\text{between}}/\sigma^2_{\text{total}}$: Apparent Work$_{\text{HDXer}}$ is highly stable ($0.809$–$0.977$) whereas MSE$_{\text{Training}}$ is unstable ($0.079$–$0.070$). (**E**) Coefficient of variation (CV) across splits is lower for Apparent Work$_{\text{HDXer}}$ ($0.049$–$0.018$) than for MSE$_{\text{Training}}$ ($0.171$–$0.191$). Together, these analyses show that Apparent Work$_{\text{HDXer}}$ is a robust, split-insensitive metric that better aligns with ground-truth performance, while MSE$_{\text{Training}}$ is more sensitive to how the data are split and is therefore less reliable for model selection; patterns are consistent for both ISO-BiModal and ISO-TriModal ensembles.

##### S0.1.2 RW-Only Experiment

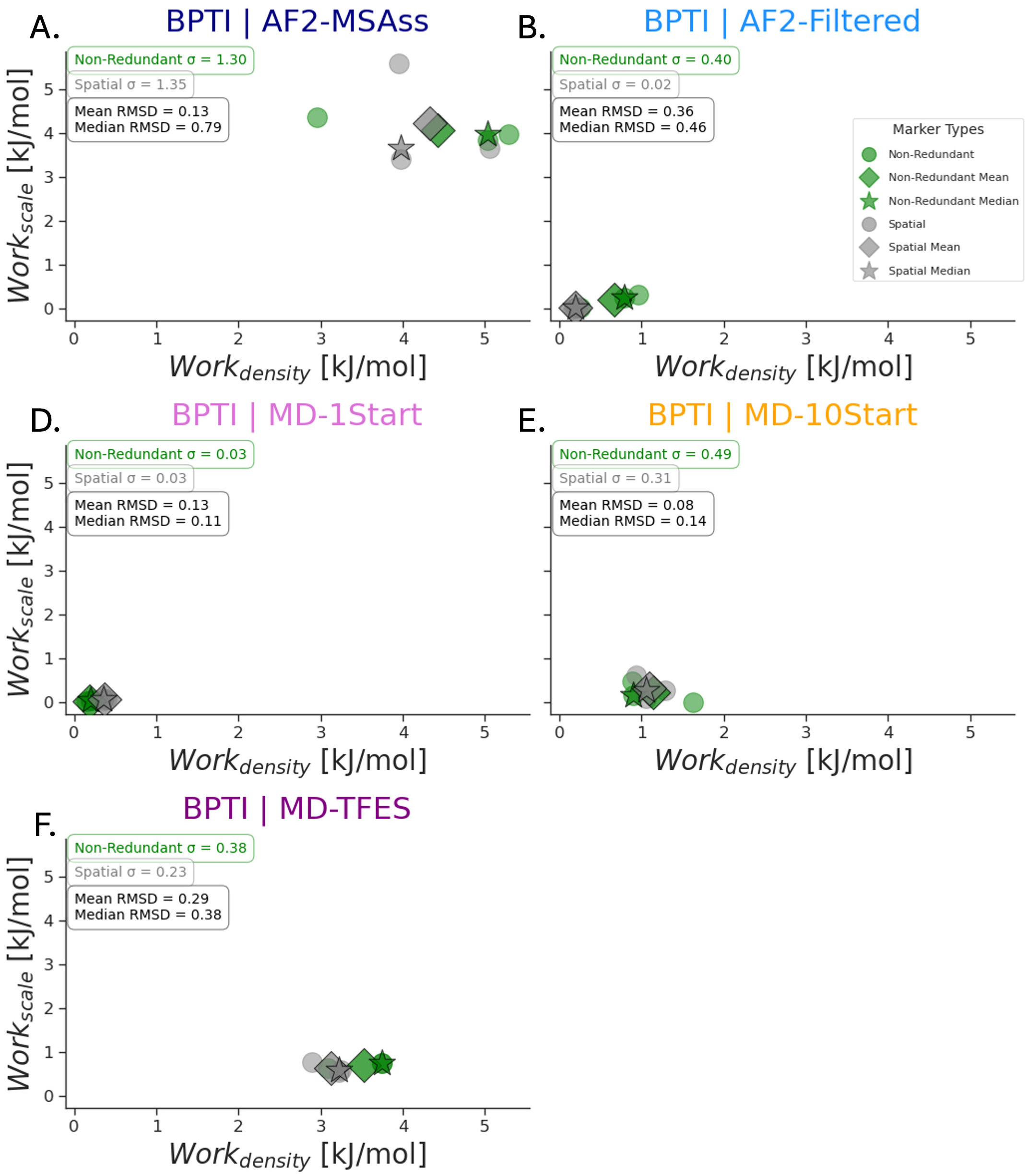


Supplementary Figure S4: Ensemble confidence assessment through Work Done metric variability across BPTI hierarchical ensembles. Scatter plots of absolute enthalpy change ($\text{Work}_{\text{scale}}$) versus entropy change during optimisation ($\text{Work}_{\text{density}}$) for **(A)** AF2-MSAss, **(B)** AF2-Filtered, **(C)** MD-1Start, **(D)** MD-10Start, and **(E)** MD-TFES ensemble experiments. Individual replicates shown as circles for Non-Redundant (green) and Spatial (grey) splits, with corresponding mean (diamonds) and median (stars) values. Multivariate Standard deviation ($\sigma$) values indicate ensemble confidence, with lower values representing higher confidence in optimisation outcomes. Mean and median RMSD values characterise split-agreement within each ensemble. MD-1Start (C) demonstrates both high representativeness (small changes upon optimisation) and high confidence (low $\sigma = 0.03$ for both splits), while AF2-MSAss (A) shows poor performance with high variability ($\sigma$ = 1.30-1.35). AF2-Filtered (B) exhibits intermediate confidence with better performance in Spatial splits ($\sigma = 0.02$) compared to Non-Redundant splits ($\sigma = 0.40$). Results support the use of multivariate standard deviation as a metric for ensemble quality assessment, enabling quantitative comparison of structural hypothesis reliability.

#### Supplementary Table S1: Statistical test results for BPTI analysis panels C-F

| Panel | Metric | Test Type | Comparison | p-value |
|---|---|---|---|---|
| C | $\text{MSE}_{\text{validation}}$ | 2-sample t-test | AF2-Filtered vs MD-1Start (Spatial) | $9.75 \times 10^{-1}$ |
| | | 2-sample t-test | AF2-Filtered vs MD-1Start (Non-Redundant) | $7.65 \times 10^{-1}$ |
| D | $\Delta\text{MSE}_{\text{opt}}$ | 1-sample t-test | AF2-Filtered, Non-Redundant vs 0 | $8.24 \times 10^{-1}$ |
| | | 1-sample t-test | AF2-Filtered, Spatial vs 0 | $1.77 \times 10^{-1}$ |
| | | 1-sample t-test | MD-1Start, Non-Redundant vs 0 | $1.70 \times 10^{-1}$ |
| | | 1-sample t-test | MD-1Start, Spatial vs 0 | $2.26 \times 10^{-1}$ |
| E | $\text{Work}_{\text{shape}}$ $(\Delta H_{\text{opt}})$ | 1-sample t-test | Spatial vs 0 | $5.17 \times 10^{-2}$ |
| | | 1-sample t-test | Non-Redundant vs 0 | $2.49 \times 10^{-1}$ |
| F | $\text{Work}_{\text{density}}$ $(-T\Delta S_{\text{opt}})$ | 1-sample t-test | Spatial vs 0 | $6.78 \times 10^{-4}$ |
| | | 1-sample t-test | Non-Redundant vs 0 | $1.50 \times 10^{-1}$ |

##### S0.1.3 BV-Only Experiment

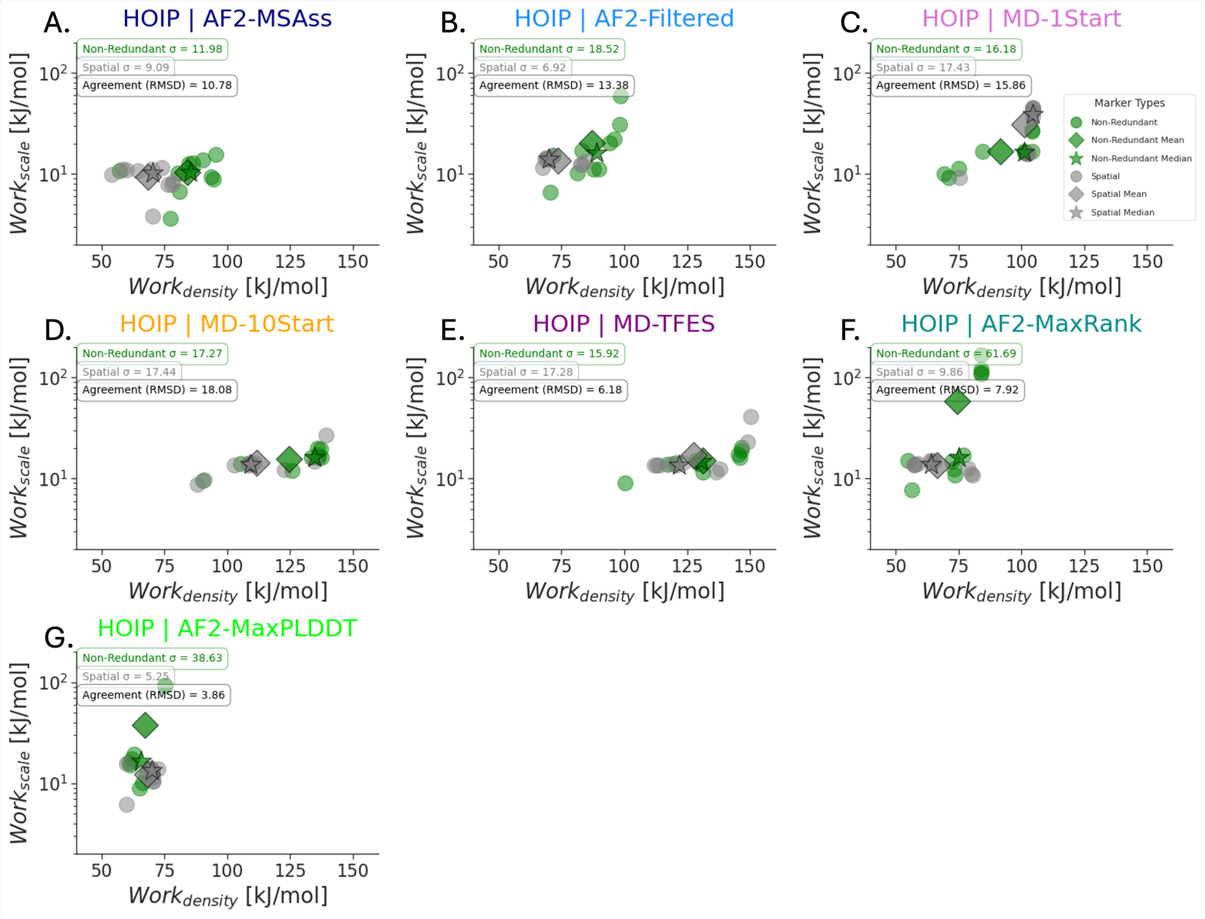


Supplementary Figure S5: BV-only optimisation confidence assessment through Work Done metric variability across HOIP structural hypotheses. Scatter plots of absolute enthalpy change ($\text{Work}_{\text{scale}}$) versus entropy change during optimisation ($\text{Work}_{\text{density}}$) for **(A)** AF2-MSAss, **(B)** AF2-Filtered, **(C)** MD-1Start, **(D)** MD-10Start, **(E)** MD-TFES, **(F)** AF2-MaxRank, and **(G)** AF2-MaxPLDDT structural hypotheses. Individual replicates shown as circles for Non-Redundant (green) and Spatial (grey) splits, with corresponding mean (diamonds) and median (stars) values. Multivariate standard deviation ($\sigma$) values indicate ensemble confidence, with lower values representing higher confidence in optimisation outcomes. Agreement (RMSD) quantifies concordance between split-types using median values, where lower RMSD indicates better agreement between global and local structural assessments. AF2-MaxPLDDT **(G)** demonstrates best data-split agreement (RMSD = 3.86) and highest confidence in Spatial splits ($\sigma$ = 5.25), while AF2-MSAss **(A)** shows misleadingly high confidence in Non-Redundant splits ($\sigma$ = 11.98) despite containing implausible structures. MD-TFES **(E)** exhibits strong confidence in both splits relative to other valid ensembles, supporting its role as the most diverse conformationally valid sampling strategy for the flexible HOIP system. Results demonstrate that unconstrained BV-model parameter optimisation requires careful interpretation, with replicates and data-split agreement providing essential uncertainty quantification for distinguishing valid from invalid structural hypotheses.

##### S0.1.4 Detailed Ensemble Confidence Assessment

This section extends the model parameter optimisation analysis presented in the main text by providing detailed confidence assessments across all HOIP structural hypotheses (Figure S5).

**Ensemble-level confidence patterns.** AF2-Filtered (Figure S5.B) exhibited the second highest local confidence ($\sigma = 6.92$) but the lowest global confidence ($\sigma = 18.52$), indicating reliable local structure representation but poor global conformational coverage. Notably, AF2-MSAss (Figure S5.A) reported the highest confidence in the Non-Redundant split ($\sigma = 11.98$), demonstrating that model parameter optimisation cannot detect individual structure validity since fitting operates at the ensemble-average level.

**Individual structure assessment.** AF2-MaxPLDDT (Figure S5.G) demonstrated the best data-split agreement (3.86 kJ/mol) overall, with the highest Spatial confidence ($\sigma = 5.25$) but second lowest Non-Redundant confidence ($\sigma = 38.63$). This pattern suggests a well-folded structure that partially represents the conformational landscape but cannot capture the full extent of global dynamics. AF2-MaxRank (Figure S5.F) showed better agreement (7.92 kJ/mol) than most ensembles despite containing only a single structure, though extremely high $\text{Work}_{\text{scale}}$ in some Non-Redundant replicates indicated large magnitude adjustments are required to fit the global conformation.

**MD ensemble comparisons.** MD-1Start (Figure S5.C) was the only ensemble favouring the Non-Redundant split, though with mediocre agreement (15.86 kJ/mol) and poor confidence ($\sigma = 16.18$ to $17.43$). The increased $\text{Work}_{\text{density}}$ for MD-10Start (Figure S5.D) relative to MD-1Start confirms that AF2-MaxPLDDT, used as the single starting structure for MD-1Start, represents the more dominant conformation. MD-10Start showed similar confidence ($\sigma = 17.43$ to $17.44$) to MD-1Start but lower $\text{Work}_{\text{scale}}$, suggesting both ensembles are similarly plausible despite MD-10Start's additional diversity. However, poor agreement for MD-10Start (18.08 kJ/-mol) indicates that conformational density estimates remain inadequate.

**Enhanced sampling performance.** MD-TFES (Figure S5.E) demonstrated the best split-type agreement amongst ensembles (6.18 kJ/mol) with strong confidence in both splits ($\sigma = 15.92$ to $17.28$) relative to other valid ensembles. Despite showing the largest $\text{Work}_{\text{density}}$ across all hypotheses, its superior agreement and confidence support its role as the most representative structural hypothesis for the flexible HOIP system. This contrasts with MD-10Start, which exhibited similar $\text{Work}_{\text{scale}}$ but substantially worse agreement and confidence, suggesting that the enhanced sampling strategy of MD-TFES provides genuinely improved conformational coverage rather than merely increased structural diversity.

##### S0.1.5 Fitting Protocol Experiment

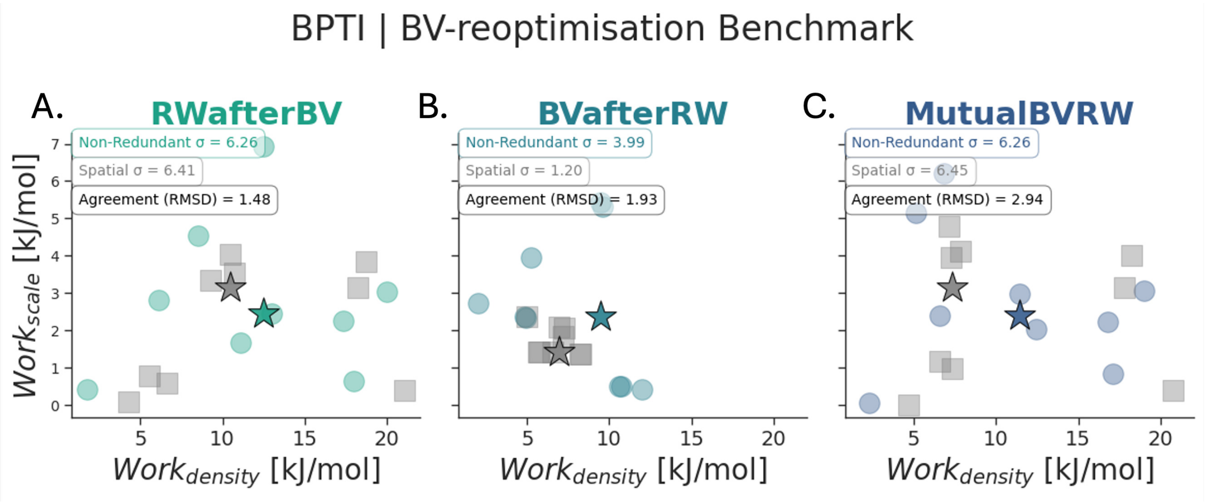


Supplementary Figure S6: BV-model parameter re-optimisation benchmark demonstrates protocol robustness differences across BPTI optimisation protocols. Scatter plots of absolute enthalpy change ($\Delta H_{\text{abs}}$) versus entropy change during optimisation ($-T\Delta S_{\text{opt}}$) following BV-model parameter re-optimisation for **(A)** RWafterBV (teal), **(B)** BVafterRW (blue), and **(C)** MutualBVRW (dark blue) protocols. Individual replicates shown as circles for Non-Redundant (coloured by protocol) and Spatial (grey) splits, with corresponding mean (squares) and median (stars) values. Multivariate standard deviation ($\sigma$) values indicate protocol confidence, while Agreement (RMSD) quantifies concordance between split-types using median values. BVafterRW **(B)** demonstrates superior performance with lower median entropy change for Non-Redundant splits, lower scale change in Spatial splits, and substantially higher confidence in both splits ($\sigma$ = 3.99 Non-Redundant, 1.20 Spatial) compared to RWafterBV **(A)** and MutualBVRW **(C)** which show similar uncertainty levels ($\sigma$ = 6.26-6.45). RWafterBV shows slightly better data-split agreement (RMSD = 1.48) but worse median performance, suggesting BV-model parameter smoothing at global (Non-Redundant split) and local (Spatial split) levels. Results confirm that forward model parameters significantly affect ensemble generalizability. The BVafterRW protocol produces the most robust ensemble-parameter set. Other protocols lead astray even when ensemble representativeness is already high.

Supplementary Table S2: Statistical test results for fitting protocols analysis panels A-C

| Panel | Metric | Test Type | Method | p-value |
|---|---|---|---|---|
| | | 1-sample t-test | BV-Only vs 0 | $5.73 \times 10^{-1}$ |
| | | 1-sample t-test | RW-Only vs 0 | $8.31 \times 10^{-2}$ |
| A | $\Delta\text{MSE}_{\text{opt}}$ | 1-sample t-test | RWafterBV vs 0 | $7.30 \times 10^{-1}$ |
| | | 1-sample t-test | BVafterRW vs 0 | $9.07 \times 10^{-1}$ |
| | | 1-sample t-test | MutualBVRW vs 0 | $5.00 \times 10^{-1}$ |
| | | 1-sample t-test | BV-Only vs 0 | $1.16 \times 10^{-2}$ |
| | | 1-sample t-test | RW-Only vs 0 | $9.83 \times 10^{-2}$ |
| | | 1-sample t-test | RWafterBV vs 0 | $7.53 \times 10^{-3}$ |
| B | $\text{Work}_{\text{opt}}$ | 1-sample t-test | BVafterRW vs 0 | $7.16 \times 10^{-2}$ |
| | | 1-sample t-test | MutualBVRW vs 0 | $1.13 \times 10^{-2}$ |
| | | 1-sample t-test | RWafterBV* vs 0 | $9.45 \times 10^{-3}$ |
| | | 1-sample t-test | BVafterRW* vs 0 | $8.51 \times 10^{-2}$ |
| | | 1-sample t-test | BV-Only vs 0 | $1.58 \times 10^{-1}$ |
| | | 1-sample t-test | RW-Only vs 0 | $1.55 \times 10^{-1}$ |
| | | 1-sample t-test | RWafterBV vs 0 | $1.85 \times 10^{-1}$ |
| C | $\text{Work}_{\text{scale}}$ | 1-sample t-test | BVafterRW vs 0 | $8.19 \times 10^{-2}$ |
| | | 1-sample t-test | MutualBVRW vs 0 | $1.51 \times 10^{-1}$ |
| | | 1-sample t-test | RWafterBV* vs 0 | $1.68 \times 10^{-1}$ |
| | | 1-sample t-test | BVafterRW* vs 0 | $3.29 \times 10^{-4}$ |

Supplementary Table S3: Standard deviations and correlations for BRD4 perturbation analysis

| Perturbation | Metric | Panel | Ensemble | SD | R |
|---|---|---|---|---|---|
| Gaussian Noise | $\Delta MSE_{opt}$ | B | AF2-MSAss | $6.10 \times 10^{-1}$ | $-8.60 \times 10^{-1}$ |
| | | | AF2-Filtered | $1.30 \times 10^{-1}$ | $-1.00 \times 10^{0}$ |
| | | | MD-1Start | $1.60 \times 10^{-1}$ | $-1.00 \times 10^{0}$ |
| | | | MD-10Start | $1.60 \times 10^{-1}$ | $-1.00 \times 10^{0}$ |
| | | | MD-TFES | $1.60 \times 10^{-1}$ | $-1.00 \times 10^{0}$ |
| | $Work_{opt}$ | C | AF2-MSAss | $4.30 \times 10^{-1}$ | $9.80 \times 10^{-1}$ |
| | | | AF2-Filtered | $2.10 \times 10^{-1}$ | $9.80 \times 10^{-1}$ |
| | | | MD-1Start | $1.30 \times 10^{-1}$ | $1.00 \times 10^{0}$ |
| | | | MD-10Start | $1.10 \times 10^{-1}$ | $9.90 \times 10^{-1}$ |
| | | | MD-TFES | $1.20 \times 10^{-1}$ | $1.00 \times 10^{0}$ |
| Mix Coordinates | $\Delta MSE_{opt}$ | E | AF2-MSAss | $2.40 \times 10^{-1}$ | $-4.80 \times 10^{-1}$ |
| | | | AF2-Filtered | $1.20 \times 10^{-1}$ | $9.30 \times 10^{-1}$ |
| | | | MD-1Start | $4.80 \times 10^{-1}$ | $4.40 \times 10^{-1}$ |
| | | | MD-10Start | $2.60 \times 10^{-1}$ | $7.20 \times 10^{-1}$ |
| | | | MD-TFES | $1.60 \times 10^{-1}$ | $9.70 \times 10^{-1}$ |
| | $Work_{opt}$ | F | AF2-MSAss | $2.60 \times 10^{-1}$ | $-4.00 \times 10^{-2}$ |
| | | | AF2-Filtered | $1.40 \times 10^{-1}$ | $5.60 \times 10^{-1}$ |
| | | | MD-1Start | $5.10 \times 10^{-1}$ | $3.00 \times 10^{-1}$ |
| | | | MD-10Start | $3.10 \times 10^{-1}$ | $7.00 \times 10^{-2}$ |
| | | | MD-TFES | $2.50 \times 10^{-1}$ | $4.40 \times 10^{-1}$ |
| Shuffle Protons | $\Delta MSE_{opt}$ | H | AF2-MSAss | $2.00 \times 10^{-1}$ | $-1.60 \times 10^{-1}$ |
| | | | AF2-Filtered | $7.00 \times 10^{-1}$ | $8.20 \times 10^{-1}$ |
| | | | MD-1Start | $6.00 \times 10^{-2}$ | $9.80 \times 10^{-1}$ |
| | | | MD-10Start | $5.10 \times 10^{-1}$ | $8.50 \times 10^{-1}$ |
| | | | MD-TFES | $1.10 \times 10^{-1}$ | $2.30 \times 10^{-1}$ |
| | $Work_{opt}$ | I | AF2-MSAss | $1.90 \times 10^{-1}$ | $-1.20 \times 10^{-1}$ |
| | | | AF2-Filtered | $4.70 \times 10^{-1}$ | $-9.70 \times 10^{-1}$ |
| | | | MD-1Start | $1.20 \times 10^{-1}$ | $7.00 \times 10^{-1}$ |
| | | | MD-10Start | $6.20 \times 10^{-1}$ | $-7.30 \times 10^{-1}$ |
| | | | MD-TFES | $1.60 \times 10^{-1}$ | $-5.80 \times 10^{-1}$ |

###### S0.1.6 Structure Poisoning Experiment

#### S0.2 Ensemble Analyses

###### S0.2.1 AF2-MSAss Ensemble Cleaning

###### S0.2.2 pTM and pLDDT Thresholding

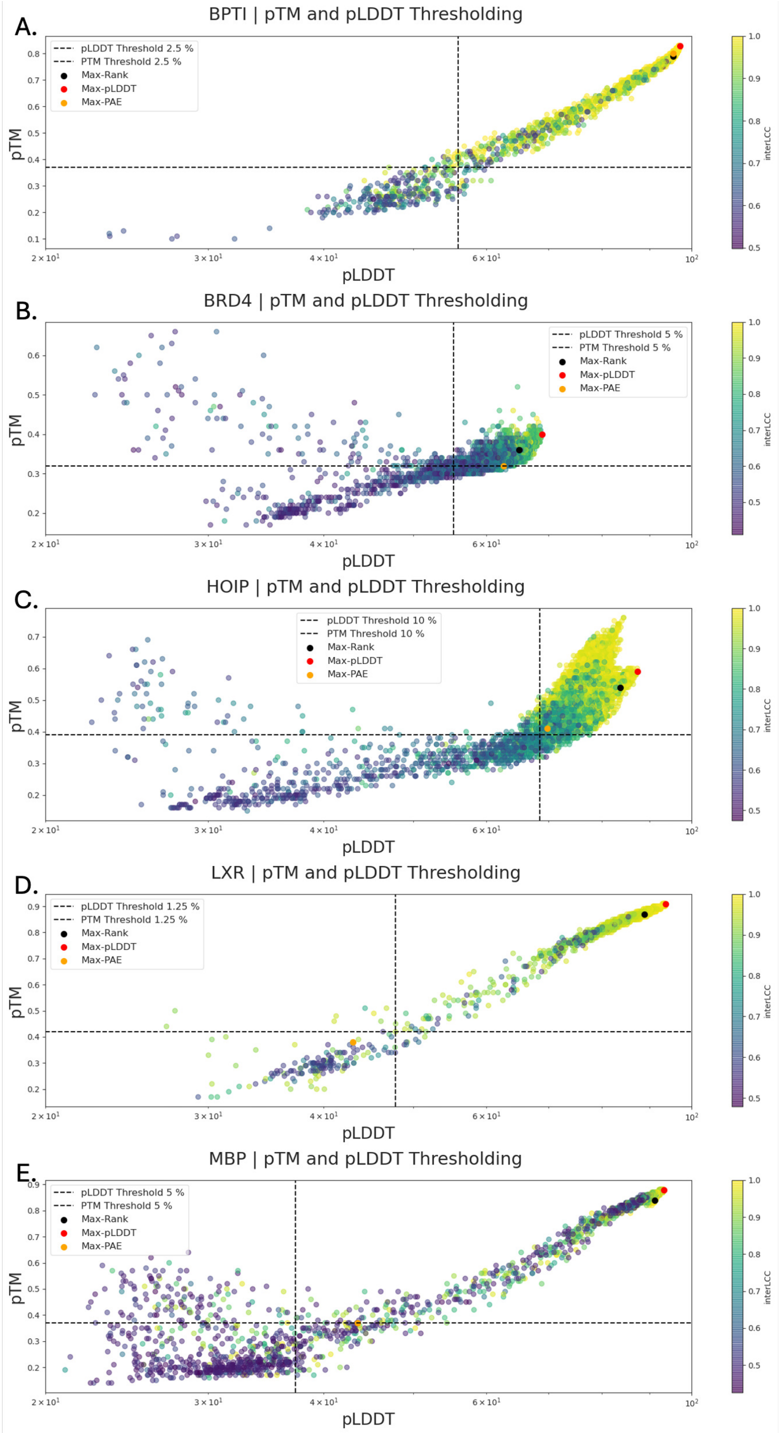


Supplementary Figure S7: AF2-MSAss ensemble cleaning using confidence-based quality assessment. Predicted Template Modelling (pTM) scores plotted against predicted Local Distance Difference Test (pLDDT) values for (**A**) BPTI, (**B**) BRD4, (**C**) HOIP, (**D**) LXR, and (**E**) MBP protein systems, with points coloured by local correlation to highest pLDDT structure (Max-pLDDT). Confidence thresholds for structure filtering shown as dashed black lines, with threshold percentages set *a priori*. Median values are shown by the dotted lines. Greatest scoring structures from different regimes labelled: Max-Rank (black crosses), Max-pLDDT (red stars), and Max-PAE (orange triangles) representing structures: Highest pLDDT structure from sub-ensemble 127-MSA, highest pLDDT from all structures 1-127 MSA (complete), and highest predicted Absolute Error (PAE) from all structures 1-127 MSA (complete), respectively.

##### S0.2.3 Contact Order and Local Correlation Thresholding

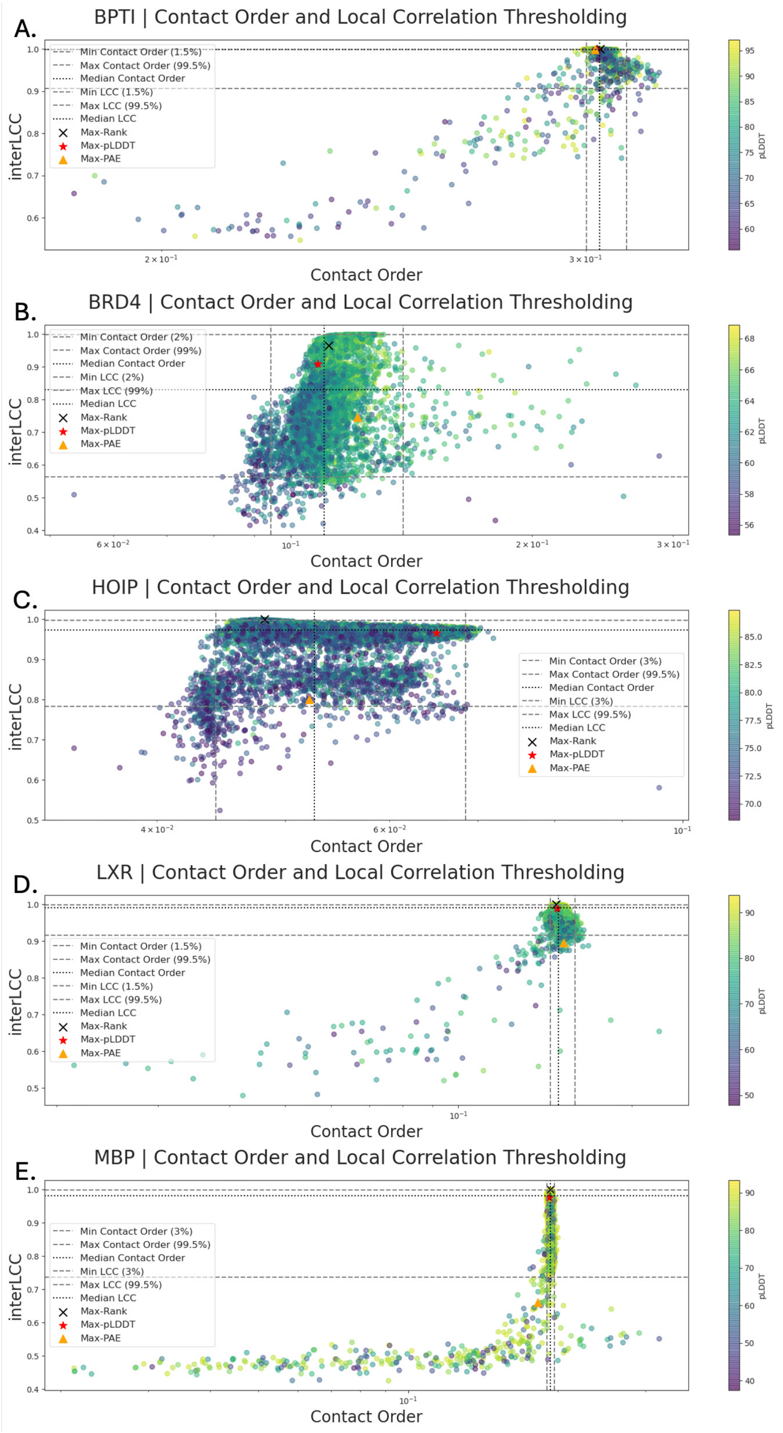


Supplementary Figure S8: AF2-MSAss ensemble cleaning using contact order and local correlation assessment across five protein systems. Contact Order plotted against inter-LCC (Local Cross Correlation) for (**A**) BPTI, (**B**) BRD4, (**C**) HOIP, (**D**) LXR, and (**E**) MBP protein systems, with points coloured by pLDDT confidence scores. Quality thresholds for structure filtering shown as grey dashed lines for minimum/maximum Contact Order and Local Correlation Coefficient, with percentile values varying by protein complexity. Median values indicated by black dotted lines. Representative structures labelled: Max-Rank (black crosses), Max-pLDDT (red stars), and Max-PAE (orange triangles) corresponding to highest pLDDT structure from 127-MSA subset, highest pLDDT from complete 1-127 MSA ensemble, and highest predicted Absolute Error structure from complete ensemble, respectively. Results demonstrate systematic structural quality gradients, with well-folded proteins showing distinct high-confidence clusters while flexible proteins display broader distributions.

##### S0.2.4 Ensemble PCA Comparison

##### S0.2.5 BPTI

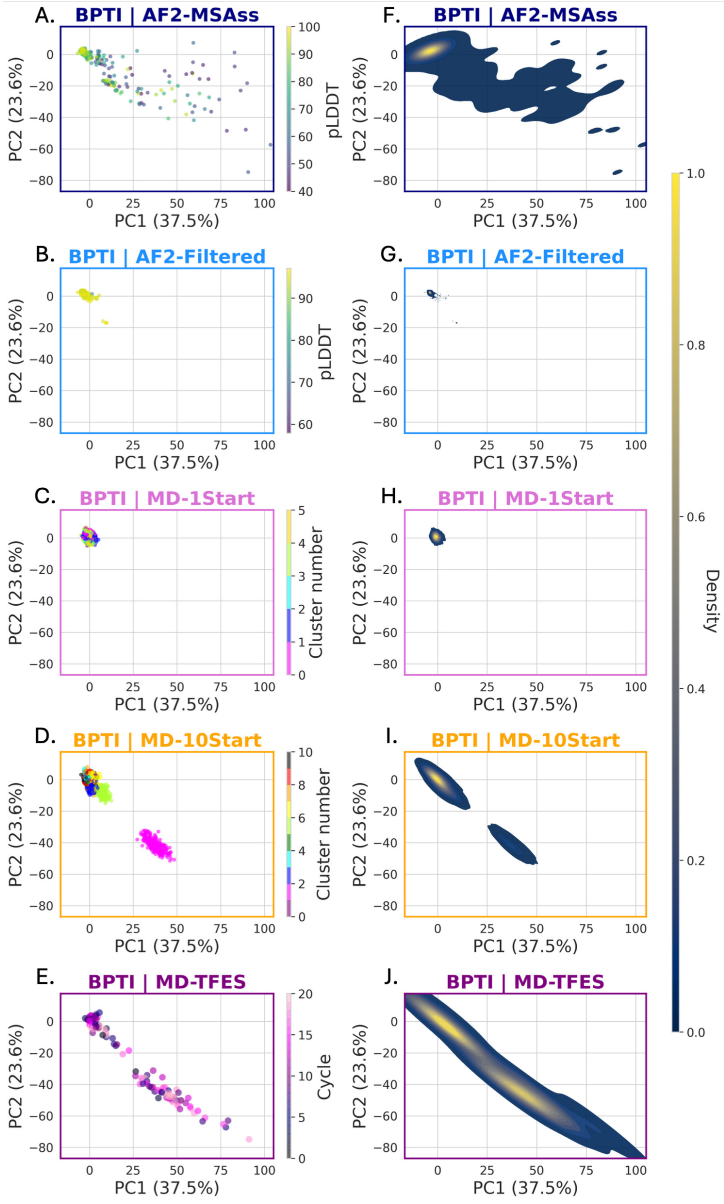


Supplementary Figure S9: BPTI ensemble comparison through PCA reveals distinct conformational landscapes across hierarchically sampled generation protocols. Principal component analysis scatter plots (**A-E**) and corresponding density distributions (**F-J**) for five BPTI ensembles: (**A,F**) AF2-MSAss coloured by pLDDT confidence scores, (**B,G**) AF2-Filtered, (**C,H**) MD-1Start coloured by cluster assignment, (**D,I**) MD-10Start showing multiple conformational clusters, and (**E,J**) MD-TFES displaying broad conformational sampling. Left panels show individual structure distributions in PC1-PC2 space, while right panels display density contours revealing population distributions. PCA was computed across all ensembles, MD-TFES was k-means clustered to 500 structures to better highlight the differences between cycles. For this ensemble, each point therefore represents a individual sampling distribution rather than a single structure. Results demonstrate progressive conformational diversity from filtered AF2 predictions through equilibrium MD to enhanced sampling, with AF2-MSAss showing scattered low-confidence structures, MD simulations exhibiting compact clusters, and MD-TFES achieving comprehensive landscape coverage.

##### S0.2.6 BRD4

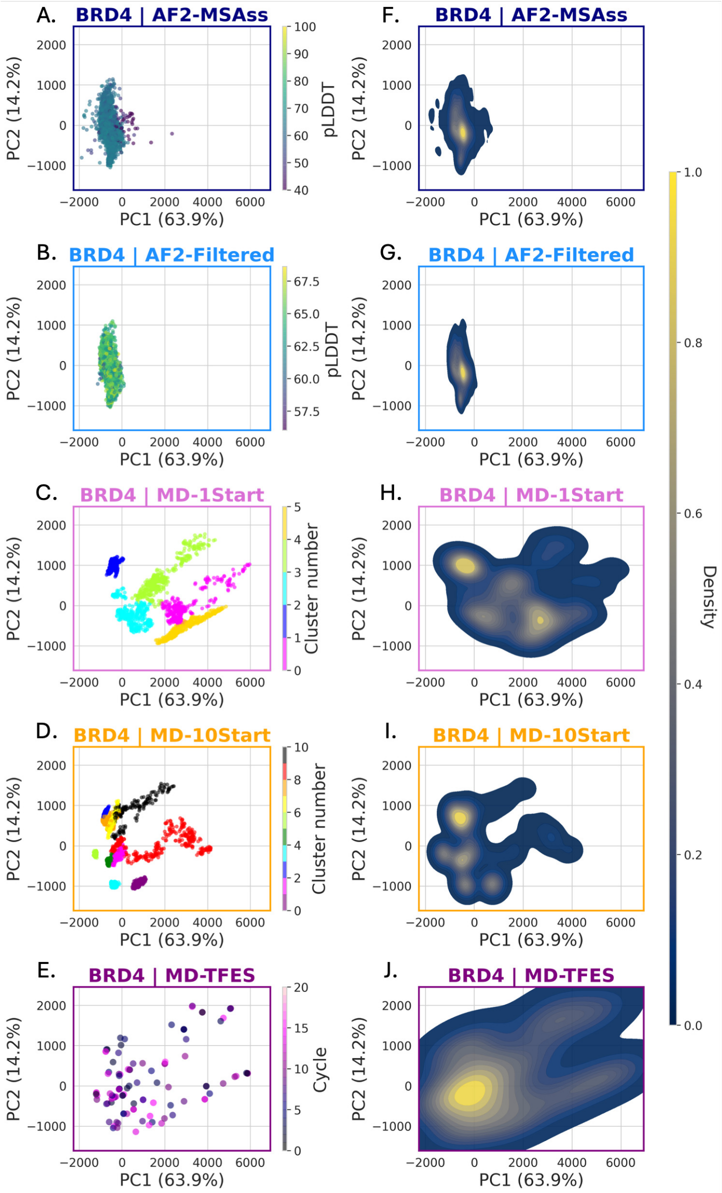


Supplementary Figure S10: BRD4 ensemble comparison through PCA reveals distinct conformational landscapes across hierarchically sampled generation protocols. Principal component analysis scatter plots (**A-E**) and corresponding density distributions (**F-J**) for five BRD4 ensembles: (**A,F**) AF2-MSAss coloured by pLDDT confidence scores, (**B,G**) AF2-Filtered, (**C,H**) MD-1Start coloured by cluster assignment, (**D,I**) MD-10Start showing multiple conformational clusters, and (**E,J**) MD-TFES displaying broad conformational sampling. Left panels show individual structure distributions in PC1-PC2 space, while right panels display density contours revealing population distributions. PCA was computed across all ensembles, MD-TFES was k-means clustered to 500 structures to better highlight the differences between cycles. For this ensemble, each point therefore represents a individual sampling distribution rather than a single structure.

##### S0.2.7 HOIP

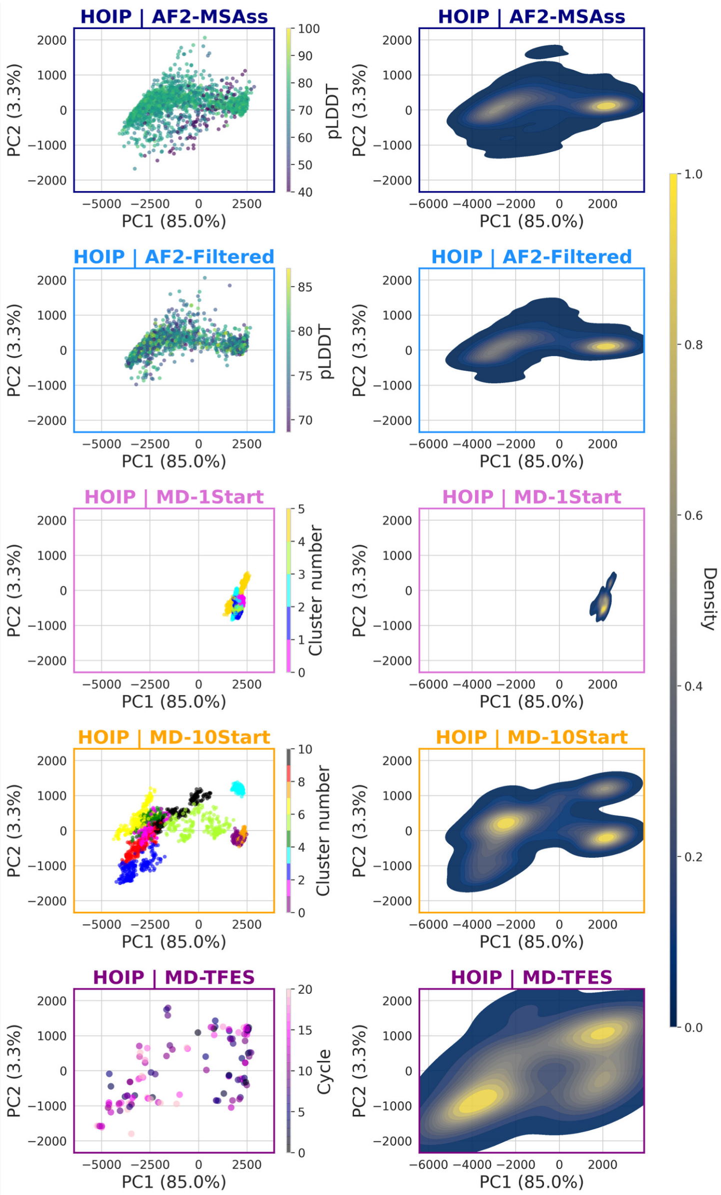


Supplementary Figure S11: HOIP ensemble comparison through PCA reveals distinct conformational landscapes across hierarchically sampled generation protocols. Principal component analysis scatter plots (**A-E**) and corresponding density distributions (**F-J**) for five HOIP ensembles: (**A,F**) AF2-MSAss coloured by pLDDT confidence scores, (**B,G**) AF2-Filtered, (**C,H**) MD-1Start coloured by cluster assignment, (**D,I**) MD-10Start showing multiple conformational clusters, and (**E,J**) MD-TFES displaying broad conformational sampling. Left panels show individual structure distributions in PC1-PC2 space, while right panels display density contours revealing population distributions. PCA was computed across all ensembles, MD-TFES was k-means clustered to 500 structures to better highlight the differences between cycles. For this ensemble, each point therefore represents a individual sampling distribution rather than a single structure.

##### S0.2.8 LXRa

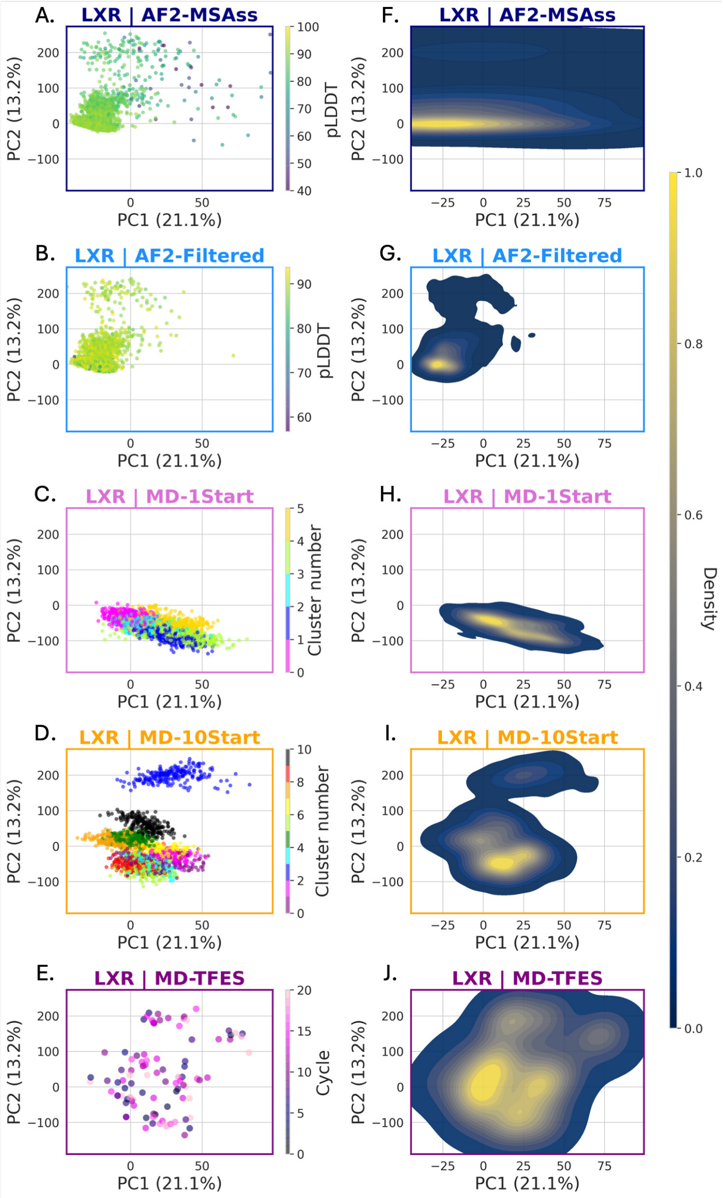


Supplementary Figure S12: LXRa ensemble comparison through PCA reveals distinct conformational landscapes across hierarchically sampled generation protocols. Principal component analysis scatter plots (**A-E**) and corresponding density distributions (**F-J**) for five LXRa ensembles: (**A,F**) AF2-MSAss coloured by pLDDT confidence scores, (**B,G**) AF2-Filtered, (**C,H**) MD-1Start coloured by cluster assignment, (**D,I**) MD-10Start showing multiple conformational clusters, and (**E,J**) MD-TFES displaying broad conformational sampling. Left panels show individual structure distributions in PC1-PC2 space, while right panels display density contours revealing population distributions. PCA was computed across all ensembles, MD-TFES was k-means clustered to 500 structures to better highlight the differences between cycles. For this ensemble, each point therefore represents a individual sampling distribution rather than a single structure.

##### S0.2.9 MBP

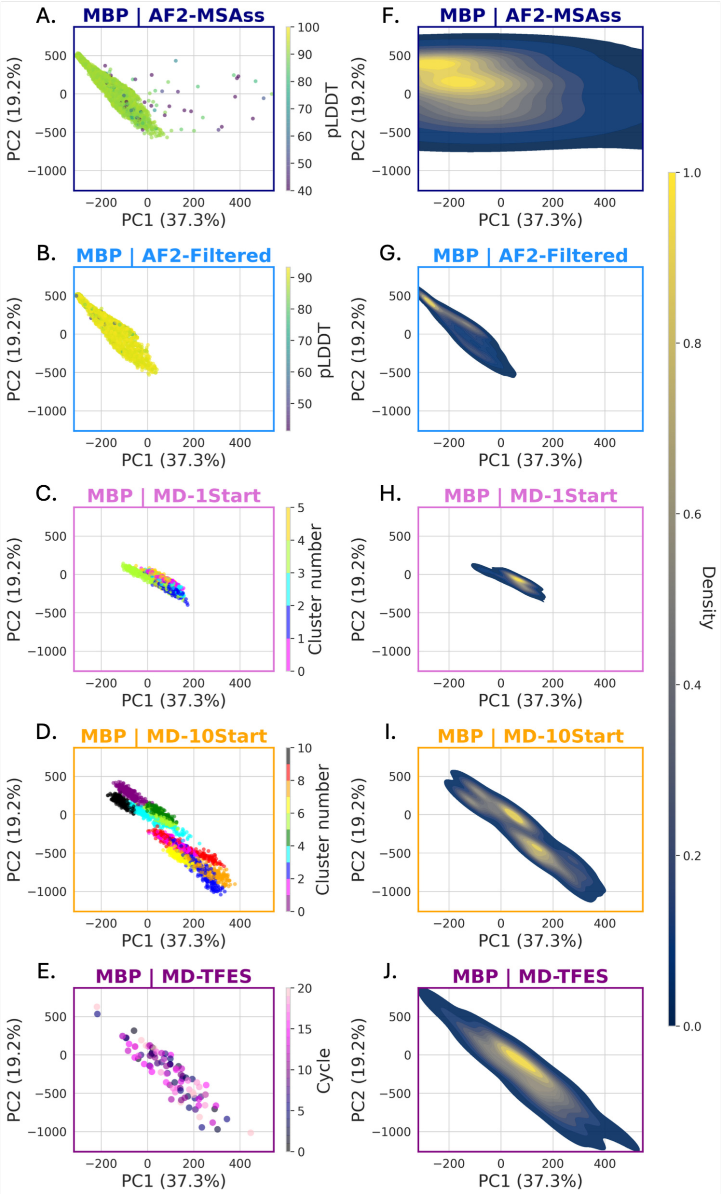


Supplementary Figure S13: MBP ensemble comparison through PCA reveals distinct conformational landscapes across hierarchically sampled generation protocols. Principal component analysis scatter plots (**A-E**) and corresponding density distributions (**F-J**) for five MBP ensembles: (**A,F**) AF2-MSAss coloured by pLDDT confidence scores, (**B,G**) AF2-Filtered, (**C,H**) MD-1Start coloured by cluster assignment, (**D,I**) MD-10Start showing multiple conformational clusters, and (**E,J**) MD-TFES displaying broad conformational sampling. Left panels show individual structure distributions in PC1-PC2 space, while right panels display density contours revealing population distributions. PCA was computed across all ensembles, MD-TFES was k-means clustered to 500 structures to better highlight the differences between cycles. For this ensemble, each point therefore represents a individual sampling distribution rather than a single structure.

#### S0.2.10 Ensemble LCC Comparison

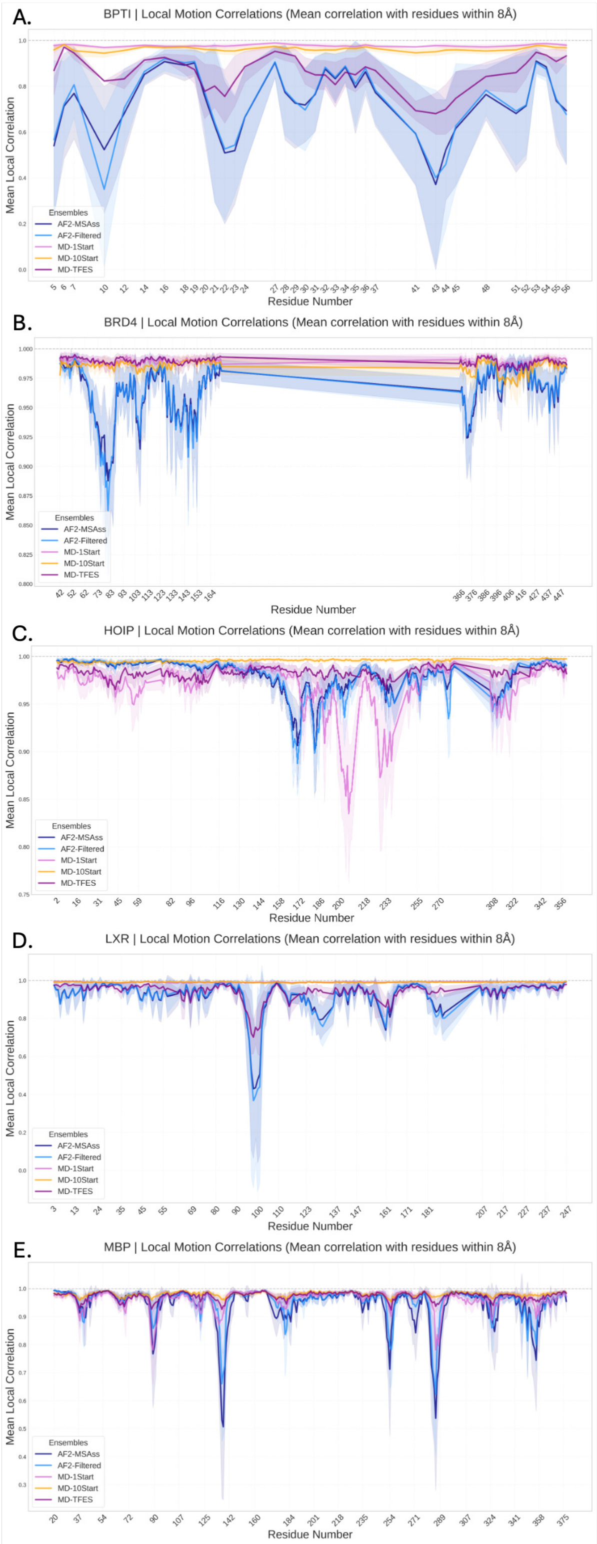


Supplementary Figure S14: Local motion correlations (LCC) reveal ensemble-specific dynamics patterns across protein systems. Mean local correlation with residues within 8Å plotted against residue number for (**A**) BPTI, (**B**) BRD4, (**C**) HOIP, (**D**) LXR, and (**E**) MBP across five ensemble types: AF2-MSAss (navy), AF2-Filtered (blue), MD-1Start (pink), MD-10Start (orange), and MD-TFES (purple), with confidence intervals shown as shaded regions. Results demonstrate that AF2-MSAss and AF2-Filtered exhibit similar local dynamics despite structure cleaning process, while MD-10Start shows the flattest correlation profiles due to short simulations across multiple starting conformations. MD-1Start displays close agreement with other ensembles in some regions but strongly diverges in other areas, as this uses a single starting structure this indicates a single conformational mode sampled. MD-TFES lies intermediate between AF2 and MD ensembles with systematic shifts reflecting uneven structural sampling from maximum diversity protocols.

##### S0.2.11 Pairwise Ensemble Comparison

##### S0.2.12 BPTI

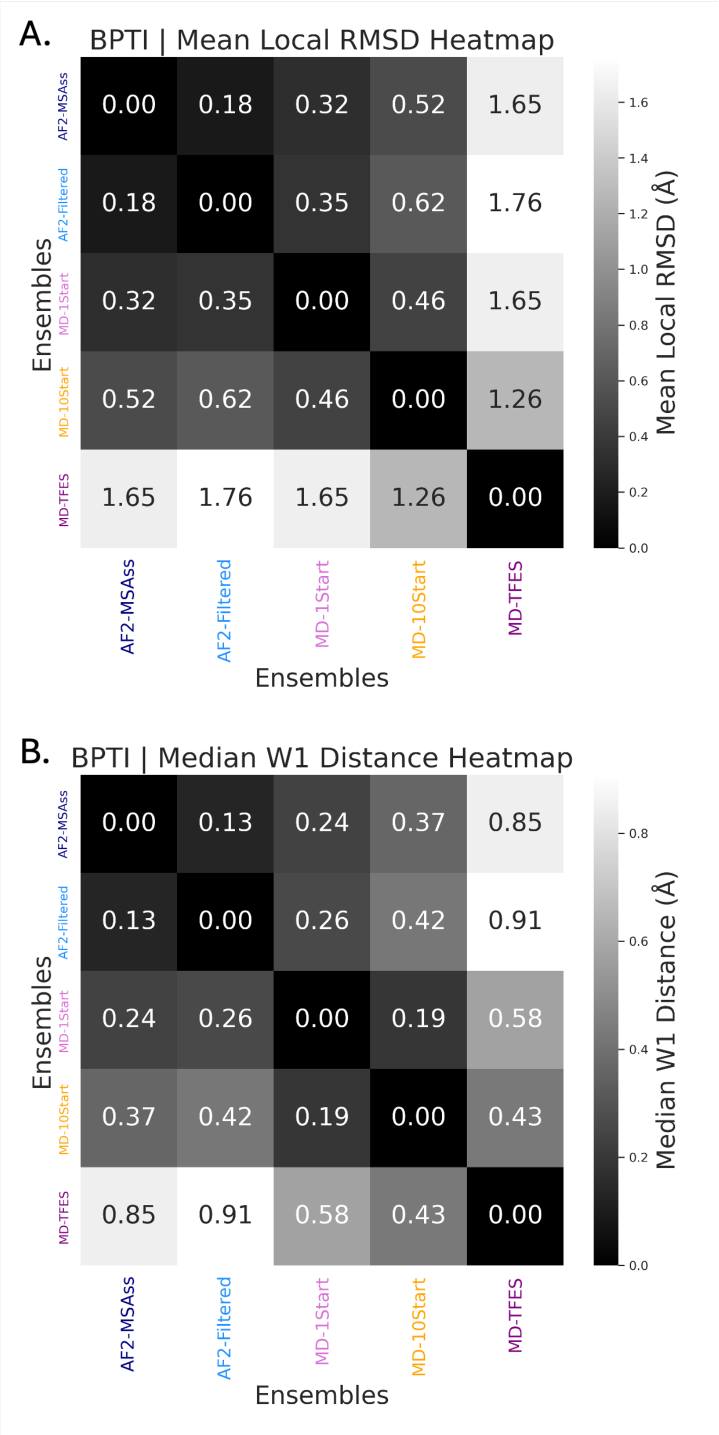


Supplementary Figure S15: BPTI pairwise ensemble structural comparison using RMSD and Wasserstein distance metrics. (**A**) Mean local RMSD heatmap showing structural differences between ensemble-averaged coordinates for five BPTI ensembles: AF2-MSAss, AF2-Filtered, MD-1Start, MD-10Start, and MD-TFES. Values represent root-mean-square deviations in Å between ensemble-averaged $C_\alpha$ positions, with darker colors indicating greater structural similarity. (**B**) Median Wasserstein-1 (W1) distance heatmap quantifying dissimilarity between ensemble distributions of residue positions. W1 distances measure the effort required to transform one ensemble's conformational distribution into another, accounting for the full conformational landscape rather than just average structures. Lower values (darker colours) indicate more similar conformational distributions. Both metrics reveal that MD-derived ensembles (MD-1Start, MD-10Start, MD-TFES) show greater structural similarity to each other compared to AF2-derived ensembles, with MD-TFES showing the largest distances from other ensembles, reflecting its enhanced sampling diversity.

##### S0.2.13 BRD4

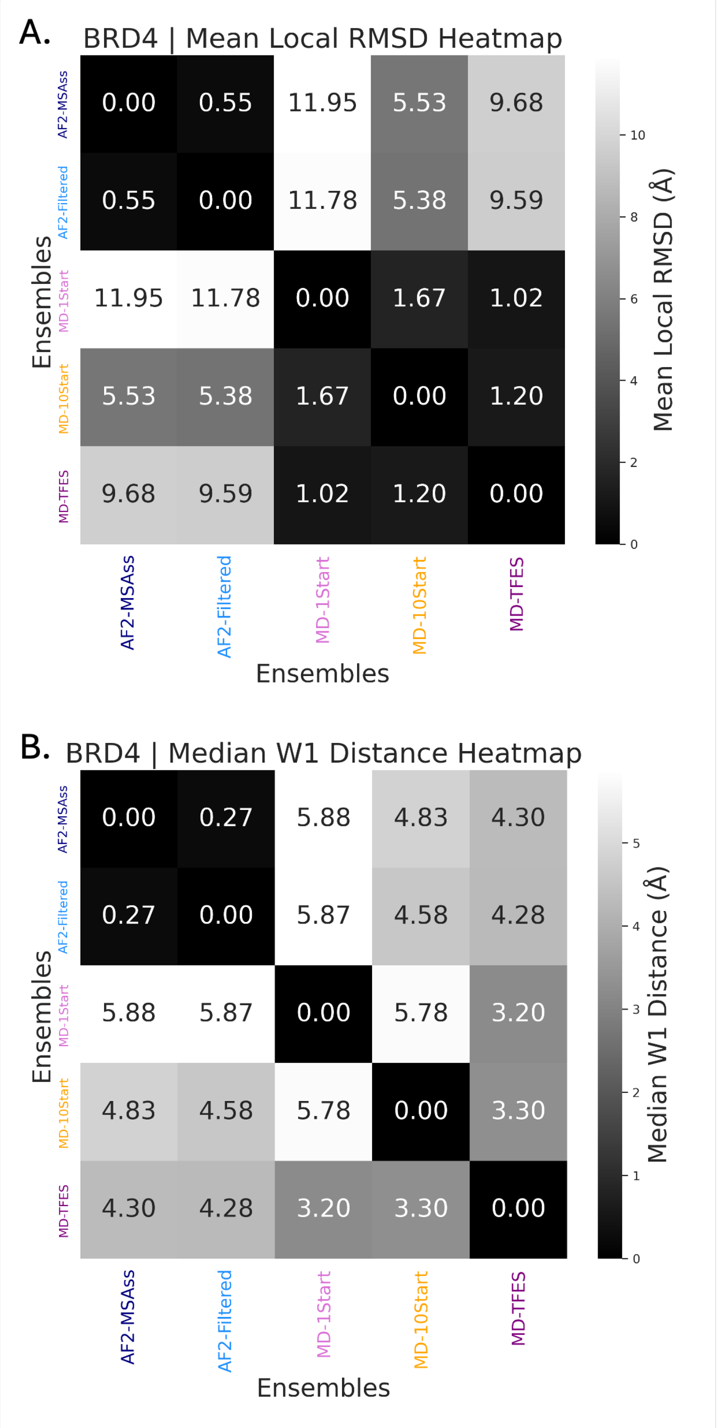


Supplementary Figure S16: BRD4 pairwise ensemble structural comparison using RMSD and Wasserstein distance metrics. (**A**) Mean local RMSD heatmap showing structural differences between ensemble-averaged coordinates for five BRD4 ensembles: AF2-MSAss, AF2-Filtered, MD-1Start, MD-10Start, and MD-TFES. Values represent root-mean-square deviations in Å between ensemble-averaged $C_\alpha$ positions, with darker colors indicating greater structural similarity. Large RMSD values (>11 Å) between AF2 and MD ensembles reflect substantial conformational differences in this flexible, partially disordered protein. (**B**) Median Wasserstein-1 (W1) distance heatmap quantifying dissimilarity between ensemble distributions of residue positions. W1 distances measure the effort required to transform one ensemble's conformational distribution into another, with higher values (3-6 Å) compared to BPTI indicating greater conformational diversity. AF2 ensembles (AF2-MSAss, AF2-Filtered) show high similarity to each other but large differences from MD-derived ensembles, while MD ensembles exhibit closer agreement among themselves. The elevated distance values reflect BRD4's inherent flexibility and the distinct conformational landscapes sampled by different generation protocols.

##### S0.2.14 HOIP

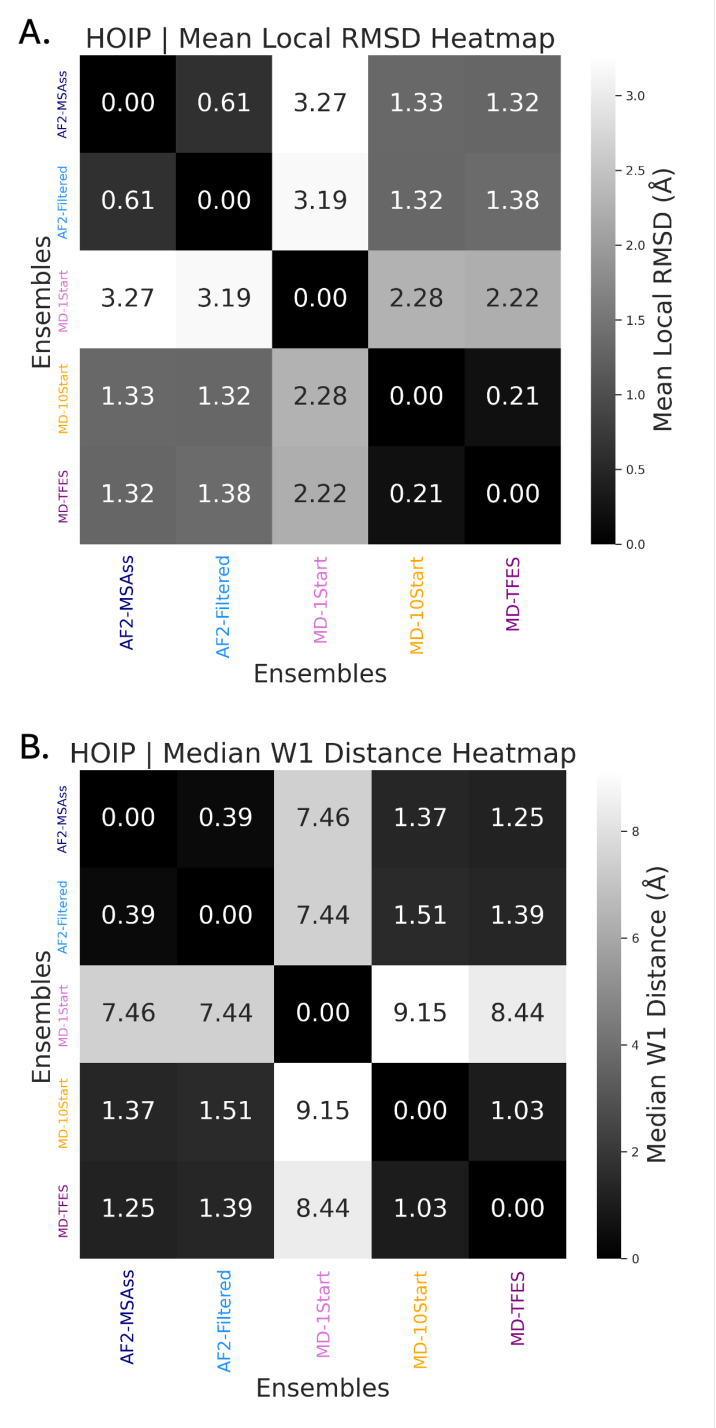


Supplementary Figure S17: HOIP pairwise ensemble structural comparison using RMSD and Wasserstein distance metrics. (**A**) Mean local RMSD heatmap showing structural differences between ensemble-averaged coordinates for five HOIP ensembles: AF2-MSAss, AF2-Filtered, MD-1Start, MD-10Start, and MD-TFES. Values represent root-mean-square deviations in Ångströms between ensemble-averaged $C_\alpha$ positions, with darker colors indicating greater structural similarity. MD-1Start shows the largest deviation ( 3.2 Å) from AF2 ensembles, while MD-10Start and MD-TFES remain closer ( 1.3 Å) to AF2 predictions. (**B**) Median Wasserstein-1 (W1) distance heatmap quantifying dissimilarity between ensemble distributions of residue positions. MD-1Start exhibits dramatically different conformational distributions compared to all other ensembles (7.4-9.2 Å), suggesting sampling of a distinct conformational region due to its single starting structure approach. In contrast, MD-10Start and MD-TFES show much closer agreement with each other (1.03 Å) and moderate similarity to AF2 ensembles ( 1.3-1.5 Å). This pattern highlights the importance of diverse starting conformations for flexible proteins like HOIP, where single-structure MD simulations may become trapped in local conformational minima.

##### S0.2.15 LXRa

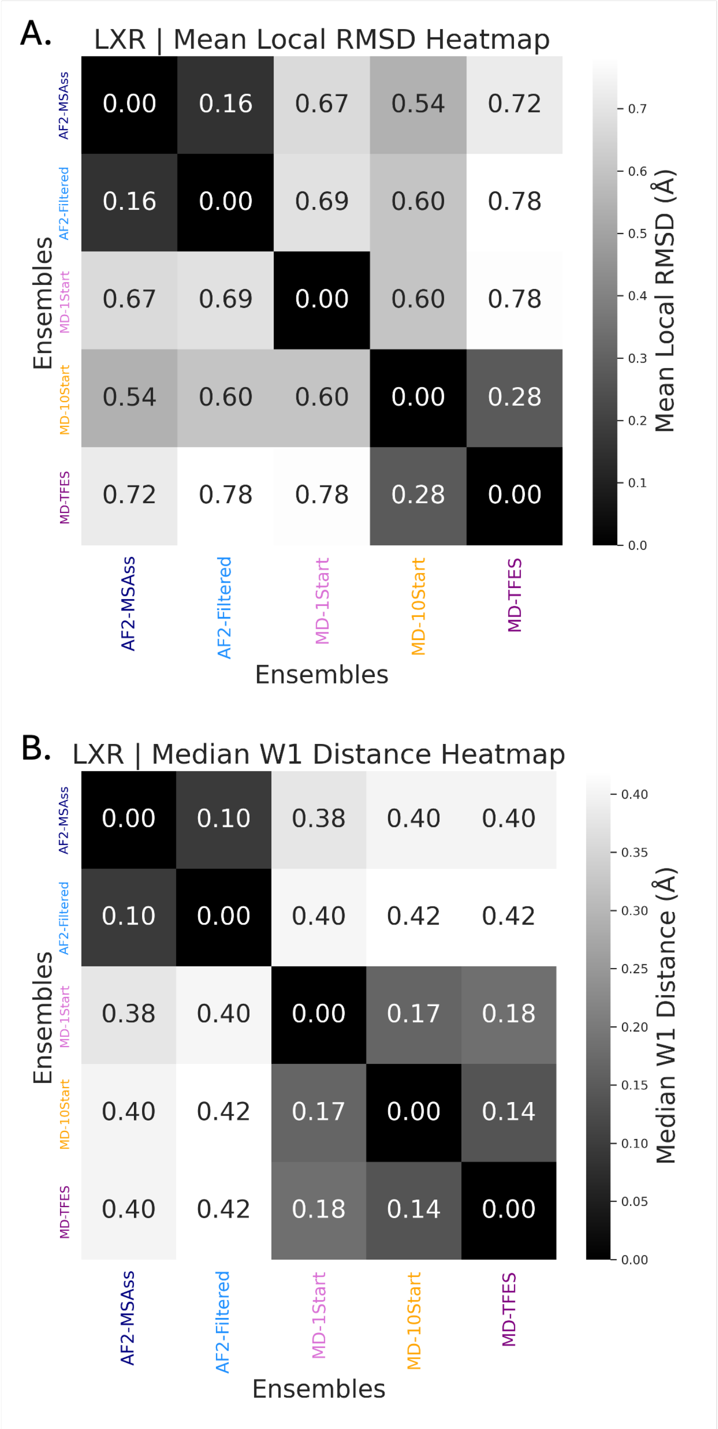


Supplementary Figure S18: LXR pairwise ensemble structural comparison using RMSD and Wasserstein distance metrics. (**A**) Mean local RMSD heatmap showing structural differences between ensemble-averaged coordinates for five LXR ensembles: AF2-MSAss, AF2-Filtered, MD-1Start, MD-10Start, and MD-TFES. Values represent root-mean-square deviations in Å between ensemble-averaged $C_\alpha$ positions, with notably small differences (0.16-0.78 Å) reflecting the rigid nature of this nuclear receptor ligand-binding domain. (**B**) Median Wasserstein-1 (W1) distance heatmap quantifying dissimilarity between ensemble distributions of residue positions. The uniformly small W1 distances (0.10-0.42 Å) indicate that all ensemble generation methods sample similar conformational distributions for this well-folded, structured protein. AF2 ensembles show high similarity (0.10 Å), while MD ensembles exhibit slightly greater diversity but remain within a narrow conformational range. The consistently low distance values across all pairwise comparisons demonstrate that LXR's rigid structure constrains sampling to a compact conformational landscape regardless of the generation protocol employed.

##### S0.2.16 MBP

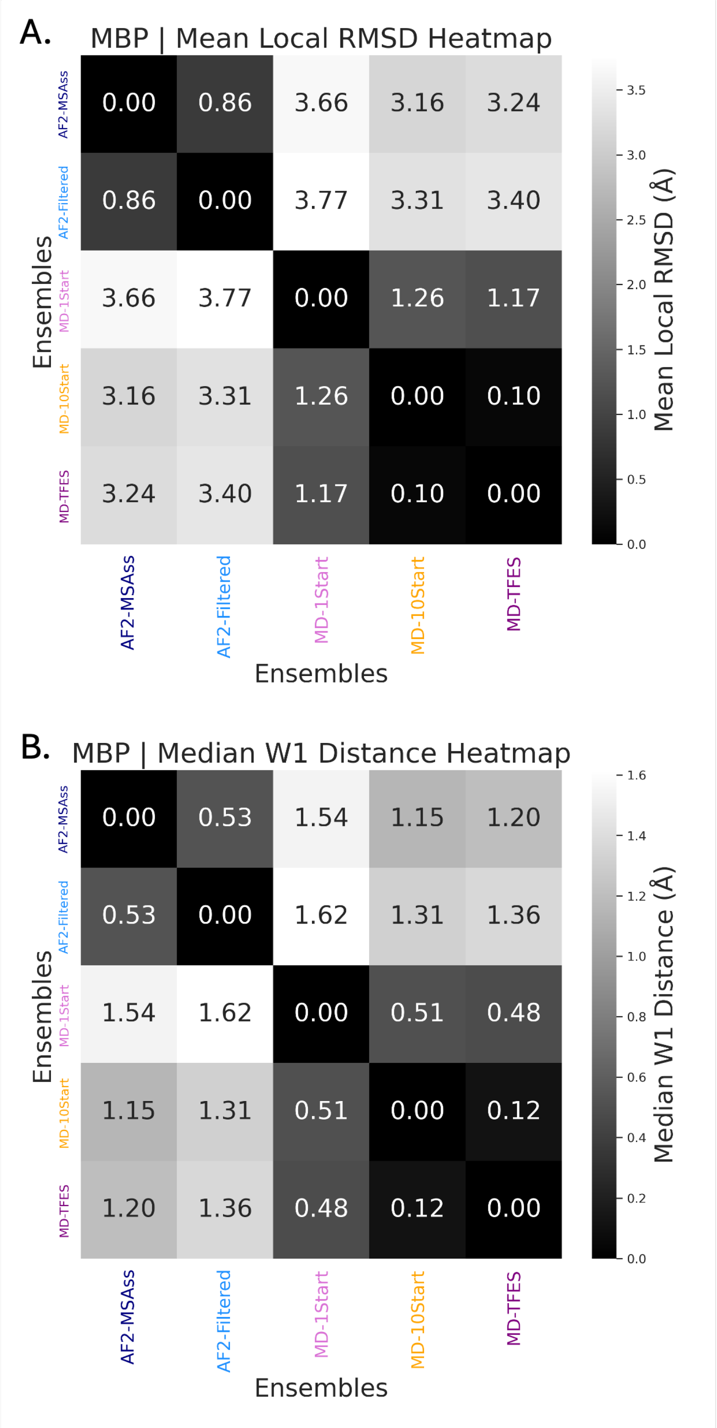


Supplementary Figure S19: MBP pairwise ensemble structural comparison using RMSD and Wasserstein distance metrics. **(A)** Mean local RMSD heatmap showing structural differences between ensemble-averaged coordinates for five MBP ensembles: AF2-MSAss, AF2-Filtered, MD-1Start, MD-10Start, and MD-TFES. Values represent root-mean-square deviations in Å between ensemble-averaged $C_\alpha$ positions. AF2 and MD ensembles show substantial structural differences ( 3.2-3.8 Å), while MD-derived ensembles exhibit closer agreement among themselves, particularly MD-10Start and MD-TFES (0.10 Å). **(B)** Median Wasserstein-1 (W1) distance heatmap quantifying dissimilarity between ensemble distributions of residue positions. The W1 distances reveal more moderate conformational differences (0.12-1.6 Å) compared to the highly flexible BRD4, reflecting MBP's intermediate flexibility. AF2 ensembles show good agreement (0.53 Å), while MD ensembles demonstrate close conformational sampling convergence, especially between MD-10Start and MD-TFES (0.12 Å). The pattern suggests that multiple-start MD protocols (MD-10Start) and enhanced sampling (MD-TFES) converge on similar conformational landscapes for this maltose-binding protein, distinct from AF2 predictions.

#### S0.3 Derivation of Work Done Metrics

##### S0.3.1 $\Delta H$ (Enthalpy change)

We first start with the Linderstrom-Lang definition for the Free Energy change due to the 'opening', of a backbone amide proton position obtained through the relative rate between the extrinsic (measured) and intrinsic (innate chemical potential).

$$k_{\text{extrinsic}}(i) = K_{\text{opening}}(i) k_{\text{intrinsic}}(i)$$
$$\Delta G(i) = -RT \ln \frac{k_{\text{extrinsic}}(i)}{k_{\text{intrinsic}}(i)}$$
$$\Delta H(i) \equiv \Delta G(i) = -RT \log P_f(i)$$
(Linderstrom-Lang Free Proton-Exchange Free Energy)

**Where:**

- $\Delta H(i)$ is the enthalpy change for residue $i$
- $\Delta G(i)$ is the Gibbs free energy change
- $k_{\text{extrinsic}}(i)$ is the measured rate of opening
- $k_{\text{intrinsic}}(i)$ is the intrinsic chemical exchange rate
- $K_{\text{opening}}(i)$ is the equilibrium constant of unfolding
- $P_f(i)$ is the protection factor

For the case of a single residue we propose that this is equivalent to the Enthalpy through the free energy relation, Equation Free Energy Change, where $\Delta S = 0$ in the case of a single residue in isolation.

$$\Delta G = \Delta H - T\Delta S \qquad \text{(Free Energy Change)}$$

**Where:**

- $\Delta G$ is the Gibbs free energy change of opening
- $\Delta H$ is the enthalpy change of opening
- $\Delta S$ is the entropy change of opening

To demonstrate that Enthalpy and Gibbs Free Energy is appropriate in this case we substitue the previous definitions of internal energy into the Enthalpy - Internal Enthalpy relation to generate Equation HDX Enthalpy - Internal Energy Relation. In the case of protection factors described by empirical models, we propose $\Delta V = 0$. In general, when predicting protection-factors, any volume changes are directly considered since the potential must be expressed as a locally-weighted density of interactions. Since the BV model uses contacts as inputs, any ensemble featurised in this way has a fixed volume. Many other featurisers also operate at a fixed volume, featurisers that use a sigmoidal ″switch″ function that tends to infinity, the volume measured is still constant. This extends for other empirical models that also use cut-offs for computational efficiency.

$$\Delta H(X) = \Delta U(X) + P\Delta V(X)$$

$$\Delta V(X) = 0 \qquad \text{(HDX Enthalpy - Internal Energy Relation)}$$

therefore :

$$\Delta H(X) \equiv \Delta U(X)$$

**Where:**

- $\Delta H(X)$ is the HDX enthalpy change of opening
- $\Delta U(X)$ is the internal energy change of opening
- $P$ is pressure
- $\Delta V(X)$ is the volume change of opening (zero in this case)

In these cases, Internal-Energy $U(X)$, and Enthalpy $H(X)$, are equivalent. The distinction is therefore to note the difference to the metrics used by HDXer. $H_{\text{optimisation}}(X)$ or $H_{\text{opt}}(X)$ therefore measures the total change in Enthalpy due to Optimisation and is calculated in Equation Enthalpy Change of Optimisation Definition. Where, $\hat{X}$, is an optimised ensemble.

$$\Delta\hat{H}(X) = \hat{H}(X) - H(X) \qquad \text{(Enthalpy Change of Optimisation Definition)}$$

**Where:**

- $\hat{H}(X)$ is the optimized enthalpy
- $H(X)$ is the initial enthalpy

Since an ensemble, $X$, requires additional parameters to be used in optimisation, an ideal quantity to measure change of optimisation should reflect this. Equation Enthalpy Definition shows this for the BV model case which differs from the standard approach as the $\beta$ energy scaling parameters are described as ensemble specific, $\beta(X)$, instead of global parameters.

$$H(X) = RT\,\langle \beta_{\mathrm{C}}(X)N_{\mathrm{C}}(X) + \beta_{\mathrm{H}}(X)N_{\mathrm{H}}(X)\rangle \qquad \text{(Enthalpy Definition)}$$

**Where:**

- $H(X)$ is the initial enthalpy of ensemble $X$
- $\hat{H}(X)$ is the enthalpy of optimised ensemble $\hat{X}$
- $\beta_{\mathrm{C}}(X), \beta_{\mathrm{H}}(X)$ are ensemble-specific energy scaling parameters
- $N_{\mathrm{C}}(X), N_{\mathrm{H}}(X)$ are ensemble-specific heavy-atom and hydrogen-bond contacts

##### S0.3.2 $\Delta S$ (Entropy change)

In order to complete the Free Energy relation, Equation Free Energy Change, we require the Entropy term. To derive this expression we start with the state definitions themselves. In a Gibbs entropy framework the probability of a state to the reference state can be calculated from the Boltzmann equation Equation Boltzmann probability and Ensemble Entropy. From this we can construct an entropy equation for the ensemble average from the individual components of each residue. The probability of a given state $p(i)$ is the exponent of $\Delta H(i)$ from the Generalised-Boltzmann formulation as we are operating on ensemble averages.

$$S(\langle X\rangle) = -R\sum_{i\in x} p(i)\log p(i)$$

$$p(i) = \frac{1}{Z}e^{\frac{-\Delta H(i)}{RT}} \qquad \text{(Boltzmann probability and Ensemble Entropy)}$$

$$Z = \frac{1}{\sum_{i\in x} e^{\frac{-\Delta H(i)}{RT}}}$$

**Where:**

- $S(\langle X\rangle)$ is the ensemble entropy
- $p(\langle i\rangle)$ is the probability of state $i$
- $Z$ is the partition function
- $\Delta H_{\mathrm{opt}}$ is the Potential of Mean Force Enthalpy change of optimsiation

Since $X$ represents an ensemble, it can also be described as a weighted, $p(x)$ average of substructures, $x$. For simplicity, these weights can be normalised to equal 1, as shown in Equation Ensemble Averaged Enthalpy.

$$\langle H(X)\rangle = \sum_{x\in X} p_{\text{frames}}(x)H(x)$$
$$\sum_{x\in X} p_{\text{frames}}(x) = 1 \quad \text{(Ensemble Averaged Enthalpy)}$$

**Where:**

- $\langle H(X)\rangle$ is the ensemble-averaged enthalpy for the entire ensemble $X$
- $p_{\text{frames}}(x)$ is the population fraction (weight) of structure $x$
- $H(x)$ is the enthalpy of an individual structure $x$
- $\sum_{x\in X}$ represents the sum over all structures in ensemble $X$

The probabilities, $p(x)$ in Equation Ensemble Averaged Enthalpy represent the population fraction for a given discrete microstate, $x$, with respect to the whole ensemble, $X$. From this quantised perspective, Entropy of an ensemble, $S(X)$, can be expressed in the Gibbs Entropy expression in Equation Ensemble Averaged Entropy of Reweighting.

$$S_{\text{RW-only}}(\langle X\rangle) \approx S_{\text{frames}}(\langle X\rangle) = -R\sum_{x\in X} p_{\text{frames}}(x)\log p_{\text{frames}}(x) \quad \text{(Ensemble Averaged Entropy of Reweighting)}$$

**Where:**

- $S_{\text{frames}}(\langle X\rangle)$ is the thermodynamically scaled entropy of the ensemble $X$ due changing population weights during fitting
- $p_{\text{frames}}(x)$ is the population fraction of structure $x$

Note: this considers the case where protection factors are only a product of their frame weights, meaning it only measures the Work Done due to reweighting and does not consider contribution from other sources. These include: model parameters and structural similarity.

$$\hat{S}(\langle X\rangle) - S(\langle X\rangle) = \Delta\hat{S}(X) \neq \Delta\hat{S}_{\text{frames}}(X) \quad \text{(Entropy Change Definition)}$$

**Where:**

- $\hat{S}(\langle X \rangle)$ is the optimized entropy
- $S(\langle X \rangle)$ is the initial entropy
- $\Delta S_{\text{frames}}(X)$ is the entropy change due to reweighting

For an appropriate expression for Entropy change at the ensemble-average level, we wish to be able to quantify the total contribution of changes during optimisation in a way that is compatible with the requirement of consistent state definitions. We chose to use an absolute mean-centring for $\Delta H_{\text{opt}}$ as in the context of model optimisation of HDX-MS data, the overall scale of exchange is more reflective of the experimental conditions. Since the scale change is also a useful metric, we can also define $\Delta \hat{H}_{\text{abs}}(\langle i \rangle)$ which reflects the change in magnitude or scale in protection factors for both reweighting-only and BV-only optimisation. As enthalpy is additive, it is clear how $\Delta \hat{H}_{\text{opt}}$ is a component of the absolute norm $|\Delta|\hat{H}$ instead of $\Delta \hat{H}$. In the context of optimisation, this is intuitive as a metric as any change should have a cost regardless of the direction. Moreover, the quantity of interest is the change in distribution of the ensemble-average conformational dynamics. As a result, $\Delta \hat{S}_{\text{opt}}$ derived from $H_{\text{opt}}$ is constrained to measuring the conformational change applied during optimisation on the ensemble-average protection factors.

$$
\begin{aligned}
H_{\text{opt}}(i) &= |H(i) - \langle H_{\text{i}}(\langle X \rangle) \rangle| \\
\text{Work}_{\text{shape}} = \Delta \hat{H}_{\text{opt}}(i) &= |H_{\text{opt}}(i) - \hat{H}_{\text{opt}}(i)| \\
\text{Work}_{\text{scale}} = \Delta \hat{H}_{\text{abs}}(i) &= |\langle \hat{H}_{\text{i}}(\langle X \rangle) \rangle - \langle H_{\text{i}}(\langle X \rangle) \rangle| \\
|\Delta|\hat{H}(i) &= \Delta \hat{H}_{\text{opt}}(i) + \Delta \hat{H}_{\text{abs}}(i) \\
p_{\text{opt}}(i) &= e^{-\frac{H_{\text{opt}}(i)}{RT}} = (|P_{\text{f}}(i) - \langle P_{\text{f}}(\langle X \rangle) \rangle|)^{-1} \\
S_{\text{opt}}(i) &= -R \sum_{i \in X} p_{\text{opt}}(i) \log(p_{\text{opt}}(i)) \\
\text{Work}_{\text{density}} = \Delta S_{\text{opt}}(i) &= |S_{\text{opt}}(i) - \hat{S}_{\text{opt}}(i)| \\
\text{Work}_{\text{opt}} = \Delta G_{\text{opt}}(i) &= \Delta H_{\text{opt}}(i) + \Delta S_{\text{opt}}(i)
\end{aligned}
$$
(Calculation of Probabilities from Protection Factors)

**Where:**

- $\sum_{i \in X} p_{\text{opt}}(i) = 1$
- $p_{\text{opt}}(i)$ is the pseudo-probability of residue $i$ in the ensemble average, considering frame-weights and model parameter, enthalpic ($\Delta H_{\text{opt}}$) contributions for each residue.
- $P_{\text{f}}(i)$ is the change in optimized protection factor for residue $i$

We do not define a $|\Delta S|$ or $\Delta S_{\text{abs}}$ as there is no clear interpretation for these. In practice, even in reweighting-only optimisation the entropic contributions due to the change in scale can be far in excess of the change in shape.

$$
\begin{aligned}
\text{Apparent Work}_{\text{HDXer}} &= \Delta U_{\text{opt}}(X), \\
\text{Apparent Work} &\neq \text{Work}, \\
\Delta H_{*}(X) &= |\Delta H_{\text{opt, shape}}(X)| + |\Delta H_{\text{opt, scale}}(X)|, \\
\Delta G_{*}(X) &= \Delta H_{*}(X) - T\Delta S_{*}(X), \\
\Delta H_{\text{opt, shape}}(X_1, X_2) &= \Delta H_{\text{opt}}(X) = k_{\text{B}}T \sum_i \left|\log P_f(\langle i_1\rangle) - \log P_f(\langle i_2\rangle)\right|, \\
\Delta H_{\text{opt, scale}}(X_1, X_2) &= \Delta H_{\text{abs}}(X) = k_{\text{B}}T \left|\overline{\log P_f}(\langle i_1\rangle) - \overline{\log P_f}(\langle i_2\rangle)\right|, \\
\Delta G_{\text{opt, shape}}(X_1, X_2) &= \Delta G_{\text{opt}}(X) = \Delta H_{\text{opt}}(X) - T\Delta S_{\text{opt}}(X), \\
\Delta H_{\text{opt}}(X) &= \text{Work}_{\text{shape}}, \\
\Delta H_{\text{abs}}(X) &= \text{Work}_{\text{scale}}, \\
-T\Delta S_{\text{opt}}(X) &= \text{Work}_{\text{density}}, \\
\Delta G_{\text{opt}}(X) &= \text{Work}_{\text{opt}},
\end{aligned}
\tag{Definitions of Work Done during optimisation}
$$

#### S0.4 Supplementary Methods

##### S0.4.1 AlphaFold Ensemble Generation Details

| Protein | BPTI | BRD4 | HOIP | LXRa | MBP |
|---|---|---|---|---|---|
| **N. Residues** | 58 | 480 | 376 | 247 | 396 |
| **Flexibility** | Rigid | Partially Disordered | Flexible | Rigid | Flexible |
| **PDB Available** | Yes | Binding constructs only | Complex only | Yes | Yes |
| **PDB Code** | 5PTI | BD1:8GPZ, BD2:6C7Q (HOLO) | 6CS6 (Complex) | 2ACL (HOLO) | 1ANF (HOLO) |
| **Min. Confidence** | 2.5% | 5% | 10% | 1.25% | 5% |
| **Contact Order** | 1.5%, 99.5% | 2%, 99% | 3%, 99.5% | 1.5%, 99.5% | 3%, 99.5% |
| **Correlation** | 1.5%, 99.5% | 2%, 99% | 3%, 99.5% | 1.5%, 99.5% | 3%, 99.5% |
| **Res. Distance** | 0.01%, 99.5% | 0.01%, 99.5% | 0.01%, 99.5% | 0.01%, 99.5% | 0.01%, 99.5% |

Supplementary Table S4: Parameters for AF2 Ensemble cleaning

##### S0.4.2 Structure Poisoning Algorithms

**Algorithm 1** Gaussian Noise Structure Poisoning

**Require:** Ensemble $X = \{x_1, x_2, \dots, x_n\}$

**Require:** Output path for poisoned structures

**Ensure:** Poisoned ensemble with residue-specific Gaussian noise

```
Initialize σ_residue ← {}                                   ▷ Residue variance dictionary
for each structure x ∈ X do
    for each residue r in x do
        Store coordinates: σ_residue[r.id].append(r.positions)
    end for
end for
for each residue ID r_id in σ_residue do
    positions ← array(σ_residue[r_id])
    σ_residue[r_id] ← var(positions, axis = 0)
end for
for each structure x ∈ X do
    for each residue r in x do
        noise ← N(0, sqrt(σ_residue[r.id]))
        r.positions ← r.positions + noise
    end for
    Write poisoned structure to output
end for
```

**Gaussian Noise.**

**Algorithm 2** Coordinate Mixing Structure Poisoning

**Require:** Ensemble $X = \{x_1, x_2, \ldots, x_n\}$

**Require:** Output path for poisoned structures

**Ensure:** Poisoned ensemble with randomly permuted coordinates

**for** each structure $x \in X$ **do**
    coordinates ← $x$.atoms.positions ▷ Extract all atomic coordinates
    shuffled_coords ← random_permutation(coordinates)
    $x$.atoms.positions ← shuffled_coords
    Write poisoned structure to output
**end for**

**Coordinate Mixing.**

**Algorithm 3** Backbone Proton Shuffling Structure Poisoning

**Require:** Ensemble $X = \{x_1, x_2, \ldots, x_n\}$

**Require:** Output path for poisoned structures

**Ensure:** Poisoned ensemble with shuffled backbone proton positions

**for** each structure $x \in X$ **do**
    backbone_H ← select_atoms("backbone and name H")
    original_coords ← backbone_H.positions.copy()
    shuffled_coords ← random_shuffle(original_coords)
    backbone_H.positions ← shuffled_coords
    Write poisoned structure to output
**end for**

**Proton Shuffling.**

#### S0.5 Weighted Structural Metrics

##### S0.5.1 Weighted RMSD

To avoid the alignment problem and account for ensemble weights, we computed the pairwise ensemble RMSD using internal coordinates:

$$
\begin{aligned}
\langle \mathrm{RMSD}_i(X_1, X_2)\rangle &= \frac{1}{N_i^2}\sqrt{(\langle C_\alpha(X_1)\rangle - \langle C_\alpha(X_2)\rangle)^2} \\
\text{where:} \quad \langle C_\alpha(X)\rangle &= \sum_{x\in X} p(x) C_\alpha(x) \qquad \text{(Pairwise Ensemble RMSD)} \\
\text{and:} \quad C_\alpha(x) &= \text{Pairwise } C_\alpha \text{ coordinates} \\
N_i &= \text{Number of residues, } i\text{, in structure, } x
\end{aligned}
$$

##### S0.5.2 Wasserstein Distance Details

The full calculation of the Wasserstein-1 distance, including Kernel Density Estimation (KDE) and weighting terms, is defined as:

$$
\begin{aligned}
\langle W_{1,i}(X_1, X_2)\rangle &= \frac{1}{N_i}\sum_{i=1}^{N_i} W_{\text{KDE}}(\Omega_1, \Omega_2) \\
\text{where:} \quad \Omega &= \text{KDE}(C_\alpha(X)) \\
W_{\text{KDE}}(\Omega_1, \Omega_2) &= \inf_{\gamma_{\text{HDXer}}\in\Gamma(\Omega_1,\Omega_2)} \sum_{x_1\in\Omega_1}\sum_{x_2\in\Omega_2} \gamma(x_1, x_2)\alpha(x_1, x_2)|\omega C_\alpha(x_1) - \omega C_\alpha(x_2)| \\
\text{where:} \quad \alpha(x_1, x_2) &= \frac{2p(x_1)p(x_2)}{p(x_1) + p(x_2)} \\
\gamma(C_\alpha(x_1), C_\alpha(x_2)) &\equiv \gamma(\omega_{C_\alpha}(x_1), \omega_{C_\alpha}(x_2)) = 1 \\
\text{and:} \quad \omega_{C_\alpha}(x) &= \text{KDE of pairwise } C_\alpha \text{ coordinates at substructure } x
\end{aligned}
$$

(Residue-average Wasserstein-1 Distance with KDE for Discrete Frames)

As $W_{1,i}$ distance requires normalised distributions, we use a kernel density estimate (KDE). Scipy's KDE implementation uses Scott's rule for bandwidth selection.[1]

#### S0.6 Computational Environment and Performance

##### S0.6.1 MD Simulations

**Traditional MD Simulations** MD simulations were performed using the University of Oxford's Advanced Research Cluster (ARC).[2] The internally built GROMACS (2021.3) module was used for all simulations.

| Protein | Performance (ns/day) | GPU | CPU | Box Length (nm) | Particles |
|---|---|---|---|---|---|
| BPTI | 282 | Tesla V100 | 4x Intel Xeon E5-2698 | 6.02 | 21,709 |
| BRD4 | 26.3 | Tesla V100/Quadro RTX 8000 | 4x Intel Xeon E5-2698/4x Intel Xeon Platinum 8268 | 18.6 | 636,632 |
| HOIP | 62.3 | Tesla V100 | 4x Intel Xeon E5-2698 | 13.3 | 233,244 |
| LXRa | 412 | A100-40GB | 4x Intel Xeon Platinum 8268 | 9.05 | 73,377 |
| MBP | 207 | Quadro RTX 8000 | 4x Intel Xeon Platinum 8268 | 10.4 | 212,741 |

Supplementary Table S5: MD Simulation Performance and System Details

**Enhanced MD Simulations** Enhanced MD simulations, including T-FES (multi-Topology Frontier Expansion Sampling), were performed using the University of Oxford's Advanced Research Cluster (ARC).[2] All enhanced sampling protocols were implemented using OpenMM 8.1.2 within a conda environment (anaconda 2022.10). The computational environment included Python 3.11 with the following key packages: NumPy 1.26.4 for numerical operations, SciPy 1.14.1 for scientific computing, scikit-learn 1.5.2 for machine learning algorithms, MDTraj 1.10.0 for trajectory analysis, matplotlib 3.9.2 for visualisation, and ParmEd 4.2.2 for parameter file conversion between GROMACS and AMBER formats.

| Protein | GPU | Performance (ns/day) |
|---|---|---|
| BPTI | A100-40GB | 557 |
| BRD4 | A100-40GB | 46.9 |
| HOIP | A100-40GB | 154 |
| LXRa | Tesla V100 | 156 |
| MBP | Tesla V100 | 54.2 |

Supplementary Table S6: Enhanced MD Simulation Performance

##### S0.6.2 AlphaFold2 Simulations

AlphaFold2 simulations were performed via localcolabfold,[3, 4] using ARC.[2] The package versions for this environment are as follows: localcolabfold (github commit=1208ceb) is the main package for simulations using python (3.10), cudatoolkit (11.8) for GPU acceleration, biopython (1.82) for handling biomolecular file formats, kalign2 (2.04); hhsuite (3.3.0) and; mmseqs2 (17.b804f) for sequence alignment, jax[cuda11] (0.4.23) numerical framework, matplotlib (3.10.0) for plotting, tensorflow (2.18.0) for numerical operations.

| Protein | GPU | Performance (s/structure) |
|---|---|---|
| BPTI | Quadro RTX 8000 | 1.5 |
| BRD4 | Quadro RTX 8000 | 25 |
| HOIP | Quadro RTX 8000 | 15 |
| LXRa | Quadro RTX 8000 | 9 |
| MBP | Quadro RTX 8000 | 20 |

Supplementary Table S7: AF2 Structure Prediction Performance

###### S0.6.3 Weighted RMSF

To explicitly account for the non-uniform weights of structures in the ensemble, the weighted RMSF for residue $i$ is computed as:

$$\begin{aligned} \text{RMSF}_i(X) &= \sqrt{\sum_{x\in X} p(x)\|\mathbf{r}_i(x) - \langle \mathbf{r}_i(X)\rangle\|^2} \\ \text{where:}\quad \langle \mathbf{r}_i(X)\rangle &= \sum_{x\in X} p(x)\mathbf{r}_i(x) \end{aligned} \tag{Weighted Residue-level RMSF}$$

###### S0.6.4 Inter-Ensemble LCC

To compare dynamical correlations across different ensembles, we computed the Inter-Ensemble LCC:

$$\begin{aligned} \text{structure-LCC}(X_1, X_2) &= \frac{2\langle \mathbf{d}_i(X_1)\cdot \mathbf{d}_i(X_2)\rangle}{\langle |\mathbf{d}_i(X_1)|^2\rangle + \langle |\mathbf{d}_i(X_2)|^2\rangle} \\ \text{where:}\quad \langle \mathbf{d}_i(X_1)\cdot \mathbf{d}_i(X_2)\rangle &= \sum_{x_1\in X_1}\sum_{x_2\in X_2} \alpha(x_1, x_2)[\mathbf{d}_i(x_1)\cdot \mathbf{d}_i(x_2)] \\ \langle |\mathbf{d}_i(X_k)|^2\rangle &= \sum_{x\in X_k} p(x)|\mathbf{d}_i(x)|^2 \quad \text{for } k\in\{1,2\} \\ \text{and:}\quad \mathbf{d}_i(x) &= \{C_{\alpha,j}(x) - C_{\alpha,i}(x) : |C_{\alpha,j}(x) - C_{\alpha,i}(x)| \le 8\text{Å}\} \\ \text{where:}\quad C_{\alpha,i}(x) &= \text{pairwise C}\alpha\text{ coordinates of residue } i \text{ in conformation } x \\ \text{and:}\quad \alpha(x_1, x_2) &= \frac{2p(x_1)p(x_2)}{p(x_1)+p(x_2)} \end{aligned} \tag{Inter-Ensemble Residue-level LCC (8 Å cutoff)}$$

#### S0.7 Knee Optimizer Algorithm

**Algorithm 4** Curve-of-the-Knee $\gamma_{\text{HDXer}}$ Optimisation

**Require:** Set of candidate values $\gamma_{\text{HDXer}} = \{\gamma_1, \gamma_2, \ldots, \gamma_n\}$
**Require:** Corresponding error values $E = \{e_1, e_2, \ldots, e_n\}$
**Require:** Apparent work values $W = \{w_1, w_2, \ldots, w_n\}$
**Ensure:** Optimal $\gamma^*$

Initialize $x \leftarrow W$, $y \leftarrow E$
Sort data points by $x$ values
**if** $|x| = 1$ **then return** $\gamma_1$
**else if** $|x| = 0$ **then**
  **raise** ValueError("No valid gamma values found")
**end if**
$\gamma^* \leftarrow$ null
// *Method 1: Angle-based selection*
$\theta \leftarrow []$
**for** $i = 1$ to $|x| - 1$ **do**
  $\theta_i \leftarrow \arctan\left(\frac{y_{i+1}-y_i}{x_{i+1}-x_i}\right)$
  Append $\theta_i$ to $\theta$
**end for**
**if** $\theta \neq []$ **then**
  $\theta^* \leftarrow \arg\min_{\theta_i} |\theta_i - \pi/4|$
  $\gamma^* \leftarrow \gamma_{\text{index}(\theta^*)}$
**end if**
**if** $\gamma^* =$ null **then**
  // *Method 2: Maximum distance from regression line*
  Fit linear regression: $\hat{y} = mx + b$ where $(m, b) =$ polyfit$(x, y, 1)$
  $d \leftarrow []$
  **for** $i = 1$ to $|x|$ **do**
    $d_i \leftarrow |y_i - (mx_i + b)|$
    **if** $y_i < mx_i + b$ **then**
      $d_i \leftarrow -d_i$ // *Negative for points below regression line*
    **end if**
    Append $d_i$ to $d$
  **end for**
  Filter: $d \leftarrow \{d_i : d_i < 0\}$ // *Keep only negative distances*
  **if** $d \neq []$ **then**
    $d^* \leftarrow \max\{|d_i| : d_i \in d\}$ // *Maximum absolute negative distance*
    $\gamma^* \leftarrow \gamma$[illegible]

#### S0.8 Metric Quality Quantification

We evaluated the predictive utility and stability of validation metrics using the following statistical measures.

##### S0.8.1 Predictive Utility

**Multiple Linear Regression** To assess the independent predictive utility of $\text{MSE}_{\text{Training}}$ and Apparent $\text{Work}_{\text{HDXer}}$ for ground-truth Open State Recovery%, we fitted a standardised multiple linear regression model:

$$\text{Recovery} = \theta_1 \text{MSE}_{\text{Training}} + \theta_2 \text{Apparent Work}_{\text{HDXer}} + \epsilon \tag{S1}$$

**Where:**

- $\theta_1$ and $\theta_2$ are standardised regression coefficients
- $\epsilon$ is the error term

**Interpretation:** The magnitude of $\theta$ indicates the strength of the relationship between the predictor and the recovery rate, while the sign indicates the direction of the effect.

**Partial $R^2$** To assess the unique contribution of each predictor, we calculated partial $R^2$ values, defined as the reduction in model $R^2$ when excluding that predictor:

$$\text{Partial } R_i^2 = R_{\text{full}}^2 - R_{\text{without},i}^2 \tag{S2}$$

**Where:**

- $R_{\text{full}}^2$ is the coefficient of determination for the full model
- $R_{\text{without},i}^2$ is the coefficient of determination for the model excluding predictor $i$

**Interpretation:** Higher partial $R^2$ indicates greater independent predictive value.

##### S0.8.2 Stability Across Data Splits

To quantify metric stability across different peptide selections, we employed variance decomposition to partition total variance into between-split and within-split components.

**Variance Components** Between-split variance captures systematic differences attributable to split-type selection:

$$\sigma^2_{\text{between}} = \frac{\sum_{j=1}^{k} n_j(\bar{x}_j - \bar{x}_{\text{grand}})^2}{k-1} \tag{S3}$$

**Where:**

- $k$ is the number of split types
- $n_j$ is the number of observations in split type $j$
- $\bar{x}_j$ is the mean of split type $j$
- $\bar{x}_{\text{grand}}$ is the grand mean across all observations

Within-split variance reflects variability within each split type:

$$\sigma^2_{\text{within}} = \frac{1}{k}\sum_{j=1}^{k} \text{Var}(x_j) \tag{S4}$$

**Stability Index** We defined a stability index as the proportion of total variance arising from within-split variation rather than between-split differences:

$$SI = 1 - \frac{\sigma^2_{\text{between}}}{\sigma^2_{\text{between}} + \sigma^2_{\text{within}}} \tag{S5}$$

**Interpretation:** Values approaching 1 indicate high stability (metric insensitive to split choice), while values near 0 indicate high split-dependence (metric varies substantially with peptide selection).

**Coefficient of Variation** To assess relative variability independent of scale, we computed the coefficient of variation (CV) across split-type means:

$$\text{CV} = \frac{\sigma_{\text{splits}}}{\mu_{\text{splits}}} \tag{S6}$$

**Where:**

- $\sigma_{\text{splits}}$ is the standard deviation of split-type means
- $\mu_{\text{splits}}$ is the mean of split-type means

**Interpretation:** Lower CV values indicate more consistent metric behaviour across different data partitions.

**Effect Size of Split Type** ($\eta^2$) We quantified the effect size of split-type selection using $\eta^2$, representing the proportion of total variance explained by split type:

$$\eta^2 = \frac{SS_{\text{between}}}{SS_{\text{total}}} \tag{S7}$$

**Where:**

- $SS_{\text{between}} = \sum_{j=1}^{k} n_j(\bar{x}_j - \bar{x}_{\text{grand}})^2$
- $SS_{\text{total}} = \sum_{i=1}^{N}(x_i - \bar{x}_{\text{grand}})^2$

### References


1. Virtanen P, Gommers R, Oliphant TE, Haberland M, Reddy T, Cournapeau D, et al. SciPy 1.0: Fundamental Algorithms for Scientific Computing in Python. Nat Methods 2020; **17**:261–72. doi:10.1038/s41592-019-0686-2
2. Richards A. University of Oxford Advanced Research Computing. 2015. doi:10.5281/zenodo.22558
3. Mirdita M, Schütze K, Moriwaki Y, Heo L, Ovchinnikov S, Steinegger M. ColabFold: making protein folding accessible to all. Nat Methods 2022; **19**:679–82. doi:10.1038/s41592-022-01488-1
4. Jumper J, Evans R, Pritzel A, Green T, Figurnov M, Ronneberger O, et al. Highly accurate protein structure prediction with AlphaFold. Nature 2021; **596**:583–9. doi:10.1038/s41586-021-03819-2
